# Quantification of Microtubule Stutters: Dynamic Instability Behaviors that are Strongly Associated with Catastrophe

**DOI:** 10.1101/2019.12.16.878603

**Authors:** Shant M. Mahserejian, Jared P. Scripture, Ava J. Mauro, Elizabeth J. Lawrence, Erin M. Jonasson, Kristopher S. Murray, Jun Li, Melissa Gardner, Mark Alber, Marija Zanic, Holly V. Goodson

**Affiliations:** Department of Chemistry and Biochemistry, University of Notre Dame, IN 46556; Department of Applied and Computational Mathematics and Statistics, University of Notre Dame, IN 46556; Pacific Northwest National Laboratory, Richland, WA 99352; Department of Mathematics, University of California Riverside, Riverside, CA 92521; Department of Cell and Developmental Biology, Vanderbilt University, Nashville, TN 37240; Department of Natural Sciences, Saint Martin’s University, Lacey, WA 98503; Department of Chemical and Biomolecular Engineering, Vanderbilt University, Nashville, TN 37235; Department of Biochemistry, Vanderbilt University, Nashville, TN 37205; Department of Genetics, Cell Biology, and Development, University of Minnesota, Minneapolis, MN 55455; Department of Mathematics and Statistics, University of Massachusetts Amherst, Amherst MA, 01003

## Abstract

Microtubules (MTs) are cytoskeletal fibers that undergo dynamic instability (DI), a remarkable process involving phases of growth and shortening separated by stochastic transitions called catastrophe and rescue. Dissecting dynamic instability mechanism(s) requires first characterizing and quantifying these dynamics, a subjective process that often ignores complexity in MT behavior. We present a <u>S</u>tatistical <u>T</u>ool for <u>A</u>utomated <u>D</u>ynamic <u>I</u>nstability <u>A</u>nalysis (STADIA), which identifies and quantifies not only growth and shortening, but also a category of intermediate behaviors that we term ‘stutters.’ During stutters, the rate of MT length change tends to be smaller in magnitude than during typical growth or shortening phases. Quantifying stutters and other behaviors with STADIA demonstrates that stutters precede most catastrophes in our dimer-scale MT simulations and *in vitro* experiments, suggesting that stutters are mechanistically involved in catastrophes. Related to this idea, we show that the anti-catastrophe factor CLASP2γ works by promoting the return of stuttering MTs to growth. STADIA enables more comprehensive and data-driven analysis of MT dynamics compared to previous methods. The treatment of stutters as distinct and quantifiable DI behaviors provides new opportunities for analyzing mechanisms of MT dynamics and their regulation by binding proteins.

## INTRODUCTION

Microtubules (MTs) are protein-based biological polymers that have a central role in fundamental eukaryotic processes including cellular organization, chromosome separation during cell division, and intracellular transport (Goodson and Jonasson 2018). Crucial to the function of MTs in these processes is a well-known behavior termed dynamic instability (DI), where the polymers switch stochastically between periods of growth and shortening as seen in traditional MT length-history plots (**Figure 1 A,B**) (Mitchison and Kirschner 1984; Desai and Mitchison 1997). Accurate quantification of MT DI behavior is essential for understanding its significance and mechanism and also for investigating the activities of DI-regulating proteins and pharmaceutical agents (e.g., chemotherapy drugs, fungicides).

**Figure 1.**
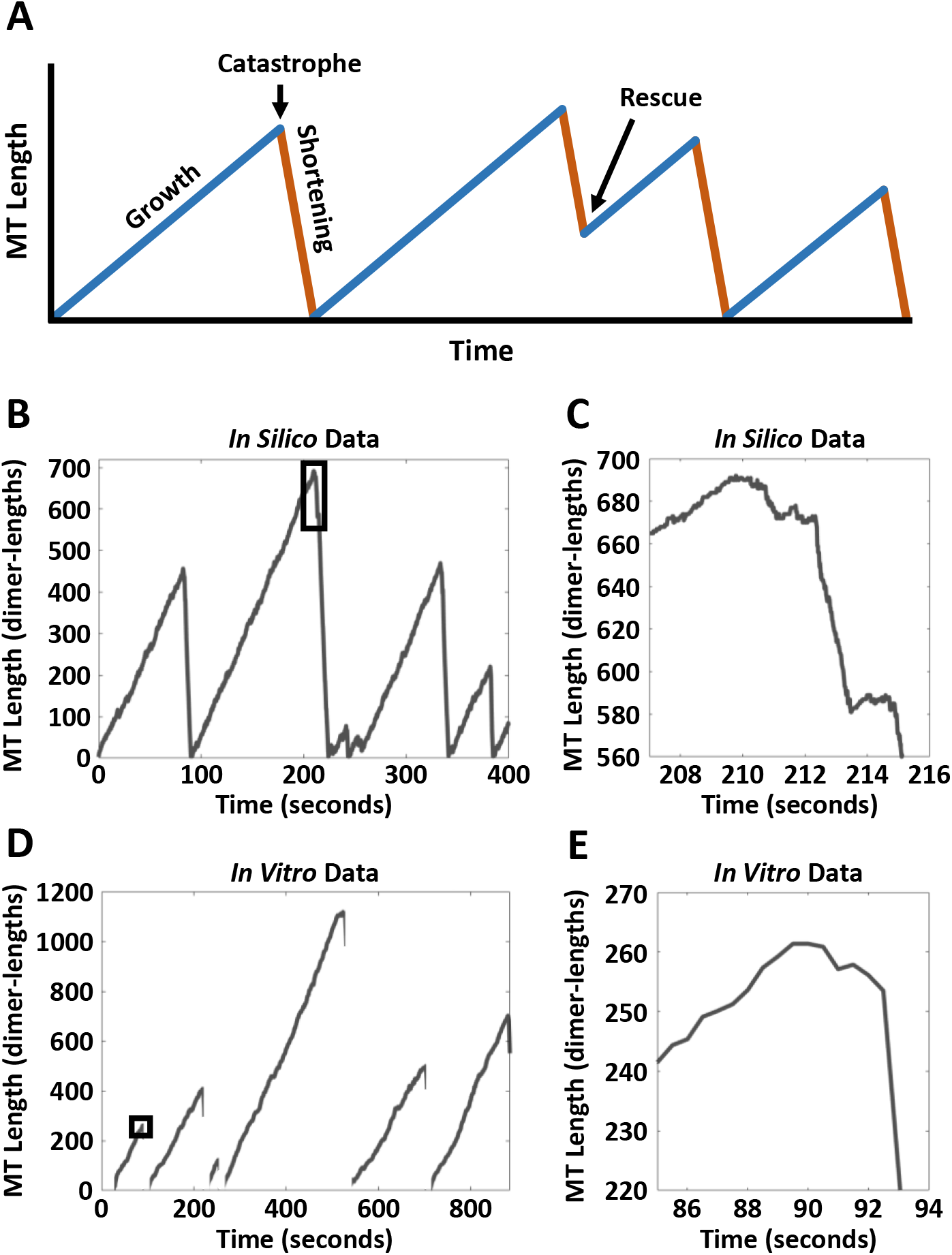
Qualitative examples of behavior that does not fit two-state (growth-shortening) framework in high-resolution simulation (*in silico*) and experimental (*in vitro* control) data. **(A)** An illustration of the classically recognized two-state representation of dynamic instability (DI), in which behavior is recognized as either growth or shortening phases, with instantaneous transitions known as catastrophe and rescue events. **(B,D)** Zoomed-out length-history plots of simulation data (B, detailed 13-protofilament model, see Methods) and experimental data (D, note that depolymerizations were not tracked in their entirety in these experiments). Black rectangles in B and D indicate the zoomed-in portions shown in C and E, respectively. **(C,E)** Closer inspection of transitions shows ambiguous behavior that cannot clearly be categorized as either growth or shortening. Other behaviors need to be considered in order to fully describe MT transitions and identify the exact moments where transitions occur.

### Limitations of common methods for quantifying dynamic instability

Traditionally, MTs have been treated as two-state polymers; that is, MTs have been considered to be either growing or shortening, with abrupt, instantaneous transitions called catastrophes and rescues between these two phases (**Figure 1 A,B,D**). In this framework, MT behavior is characterized by four quantities called DI parameters (Walker et al. 1988):

- **V_growth_** - velocity of growth, commonly measured as the mean of the growth rates over the set of growth phases
- **V_short_** - velocity of shortening, commonly measured as the mean of the shortening rates over the set of shortening phases
- **F_cat_** - frequency of catastrophe, commonly measured as the number of catastrophes (transitions from growth to shortening) per time in growth
- **F_res_** - frequency of rescue, commonly measured as the number of rescues (transitions from shortening to growth) per time in shortening

In this approach to quantifying MT dynamics, the velocity of an individual growth or shortening phase is typically determined as the slope of a line drawn between the points of catastrophe and rescue (e.g., (Zanic 2016)).

While determination of DI parameters as described above is now a standard way to quantify MT behavior (e.g., (Portran et al. 2017; Zwetsloot, Tut, and Straube 2018; Kapoor et al. 2019)), there are aspects of MT behavior that are not captured using this approach. First, it has long been recognized that both growth and shortening rates are variable. This variability occurs both with and without MT binding proteins (MTBPs), and it is observed both within and between individual growth phases and similarly for shortening phases (e.g., (Gildersleeve et al. 1992; Pedigo and Williams 2002; Schek et al. 2007; Lawrence et al. 2018)). In addition, analysis of variability in growth and shortening rates suggested that the two-state (growth and shortening) model approximation agreed well with experimentally observed MT behavior at time scales greater than ~ 1 minute, but underestimated the observed variability at time scales less than ~ 1 minute (David J. Odde, Buettner, and Cassimeris 1996). These observations raise the concern that DI analysis methods that categorize an entire period between nucleation (or rescue) and catastrophe as a single growth or shortening phase could cause finer, but functionally significant, aspects of MT behavior to be missed.

Second, pauses, attenuation phases, and other intermediate states have been observed in experiments and proposed in models, but the way these behaviors have been identified and defined has varied. Pauses are commonly observed *in vivo* (e.g., (Sammak and Borisy 1988; Schulze and Kirschner 1988; Shelden and Wadsworth 1993; Dhamodharan and Wadsworth 1995; Waterman-Storer and Salmon 1997; Gierke, Kumar, and Wittmann 2010; Applegate et al. 2011)). Pauses have also been observed *in vitro* in the presence of MTBPs (e.g., (Moriwaki and Goshima 2016)), cell extracts (e.g. (Keller et al. 2008)), drugs (e.g., (Toso et al. 1993)), and occasionally for purified tubulin (e.g., (Walker et al. 1988)). Identification of pause phases in length-history data frequently relies on arbitrary fixed thresholds. A length-change threshold of 0.5 microns has been common, but the exact way in which this threshold was applied to data has varied among research groups (Sammak and Borisy 1988; Dhamodharan and Wadsworth 1995; Dhamodharan et al. 1995; Rusan et al. 2001; Kamath, Oroudjev, and Jordan 2010; Fees, Estrem, and Moore 2017). Others have used combinations of thresholds on the speed of length change (e.g., in pixels per frame, or microns per minute), length change itself, and/or number of data points involved (Panda et al. 1996; Yenjerla, Lopus, and Wilson 2010; Matov et al. 2010; Gierke, Kumar, and Wittmann 2010; Kiris, Ventimiglia, and Feinstein 2010; Mahrooghy et al. 2015; Moriwaki and Goshima 2016). Recognition of states other than growth and shortening has also led various authors to consider theoretical three- or four-state models in which the additional states are pauses or an intermediate state (D. J. Odde, Cassimeris, and Buettner 1995; David J. Odde, Buettner, and Cassimeris 1996; Tran, Walker, and Salmon 1997; Jánosi, Chréien, and Flyvbjerg 2002; Maly 2002; Keller et al. 2008; Smal et al. 2010; Blackwell et al. 2017). Thus, it is clear that many researchers are interested in methods for quantifying states beyond growth and shortening in data and the inclusion of such states in the development of theory, but it is less clear how these states should be defined.

An important related point is that recent improvements in imaging technology have enabled acquisition of MT dynamics data with both high temporal and high spatial resolution, which allows for the possibility of analyzing length-history data at finer scales. These data have verified the intrinsic variability of MT behavior. They have also demonstrated that both growth and shortening phases can include significant time periods (e.g., a few seconds in duration or longer) during which the growth or shortening velocity slows significantly (**Figure 1 C,E**; see also (Maurer et al. 2014; Duellberg, Cade, and Surrey 2016; Duellberg et al. 2016; Rickman et al. 2017)). These slowdown periods likely overlap with pauses discussed above, though it is important to note that “*bona fide*” pauses are often considered to be time periods “during which no polymerization or depolymerization occurs” (Gierke, Kumar, and Wittmann 2010), and so are separable from periods of slowed growth or shortening, at least in theory.

Significantly, these slowdown periods can also occur in association with catastrophe ((Maurer et al. 2014; Duellberg, Cade, and Surrey 2016; Duellberg et al. 2016), see also predictions based on simulations in (Margolin et al. 2012)), making it difficult to determine with reasonable precision where transitions between phases begin and end. To illustrate this problem, consider the zoomed-out length-history plots that are typically used for DI analysis (**Figure 1 B,D**). Examination of these plots can make the task of determining when transitions occur look trivial. However, the zoomed-in views made possible by high-resolution data acquisition (**Figure 1 C,E**) demonstrate the difficulty of identifying the points of transition and/or categorizing DI behaviors.

Thus, many researchers have recognized that MT dynamic instability behavior is more complex than a simple two-state system of growth and shortening with abrupt transitions. The four traditional DI parameters (V_growth_, V_short_, F_cat_, and F_res_) would be sufficient to quantify such a two-state system but are not sufficient to quantify all aspects of observed MT dynamic instability. Previous articles have quantified some aspects, particularly time durations, of the slowdown periods noted above, but have not quantified velocities and transition frequencies in a way that would provide an alternative to the traditional four DI parameters. One previous approach has been to exclude slowdown periods from quantification of DI parameters, since including them in either growth or shortening phases would reduce the magnitude of measured values of V_growth_ and V_short_ (e.g., (Rickman et al. 2017)). However, there are two problems with this approach. First, it leaves researchers to make subjective judgments or use ‘in-house’ software to identify both the periods to exclude and the points where phase transitions occur (e.g., (Yenjerla, Lopus, and Wilson 2010; Zanic 2016; Jonasson et al. 2020)). This approach impacts precision and reproducibility. Second, and perhaps more significantly, entirely excluding these behaviors from analysis could potentially result in the loss of information critical for understanding the mechanisms of the phase transitions or their regulation by MT binding proteins. Thus, capturing and quantifying these alternative behaviors is a key step towards further dissecting the recognized variations in growth and shortening rates, improving the precision of DI metrics, and elucidating mechanisms of dynamic instability.

### Summary of conclusions

Using established statistical methods, we developed the <u>S</u>tatistical <u>T</u>ool for <u>A</u>utomated <u>D</u>ynamic <u>I</u>nstability <u>A</u>nalysis (STADIA), an automated tool for characterizing and quantifying MT behavior. Applying STADIA to *in silico* and *in vitro* MT length-history data demonstrated the prevalence of a category of intermediate behaviors that we propose calling ‘stutters’. Stutters share similar characteristics with each other and are distinguishable from typical growth and shortening. The primary distinguishing factor is that during stutters the overall rate of change in MT length is markedly less in magnitude compared to the velocities of classically recognized growth and shortening phases. Stutters are also distinguishable from pauses in that during true pauses “no polymerization or depolymerization occurs” (Gierke, Kumar, and Wittmann 2010). In contrast, during stutters dimer-scale dynamics continue, and during most stutters measurable length changes do occur although at slower velocities than during typical growth and shortening. Stutters, as recognized and quantified by STADIA, overlap with previously observed behaviors such as pre-catastrophe slowdowns (e.g., (Maurer et al. 2014)) and events that have been called “pauses” despite length changes occurring (e.g., (Matov et al. 2010; Kamath, Oroudjev, and Jordan 2010; Guo et al. 2018)).

Analysis of length-history data using STADIA leads to two major observations regarding the relationship between stutters and catastrophes:

- Stutters precede most catastrophes observed in our *in silico* and *in vitro* (with purified proteins) datasets.
- The MT stabilizing protein CLASP2γ reduces catastrophe by increasing the fraction of stutters that return to growth rather than enter shortening phases. Specifically, CLASP2γ reduces the frequency of growth-to-stutter-to-shortening (which we term transitional catastrophe) and increases the frequency of growth-to-stutter-to-growth (which we term interrupted growth).

These results indicate that STADIA is able to recognize and quantify behaviors that may be missed by classical methods of analyzing MT length-history data. Furthermore, these results suggest that stutters play a mechanistically significant role in the process of catastrophe.

Different from standard DI analysis methods, STADIA combines the following features:

- STADIA uses a data-driven approach to distinguish different types of behaviors, as opposed to using predefined fixed thresholds. While stutters tend to have slower velocities than typical growth and shortening, the specific numerical values will differ in different systems, and “slower” is relative to the growth and shortening velocities in each system. STADIA’s data-driven approach allows researchers to find distinguishing values that are relevant for their particular system.
- STADIA expands on the traditional four DI parameters by measuring the frequencies of all possible transitions among growth, shortening, and stutters, and measuring the velocities of each type of segment. Similar transition analysis was applied to growth, shortening, and pause phases in cell extracts (Keller et al. 2008), but has not yet been applied to the more recently observed dynamic slowdowns discussed above (e.g., (Maurer et al. 2014)).
- STADIA comprehensively identifies stutters that occur anywhere in length-history data, in contrast to previous articles that focused on only pre-catastrophe slowdowns (Maurer et al. 2014; Duellberg, Cade, and Surrey 2016; Duellberg et al. 2016) or episodes of slow growth within larger growth phases (Rickman et al. 2017). Without a method that identifies both of these types of events, it would not have been possible to make the observation here that CLASP2γ reduces the fraction of growth-to-stutters events that become transitional catastrophes (i.e., pre-catastrophe slowdowns) and increases the fraction that become interrupted growths.

The relationship of our results to previous work is further covered in the Discussion section. We conclude that identification of stutters as distinct from growth, shortening, or pause warrants their future inclusion in DI analyses, and serves as a necessary step forward in gaining a better understanding of MTs, their dynamics, and their regulation by MT binding proteins.

## RESULTS

We first present a brief overview of STADIA and its analysis procedure (readers are encouraged to refer to the Methods for more detailed information; see also (Patel et al. 2020)). We then use STADIA to analyze MT dynamics as they are observed in simulations (*in silico*) and in experiments (*in vitro*), including detection and quantification of a category of intermediate behaviors that we term ‘stutters’. We use this analysis as a foundation on which to study the relationship between stutters and phase transitions, showing that stutters are strongly associated with catastrophe. We further test the functional significance of stutters in catastrophe and demonstrate the utility of STADIA in studying MT-binding proteins by using STADIA to analyze the dynamics of *in vitro* MTs growing in the presence of the anti-catastrophe factor CLASP2γ, thus examining for the first time its effect on stutter.

### STADIA: A Novel Tool for Analyzing Dynamic Instability Behavior of MTs

To meet the goal of more precisely identifying, categorizing, and quantifying the range of MT behaviors, we created the <u>S</u>tatistical <u>T</u>ool for <u>A</u>utomated <u>D</u>ynamic <u>I</u>nstability <u>A</u>nalysis (STADIA). Specific aims for the development of STADIA were that it be: **1)** Automated to create a consistent and reproducible method with minimal user input; **2)** Impartial such that it does not presuppose MT dynamics are restricted to two states (i.e., limited to growth and shortening); **3)** Adaptive to handle varying time durations of phases and the stochastic nature of phase changes; **4)** Compatible with classical DI analysis, enabling comparison to and continuity with previous work; **5)** Capable of analyzing MT time-series data sourced from both computational simulations and laboratory experiments.

The resulting software, STADIA, is a data-driven tool that uses time-series analysis to characterize and quantify MT behavior. STADIA can be run in two modes (described in more detail in the Methods section): Diagnostic Mode (useful for performing preliminary analyses and tuning analysis parameters), and Automated Mode (outlined below, used for performing full DI analysis). The process, presently implemented in MATLAB, has three major stages (**Figure 2 C-I, Supplemental Figure S1.1**):

**Figure 2.**
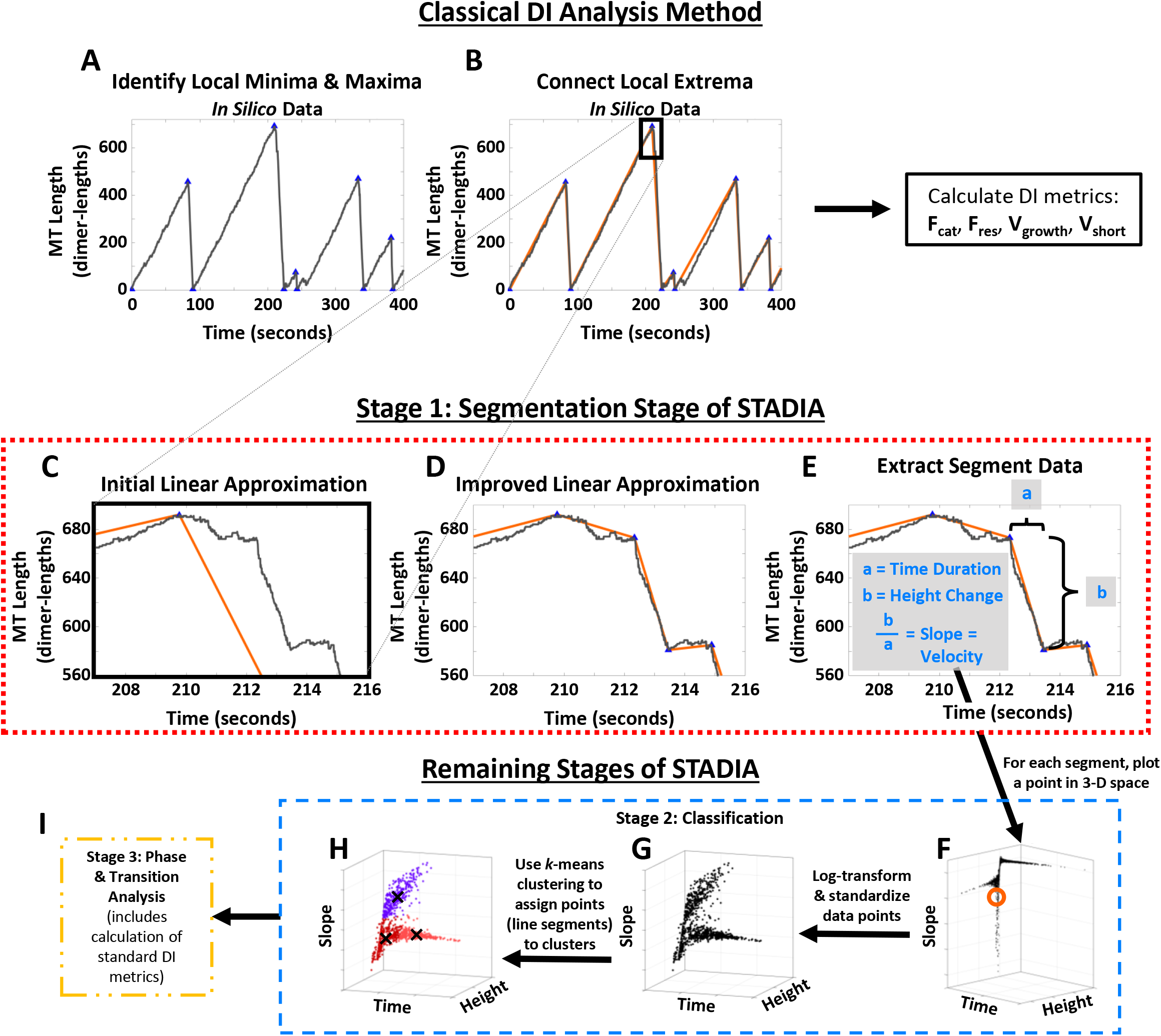
Comparison of classical DI analysis method to STADIA analysis method. **(A,B)** <u>Classical DI analysis method</u>: Identify major peaks and valleys (A, blue triangles), and define these points as catastrophes and rescues (or nucleation events), respectively. Define each segment from a nucleation event or rescue to a catastrophe as a growth phase. Define each segment from a catastrophe to a nucleation event or rescue as a shortening phase. Calculate F_cat_ and F_res_ from the number of catastrophes or rescues divided by the total time in growth or shortening, respectively. Calculate V_growth_ and V_short_ from the slopes of straight lines connecting the start and end points of each segment (B, orange lines) or from regression lines fitted to the data points in each segment. **(C-I)** <u>STADIA analysis method</u>. **C-E:** <u>Segmentation Stage of STADIA</u>: First, generate an initial approximation by finding major peaks and valleys (local extrema with high prominence) and connecting these with line segments (C). Next, iteratively include new vertices where the largest error occurs in each segment to produce the final approximation (D), such that all pointwise errors between the raw length-history data and the final approximation are less than a user-defined threshold (usually 10 to 25 dimer-lengths). Note that using the process in (C) (analogous to the classical segmentation procedure) followed by the process in (D) produces a closer approximation of the data than the process in (C) alone (compare panels D and C). Lastly, calculate time duration, height change, and slope (velocity) for each segment (E). **F-I:** <u>Brief Overview of Remaining Stages of STADIA</u>: After the Segmentation Stage of STADIA, the resulting segment data are plotted in 3-D space (the orange circle in (F) denotes the approximate location of the data point corresponding the example line segment in (E)). In the Classification Stage of STADIA, the line segment data are separated into positive and negative slopes (note that only negative slope segment data are shown in (G,H)), then log-transformed and standardized (G), and used for *k*-means clustering (H). The average segment features of each cluster are then used to assign clusters to named DI behaviors. After the Classification Stage, the Phase and Transition Analysis Stage of STADIA calculates metrics including (and expanding beyond) the traditional V_growth_, V_short_, F_cat_, and F_res_ (I). See the Methods section and **Supplemental Figure S1.1** for additional information.

1. <u>Segmentation</u>: STADIA creates a continuous piecewise linear approximation of MT length-history data, where segment endpoints mark moments of significant change in behavior, i.e., transitions between periods of different sustained slopes in the length-history plots (**Figure 2 C,D,E**). The level of accuracy of the approximation is regulated by user-defined parameters.
2. <u>Classification</u>: STADIA groups the individual segments from the linear approximation in Stage 1 using an established clustering method called *k*-means (**Figures 2 F-H, 3 A,B,D,E, Supplemental Figures S1.4,S1.5**). STADIA then bundles segment clusters with similar characteristics into DI phase/behavior classes (**Figure 3 C,F,I, Supplemental Figure S1.8**). Before running the Automated Mode, the number of clusters for *k*-means to identify is informed by running STADIA in the Diagnostic Mode.
3. <u>Phase and Transition Analysis</u>: STADIA then applies the segment classifications from Stages 1 and 2 to the length-history plots (**Figure 3 G,H, Figure 6**) and considers chronological ordering to perform quantitative phase and transition analysis (this includes measuring the frequencies of catastrophe, rescue, and other possible transitions) (**Figure 2 I, Figures 4-7, Table 1, Supplemental Figures S1.9, S1.10**).

**Table 1.**
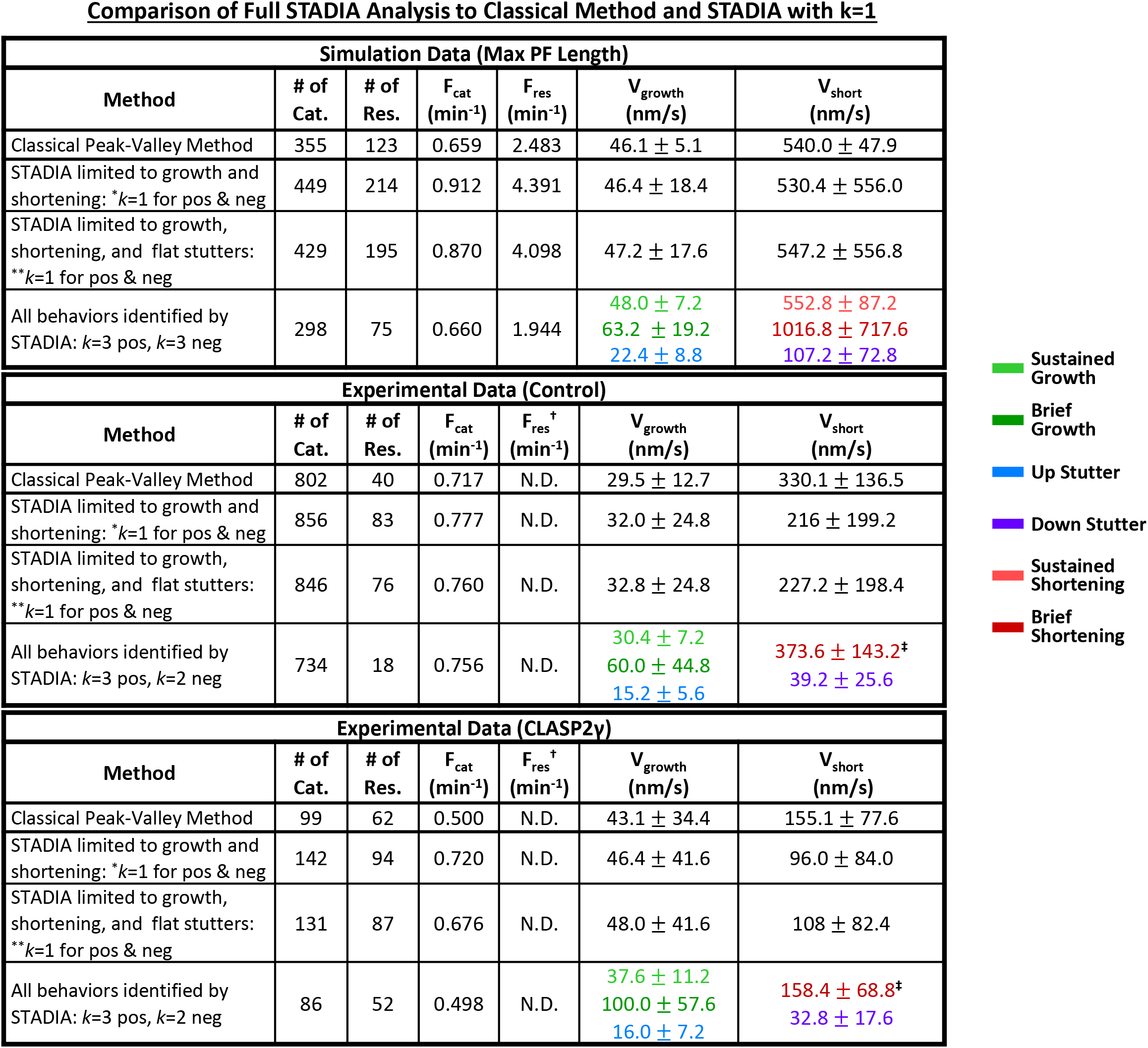
Comparison of DI metrics from classical two-state analysis, STADIA two-state analysis (i.e., STADIA with *k* =1), and STADIA full analysis. The results from the full, automated analysis conducted by STADIA (bottom row of each table) were compared to results from both a classical method (top row of each table; performed by identifying only major peaks and valleys, to form a course-grained approximation) and analyses where STADIA classification was limited to two states (middle rows of each table; here a fine-grained approximation was generated by STADIA, but phase classes were restricted to only growth and shortening (*) or growth, shortening, and flat stutters (**)). There is general, but not exact, agreement between methods. V_growth_ and V_short_ measurements are listed as mean +/− standard deviation over the set of all segments identified in each type of behavior. See **Supplemental Figure S1.9** for the number of segments in each cluster from the STADIA full analysis. See the Methods section for the number of MTs and total observation times in each dataset. †,‡: Because depolymerizations in the experimental datasets were not captured in their entirety, rescue frequency was not reported (†), and negative slope segments were separated into only two clusters, yielding only two V_short_ measurements in the full STADIA analysis (‡), instead of three as seen with the *in silico* data.

**Figure 3.**
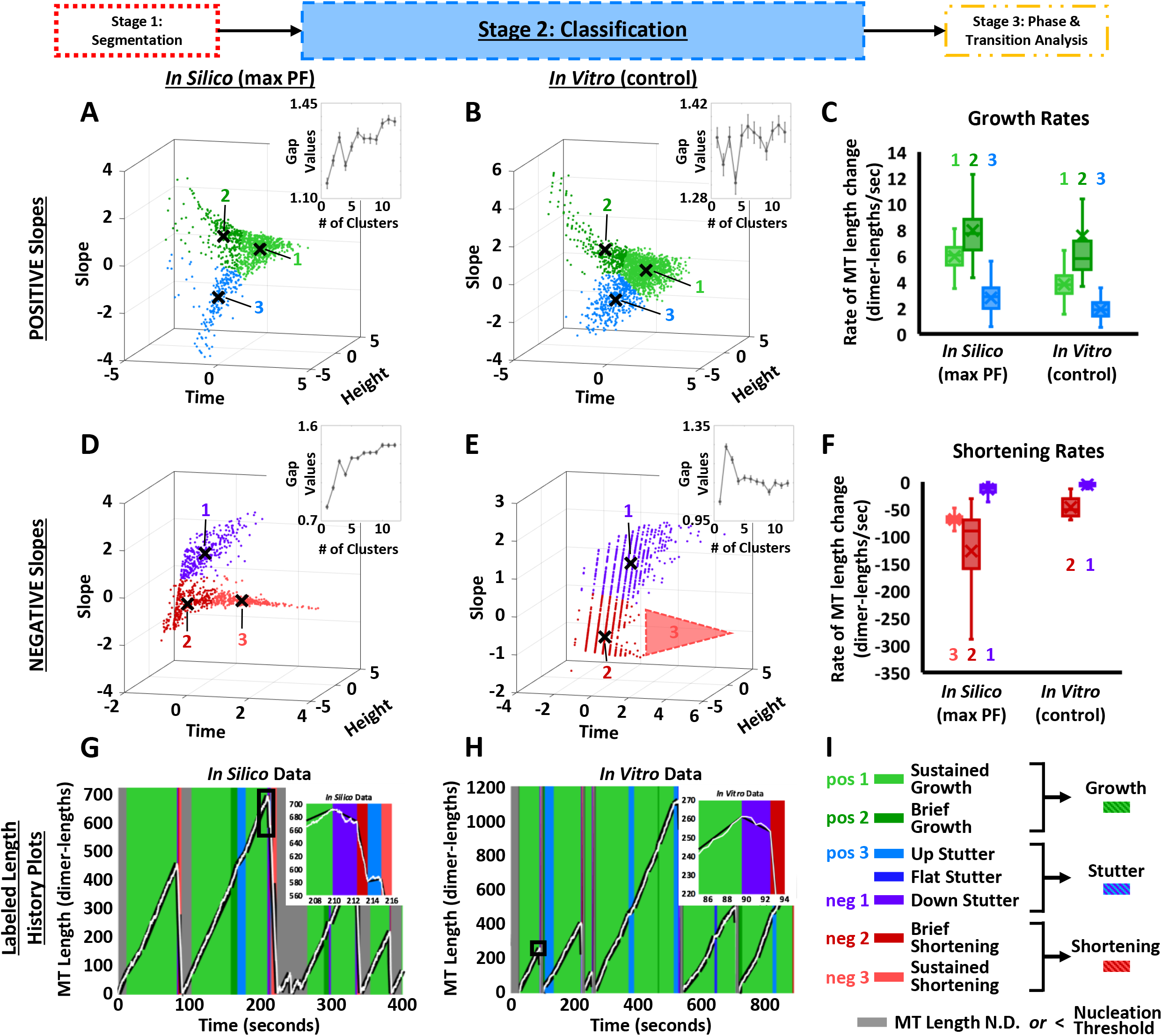
Classification Stage of STADIA. As illustrated in **Figure 2**, STADIA converts length-history data to segments, and then for each segment plots a point in 3-D space that uniquely corresponds to that segment’s time duration, height change, and slope. Using *k*-means clustering (an established machine-learning algorithm), STADIA then divides the points (and thus the segments) into *k* clusters (groups), which are then categorized into DI behavior classes. The main text and supplementary data provide justification for the conclusion that *k*=3, i.e., that there are three clusters each of positive and negative slope segments (*k*=2 for the *in vitro* negative slope segments, because complete depolymerizations to the seeds were not captured). **(A,B,D,E)** Clustering results for the log-transformed and standardized segment features of the positive (A,B) and negative (D,E) slope segments extracted from *in silico* (A,D) and *in vitro* (B,E; control, no CLASP2γ) length-history data approximations. **(C,F)** Box plots of growth rates (segment slopes) for the positive slope segment clusters (C) and shortening rates (segment slopes) for negative slope segment clusters (F) identified in the *in silico* and *in vitro* datasets. For the positive slope segments in each dataset, clusters 1 and 2 (light and dark green) have average growth rates relatively large in magnitude compared to cluster 3 (light blue). For the negative slope segments in each dataset, cluster 2 (dark red) and cluster 3 (light red, *in silico* data only) have average shortening rates relatively large in magnitude compared to cluster 1 (purple). Outliers were excluded from these plots using the default definition in MATLAB (i.e., any value that is more than 1.5 times the interquartile range away from the bottom or top of the box is considered an outlier); outliers are *not* excluded from the cluster plots in (A,B,D,E). **(G,H)** Previously unlabeled *in silico* and *in vitro* MT length-history plots (see **Figure 1 B,D**) are now labeled according to the cluster that each line segment fits into. Zoomed-in portions of previously ambiguous length-history data (see **Figure 1 C,E**) are now clearly labeled. **(I)** Examination of the average slopes of the individual clusters indicates that bundling clusters together into larger behavior classes based on the average slopes of the individual clusters is appropriate (see also **Supplemental Figure S1.8**). Clusters with positive and negative slopes relatively larger in magnitude were bundled together into ‘Growth’ and ‘Shortening’, respectively. The remaining clusters, with shallower slopes (i.e., slower rates of length change), were bundled together, along with the previously identified ‘near-zero’ slope or flat segments, into ‘Stutters’. Further, ‘Brief’ and ‘Sustained’ clusters of the Growth and Shortening classes are distinguished by their time durations. The ‘Up’, ‘Flat’, and ‘Down’ clusters of Stutter are characterized by their segment slopes being positive, near-zero, or negative, respectively.

**Figure 4.**
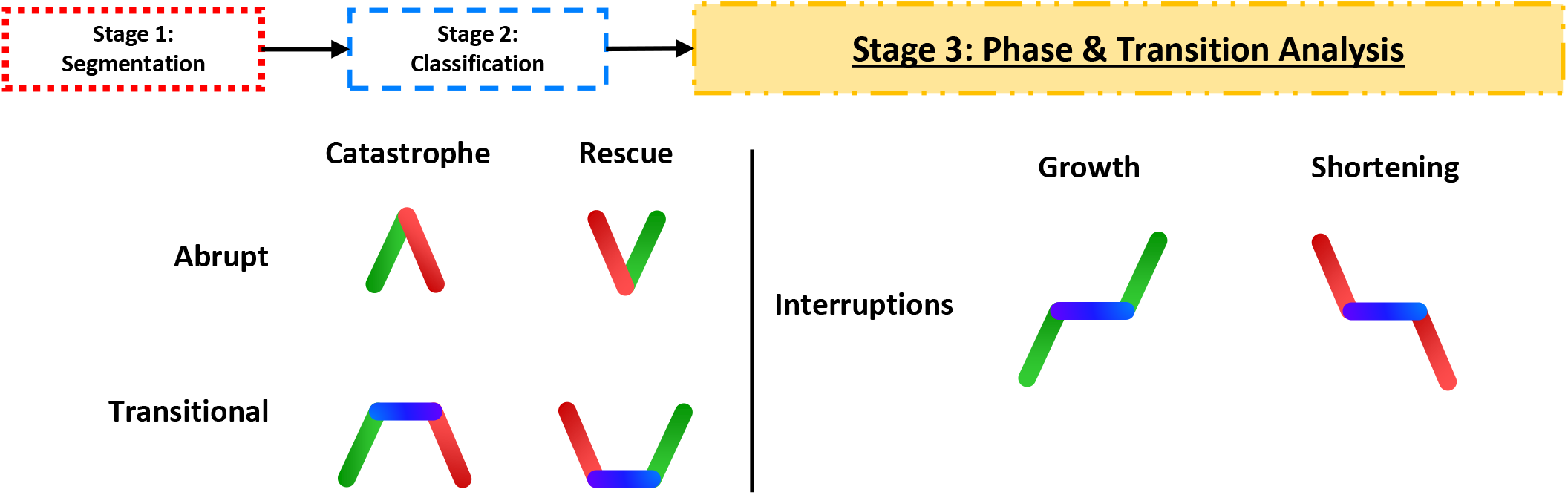
Schematic representations of transitions identifiable by STADIA. Abrupt catastrophe: growth-shortening (i.e., growth followed directly by shortening). Abrupt rescue: shortening-growth. Transitional catastrophe: growth-stutter-shortening. Transitional rescue: shortening-stutter-growth. Interrupted growth: growth-stutter-growth. Interrupted shortening: shortening-stutter-shortening.

Note that the segmentation stage occurs before the classification stage and does not rely on any assumptions regarding how segments will end up being classified. The points of transition between segment endpoints in the linear approximation include extreme events such as traditionally recognized catastrophes and rescues as well as more subtle changes in the rate of MT length change (**Figure 2 A-D**). More information about each stage of STADIA is provided in the Methods section and **Supplemental Figure S1.1**. Limitations of STADIA and guidance for users are also discussed in the Methods section. In the analysis below, we used STADIA to analyze length-history data sourced from both simulations and experiments (*in silico* and *in vitro*). The simulation dataset was obtained using our detailed dimer-scale 13-protofilament kinetic Monte Carlo model of MT dynamics (Margolin et al. 2012). A subset of the experimental dataset used here was previously analyzed using other methods in (Lawrence et al. 2018). Please see the Data Acquisition subsections of the Methods for additional information about the simulations and the physical experiments.

### Constraining STADIA to perform two-state analysis, for comparison with classical DI measurements

In initial testing, we used STADIA to analyze the *in silico* and *in vitro* datasets under settings where we forced the classification stage of STADIA to assume that MT dynamics consist of only two states: growth and shortening phases (‘STADIA limited to growth and shortening’ analysis in **Table 1**). That is, all positive slope segments were classified as growth and all negative slope segments were classified as shortening. As might be expected, the DI parameters measured by STADIA under these constrained conditions are similar, but not identical, to those measured through a more traditional DI analysis method (‘Classical Peak-Valley Method’ in **Table 1**, described in the Methods section).

Both of the methods used here begin by identifying major peaks and valleys (which correspond to catastrophes and rescues) in the length-history data such that the length change between a peak and valley exceeds a user-defined threshold (**Figure 2 A,B**). While the segmentation process in the more traditional peak-valley method stops at this point, STADIA seeks to make further improvements by iteratively identifying additional vertices (segment endpoints, **Figure 2 D**) until the difference between the piecewise linear approximation and the raw length-history data is less than the user-defined Maximum Error Tolerance. Effectively, as shown in **Figure 2**, the piecewise linear approximation produced by the segmentation stage of STADIA resembles the raw data more closely than does the approximation from classical methods.

Consequently, there are two fundamental differences between the segmentations resulting from classical methods (e.g., as implemented in the peak-valley method) and Stage 1 of STADIA. First, an individual segment of growth or shortening as identified by classical methods (**Figure 2 A,B**) may be composed of multiple segments of varying slopes in the STADIA analysis (**Figure 2 D**). Second, the refined approximation resulting from STADIA identifies segments of shallower slope that are not separated out from longer growth and shortening segments in classical methods.

Thus, STADIA can produce measurements of the traditional DI parameters (i.e., V_growth_, V_short_, F_cat_, and F_res_), but does so by using a finer linear approximation of the length-history data resulting in differences in the measured values of the DI parameters (**Table 1**). However, behaviors such as those captured in **Figure 1C** and **1E** indicate that the traditional DI parameters alone are inadequate to capture the full range of MT dynamics as revealed by data acquired at high temporal resolution, indicating the possible need for more complex considerations when analyzing DI behaviors. These observations provided a solid foundation on which to proceed with using STADIA to analyze DI without preconceptions about how many types of phases (i.e., sustained and distinguishable behaviors) exist in MT length-history data.

### Evidence for multiple types of behavior within the groups of positive and negative slope segments

After the segmentation process, the next task is to classify the segments. To begin the classification process in STADIA, we rely on an unsupervised machine learning algorithm, *k*-means clustering (Lloyd 1982; Macqueen 1967). *K*-means groups together data points that share similar characteristics, i.e., data points that are near each other in a relevant feature space (in our case, the 3-D space of segment time duration, height change, and slope (**Figure 2 F-H**)). The clustering step of STADIA uses *k*-means to assign segments to clusters and does not presuppose that the clusters correspond to any particular DI phase/behavior; after the clustering step is completed (described in this subsection), then STADIA labels the clusters with named DI behaviors based on the average characteristics of the segments in each cluster (described in the two subsections after this one).

The *k*-means algorithm requires that the number of clusters, *k*, be provided in advance (see the Methods for more information regarding *k*-means clustering and its use in this analysis). Though various approaches exist for determining the *k*-value with which to perform the clustering (reviewed by (Steinley 2006; Pham, Dimov, and Nguyen 2005)), STADIA uses a measurement called the gap statistic (Tibshirani, Walther, and Hastie 2001). Analysis of the gap statistic data (generated by running STADIA in Diagnostic Mode; see Methods for more information) provides evidence that MT dynamics can be objectively divided into more behaviors than simply growth and shortening.

More specifically, we performed the gap statistic analysis for each of the three datasets in **Table 1** (*in silico*, *in vitro* control with tubulin + EB1, and *in vitro* with tubulin + EB1 + CLASP2γ) by separately processing positively- and negatively-sloped segments. If MT growth and shortening each corresponded to one behavior (with variation), one would expect to obtain a value of *k*=1 for each of the positive slope group and the negative slope group. However, with the *in silico* dataset, the gap statistic suggested *k*=3 to be the optimal number of clusters for each of these groups (**Figure 3 A,D, Supplemental Figures S1.4,S1.5**). These observations indicate that the *in silico* dataset contains multiple clusters within the positive and negative slope segments. In total, when including the near-zero slope segments (which were separated out before analysis of the positive and negative slope segments), 7 distinct clusters were identified in the simulation data (**Supplemental Figure S1.7**).

Consistent with the *in silico* results, the analysis of the *in vitro* experimental datasets suggested *k*=3 for positive slopes (**Figure 3 B** and **Supplemental Figure S1.4**). However, differences were found between the *in silico* and *in vitro* data in analysis of the negative slopes: the optimal number of clusters was identified to be two (*k*=2) in both experimental datasets (**Figure 3 E** and **Supplemental Figure S1.5**), in contrast to the three clusters (*k*=3) identified in the simulation data. This observation can be explained by the fact that for technical reasons, the *in vitro* datasets contain the beginning of shortening phases, but not full depolymerizations of MTs to near-zero length (present in the simulation dataset). Consistent with this explanation, inspection of the clustering results for the negative slope segments in **Figure 3** shows that segments belonging to Negative Slope Cluster 3 (the cluster with the longest time durations) in the *in silico* data in **Figure 3 D** were not captured for the *in vitro* data in **Figure 3 E**. Therefore, we can only conclude that there are *at least* two clusters with negative slopes for the *in vitro* data. For illustration purposes, a ‘ghost’ region is added to **Figure 3 E** where we expect the missing third negative slope cluster to reside. Thus, including the flat slope segments, we find evidence for at least 6 distinct clusters in the experimental DI data: three clusters of positive slope segments, two clusters of negative slope segments, plus a cluster of near-zero slope segments that were separated from the positive and negative slope segments prior to *k*-means clustering (**Supplemental Figure S1.7**). Note that an additional cluster of negative slope segments might be identified if full depolymerization events were captured in experiments.

In summary, application of *k*-means clustering with gap statistic analysis to DI data from either our simulations or experiments leads to a similar conclusion: the data argue against the idea that MT DI can be characterized as a two-state process consisting of only growth and shortening with instantaneous transitions. More specifically, the results provide evidence for considering multiple types of positive slope behavior (3 clusters) and multiple types of negative slope behavior (3 clusters, or 2 clusters for the truncated experimental data). Note that STADIA’s identification of the cluster boundaries is driven by the dataset itself, in contrast to many existing DI analysis methods that use arbitrary pre-defined thresholds.

After using STADIA in Diagnostic Mode to perform gap statistic analysis and thus gain information about the optimal number of clusters to use in the *k*-means clustering process, we used STADIA in the Automated Mode with the suggested number of clusters (3 positive slope clusters, and 2 or 3 negative slope clusters) to perform a full analysis of MT behavior. In the next two subsections, we examine the differences between these clusters of length-history segments to determine how the segments in these clusters differ from each other and how these clusters might correspond to recognizably different DI behaviors.

### Growth and shortening phases consistent with classical DI analysis are among the multiple types of behavior identified by STADIA

To study the relationships between the clusters of length-history segments and recognizable phases of DI, we examined the average characteristics of the segments in each cluster (**Figure 3** and **Supplemental Figure S1.8**). This analysis showed that, for both the *in silico* and *in vitro* data, some of the clusters correspond to the well-recognized growth and shortening phases of DI. More specifically, two of the positive segment clusters (positive slope clusters 1 and 2 from **Figure 3 A and B**) have slopes (rates of length change) similar to growth rates reported in classical DI analysis (compare STADIA results in **Figure 3 C** and **Table 1** to classical analysis results in **Table 1**). Similarly, negative slope cluster 2 (*in silico* and *in vitro*, **Figure 3 D and E**) and negative slope cluster 3 (*in silico*, **Figure 3 D**) have slopes similar to shortening rates reported in classical DI analysis (compare **Figure 3 F** and **Table 1**). Based on this information, in the classification stage of STADIA, length-history segments were classified as growth if they belonged to one of the clusters with a steep positive slope (Positive Slope Cluster 1 or 2 in **Figure 3 C,I**), and as shortening if they belonged to a cluster with a steep negative slope (Negative Slope Cluster 2 or 3 in **Figure 3 F,I**).

The identification of two clusters of growth segments (and of shortening segments for the *in silico* data) was unexpected. It is notable that in each case (both positive and negative slopes), the clusters differ primarily by duration (brief or sustained, **Figure 3 I** and **Supplemental Figure S1.8**). This observation may be evidence of different behaviors of tapered or split tips (e.g., as observed by (Coombes et al. 2013; Doodhi et al. 2016; Aher et al. 2018)) relative to the rest of the MT; such structures might be able to extend or retract faster than the bulk MT lattice in the absence of lateral bonds. Future work will investigate whether the differences between brief and sustained growth (or shortening) relate to tip structure.

In the next section, we consider the length-history segments that have much shallower slopes, which set them apart from the other growth and shortening behaviors discussed above.

### STADIA detects and quantifies ‘stutters’: a category of dynamic behaviors distinct from growth and shortening

Examination of **Figure 3 A-F** shows that, in addition to clusters with slopes that correspond to rates of length change seen in classical growth or shortening behaviors, STADIA also identifies clusters with much shallower slopes (Positive Slope Cluster 3 and Negative Slope Cluster 1 in **Figure 3 A-F; Table 1**). Moreover, the segments in these shallow-slope clusters have time durations shorter than typical segments classified as sustained growth and sustained shortening, though typically longer than those classified as brief growth and brief shortening segments (**Supplemental Figure S1.8**). We term these shallow-slope clusters of segments ‘stutters’ to convey the idea that these sections of length-history data exhibit high-frequency, low-amplitude fluctuations throughout which the overall rate of MT length change is slow from a macro-level perspective.

Together, these characteristics indicate that these periods of slow length change are clearly distinct from either classical growth or shortening phases. Thus, we assigned the following clusters to a category called ‘stutters’: ‘up stutters’ (Positive Slope Cluster 3), ‘flat stutters’ (Near-zero Slope Cluster) and ‘down stutters’ (Negative Slope Cluster 1) (**Figure 3**).

At this point, every segment of length-history has been classified, and the assignment of individual segments to growth, shortening, and stutter phases can be visualized in the context of the original length-history data as in **Figure 3 G,H**.

Note that most stutters are distinguishable from previously identified ‘pauses’ during which the MT neither grows nor shortens detectably (e.g., (Yenjerla, Lopus, and Wilson 2010; Gierke, Kumar, and Wittmann 2010)). In contrast, MT lengths do indeed change dynamically during most periods identified as stutters (for examples, see **Figure 3 G,H**), with a net rate of change that is small but non-zero (**Figure 3 C,F**). In addition, it is notable that events categorized as pauses are typically described as being rare (<1% of total behavior duration) in the absence of MT stabilizing proteins (e.g., (Walker et al. 1988; Moriwaki and Goshima 2016)). These observations support the conclusion that most stutters are different from events previously classified as pauses, though there is likely some overlap, particularly with the relatively rare flat stutters (**Supplemental Figure S1.9**) and cases where pauses were allowed to be short in duration (e.g., (Walker et al. 1988; Guo et al. 2018)). Stutters as described above likely do encompass the periods of slowed growth or shortening previously noted (but not quantified or characterized in detail) in recent dynamic instability data of *in vitro* MTs acquired at high spatiotemporal resolution (e.g., (Duellberg, Cade, and Surrey 2016; Rickman et al. 2017; Maurer et al. 2014; Duellberg et al. 2016); see also (Margolin et al. 2012)). In contrast to this previous work, here we quantify spontaneously occurring stutters and examine their relationship to other dynamic instability behaviors.

In summary, stutters are a category of intermediate behaviors that share similar characteristics with each other and are distinguishable from typical growth and shortening. Distinguishing the various behaviors described above required the use of segment slope, time duration, and height change, as explained in the Methods section. For any one of these three features individually, there is overlap between different clusters identified in the data (**Figure 3, Supplemental Figures S1.3, S1.8**). Of the three segment features, slope is the primary feature distinguishing stutters from typical growth and shortening (**Supplemental Figure S1.8 A,C**). In other words, the rate of change in MT length tends to be slower during stutters than during growth and shortening. In regard to time durations, up and down stutters respectively have similar or somewhat longer time durations than brief growth/shortening segments, but shorter time durations than sustained growth/shortening segments (**Supplemental Figure S1.8 B,D**).

### MTs spend a significant fraction of time in stutters

We begin to investigate the significance of stutters by first examining the fraction of time that MTs spend in stutters. As one might expect, both *in silico* MTs and physical MTs spend the majority of their time in growth phases. However, in both the simulations and experiments, MTs spend a substantial amount of time in behaviors categorized as stutters. Notably, in our *in silico* datasets, the MTs spent more time in stutters (8%) than in shortening (6%) (**Supplemental Figures S1.9**). The *in vitro* MTs spent a substantial amount of the time in stutters (**Supplemental Figures S1.9**), but direct comparison to time spent in shortening phases is not conclusive because depolymerizations were not fully captured. These observations indicate that stutters contribute appreciably to MT behavior as assessed in length-history plots.

### Negative control: two-state growth-shortening model

As a negative control, we ran STADIA on length-history simulation data from a model designed to have only two states: growth and shortening. As would be expected, STADIA analysis of the length-history data from the two-state model did not identify behaviors comparable to the stutters detected in the dimer-scale simulation data and the *in vitro* data. For a description of the two-state simulations and the analysis results, please see **Supplemental Material Section 4**“Negative Control: Simulations of a Two-State (Growth-Shortening) Model” and **Supplemental Figures S4.1, S4.2, and S4.3**.

### Transition definitions

To investigate the functional significance of stutters, we used STADIA to examine how transitions between phases occur (see **Figure 4** for schematic, **Figure 5 D-I** for *in silico* examples, **Figure 6** for *in vitro* examples with corresponding kymographs, **Figure 7 D-I** for additional *in vitro* examples, and **Supplemental Figure S1.10** for frequencies). Specifically, we wished to quantify all examples of transitions to/from growth and shortening, with or without stutters. Considering the chronological ordering of phases, STADIA automatically categorizes the following variations of these phase transitions:

- ‘Abrupt Catastrophe’: growth → shortening directly (synonymous with classically recognized catastrophe) (**Figures 5 D, 6 A,B, 7 D,E**)
- ‘Abrupt Rescue’: shortening → growth directly (synonymous with classically recognized rescue) (**Figure 5 E**)
- ‘Transitional Catastrophe’: growth → stutter → shortening (**Figures 5 F, 6 C,D, 7 F,G**)
- ‘Transitional Rescue’: shortening → stutter → growth (**Figure 5 G**)
- ‘Interrupted Growth’: growth → stutter → growth (**Figures 5 H, 6, 7 H,I**)
- ‘Interrupted Shortening’: shortening → growth → shortening (**Figure 5 I**)

**Figure 5.**
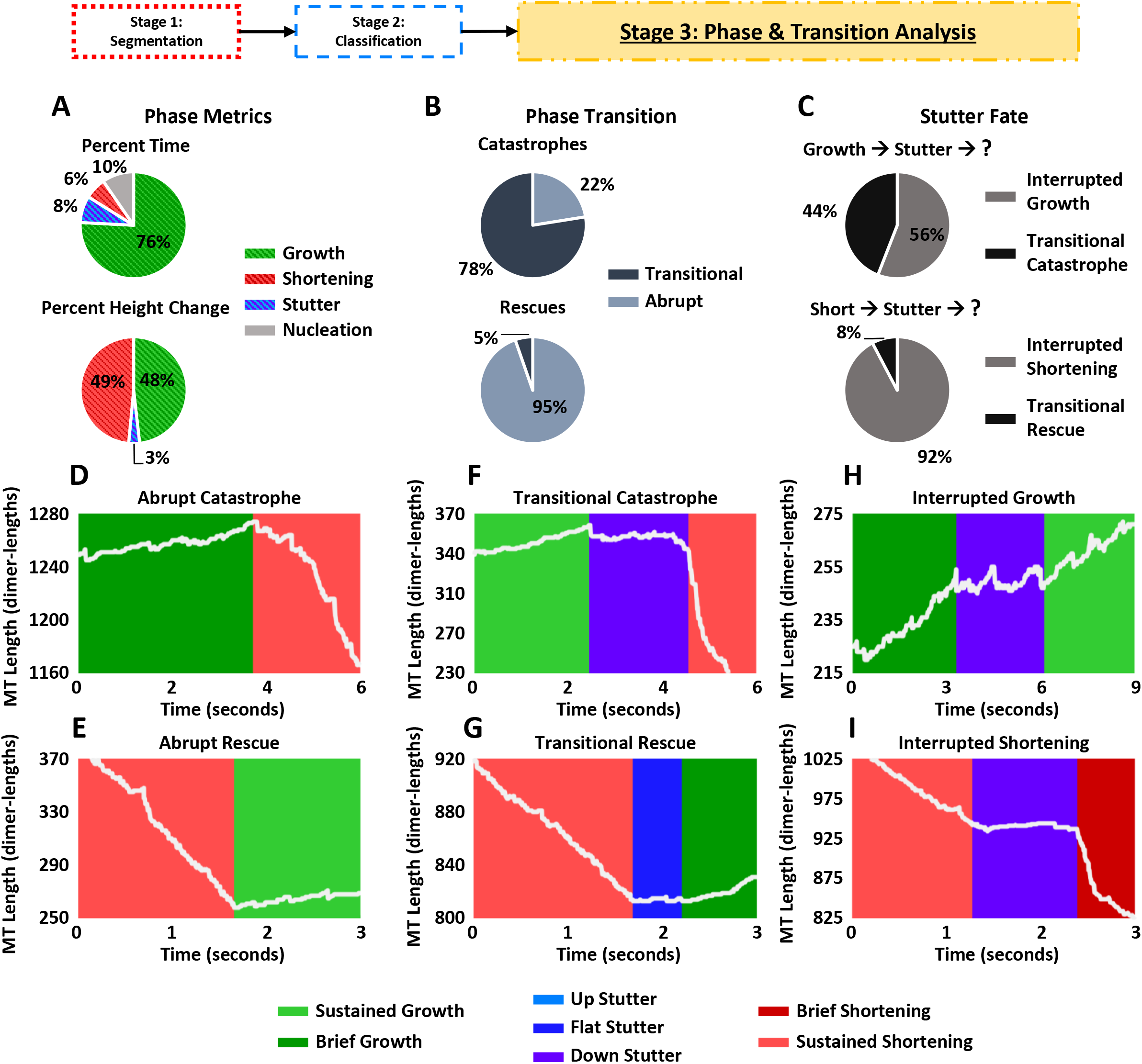
STADIA’s Phase and Transition Analysis of the dimer-scale *in silico* data. **(A)** Percent time spent in each class of phases/behaviors and percent height (MT length) change occurring during each class of phases/behaviors. A large majority of time is spent in growth. Notably, the *in silico* MTs spend more time in stutters than in shortening, emphasizing the importance of studying stutter behaviors. Most height change occurs during growth and shortening phases, as expected. **(B)** Percentages of catastrophes (top) and rescues (bottom) that are transitional vs. abrupt (see **Figure 4** for definitions). These data show that catastrophes are mostly transitional, whereas rescues are overwhelmingly abrupt. **(C)** Examination of stutter fate. These data show that when growth-to-stutter occurs (top), interrupted growth is slightly more likely than transitional catastrophe. However, when shortening-to-stutter occurs (bottom), interrupted shortening is much more likely than transitional rescue. **(D-H)** Examples of abrupt/transitional catastrophes (D,F), abrupt/transitional rescues (E,G), and interrupted growth/shortening (H,I).

**Figure 6.**
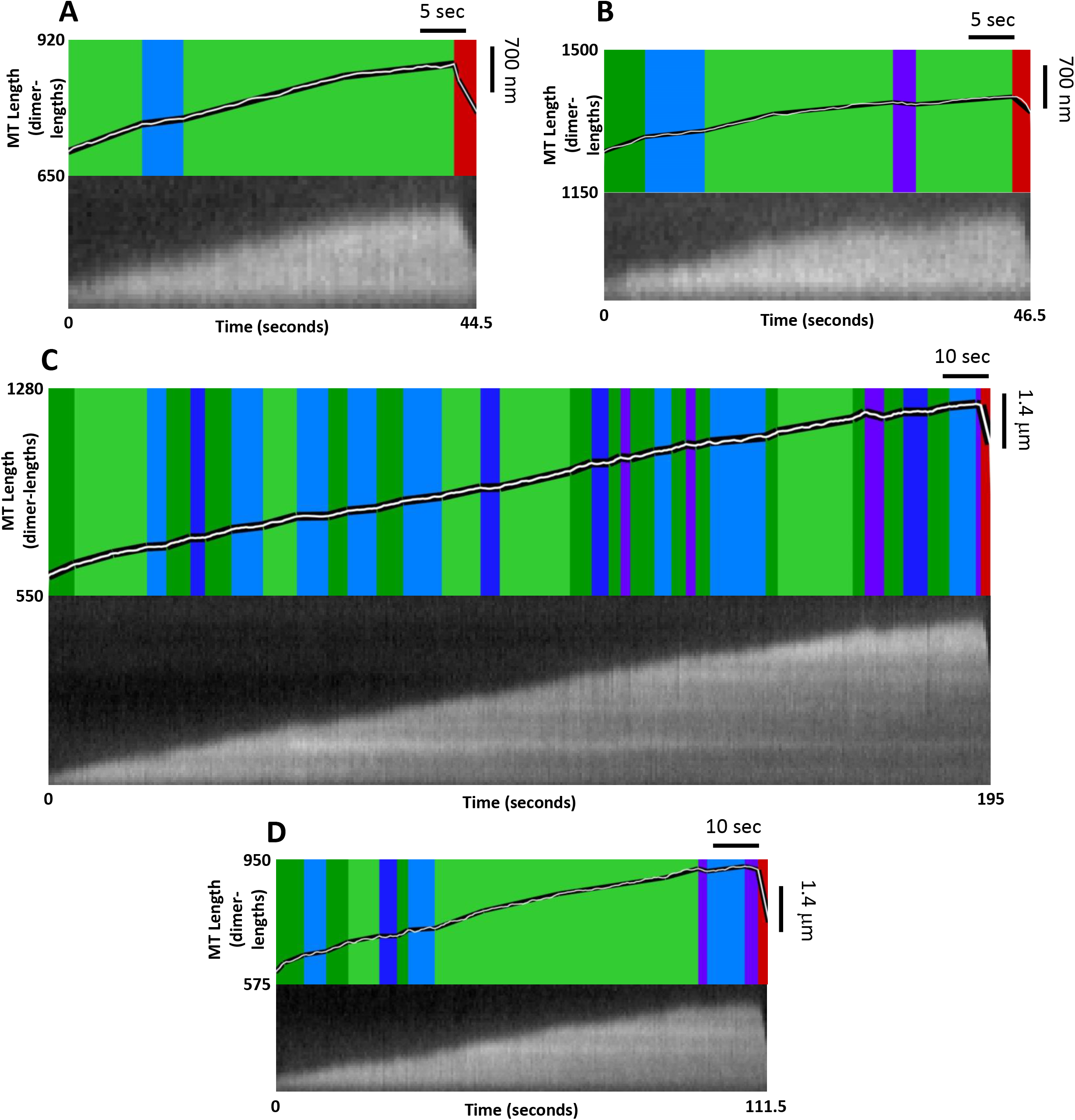
Alignment of STADIA length-history plots and their corresponding kymographs from the *in vitro* control dataset (upper and lower parts of each panel respectively, with STADIA colors as in **Figure 3** and scale bars as indicated). **(A,B)** Examples of abrupt catastrophes, where a shortening phase (red) directly follows a growth phase (green). **(C,D)** Examples of transitional catastrophes, where a shortening phase (red) follows one or more types of stutter (blue, purple). Note also the numerous stutters (blue, purple) that interrupt growth phases (green). The kymographs include examples of all three types of stutters that we distinguish based on slope: up stutters (light blue), flat stutters (dark blue), and down stutters (purple). The kymographs and the length-history traces inputted into STADIA were generated from the *in vitro* imaging data as described in the Methods section. The movies corresponding to each kymograph are provided as Supplemental Materials. Note that the movies are presented at 3.5 x real time (i.e., 3.5 x the time labeled on the kymographs and length-history plots).

**Figure 7.**
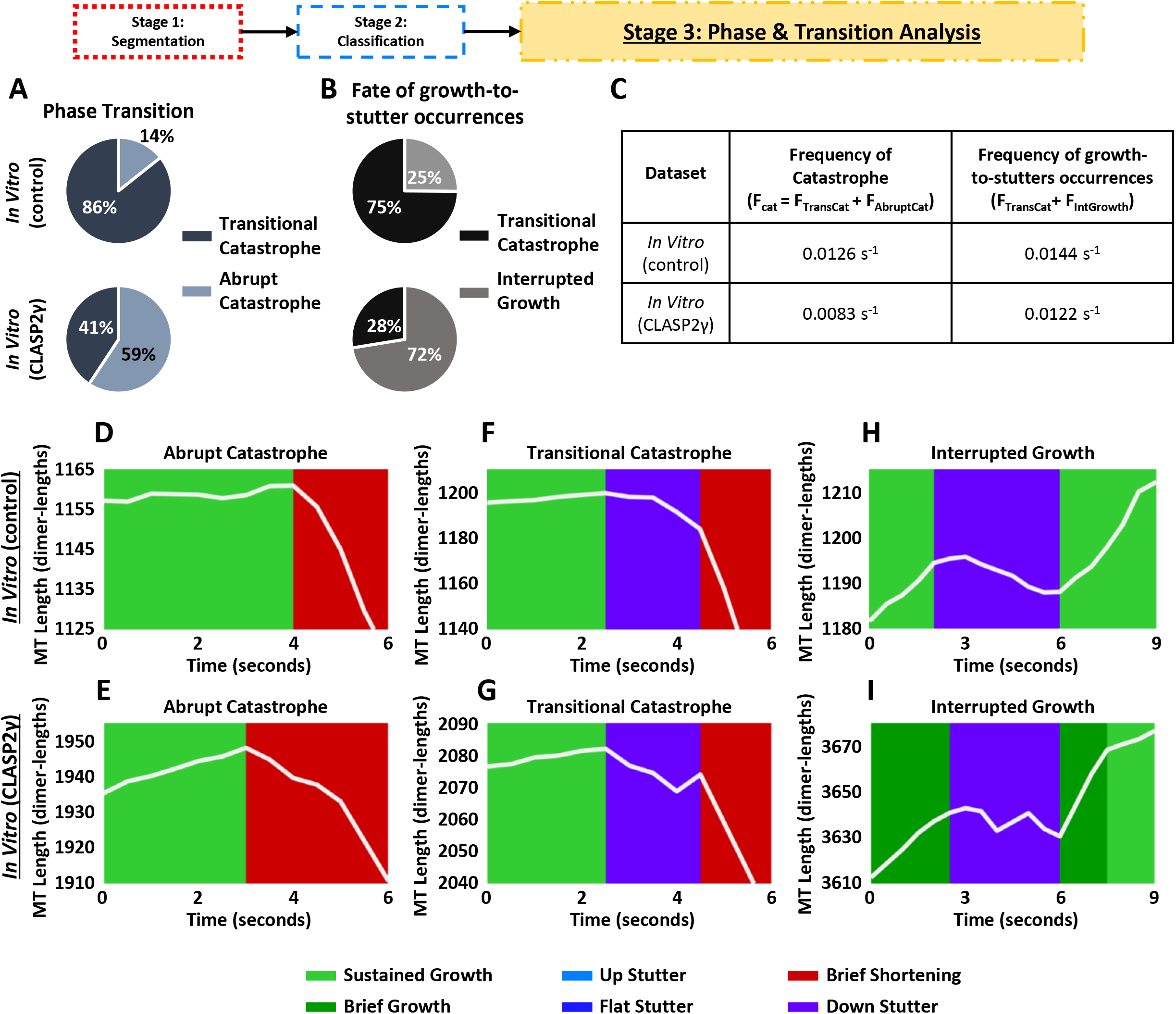
STADIA’s transition analysis of *in vitro* data: Effect of CLASP2γ on the nature of catastrophes and the fate of stuttering MTs. See **Figure 4** for transition definitions. **(A)** Consistent with what was seen for *in silico* MTs (**Figure 5B**), the majority of catastrophes for *in vitro* MTs without CLASP2γ are transitional (top). However, introduction of CLASP2γ increases the fraction of catastrophes that are abrupt (bottom). **(B)** Unlike the situation for the *in silico* MTs (**Figure 5C**), most growth-to-stutter occurrences for the *in vitro* MTs without CLASP (top) result in catastrophe. Addition of CLASP2γ (bottom) increases the likelihood that a stuttering MT returns to growth. **(C)** CLASP2γ decreases the overall frequency of catastrophe without greatly reducing the frequency of stutter-to-growth occurrences. Taken together, these data suggest that CLASP2γ reduces catastrophes by promoting growth following stutters without preventing stutters altogether. More specifically, CLASP2γ decreases the probability that a growth-to-stutter occurrence will proceed to shortening and increases the probability of returning to growth. In other words, transitions that would have been transitional catastrophes without CLASP2γ tend to become interrupted growths with CLASP2γ. **(D-I)** Examples of transitions as observed for the *in vitro* MTs both with (bottom) and without (top) CLASP2γ.

Similar chronological orderings of phases have previously been considered with pauses in the presence of cell extracts (Keller et al. 2008). Note that we report interrupted shortening and rescues only in the simulation data, because only short portions of *in vitro* shortening phases were captured (see examples in **Figure 6**).

### Catastrophes are usually preceded by stutters *in silico* and *in vitro*

Remarkably, when we examined the simulation data, we found that 78% of catastrophes commenced with a stutter, i.e., were transitional (**Figure 5 B**). A related observation is that almost half (44%) of stutters that occurred after a growth segment ended in catastrophe as opposed to returning to growth (**Figure 5 C**). A similar but more dramatic association between stutters and catastrophe was observed in the *in vitro* control data: 86% of catastrophes commenced from a stutter (**Figure 7 A**), and 75% of stutters from growth ended in a catastrophe (**Figure 7 B**).

In contrast to catastrophes, rescues as observed *in silico* rarely occurred with stutter. More specifically, only 5% of *in silico* rescues were transitional (i.e., few rescues initiated from a stutter) (**Figure 5 B**), and only 8% of stutters that occurred during depolymerization resulted in a rescue (**Figure 5 C**). Because we do not have sufficient data for rescues *in vitro*, we cannot make strong conclusions on the correlation between stutters and rescue in physical MTs. However, these results do suggest that catastrophe and rescue are not simply the mechanistic opposites of each other.

### CLASP2γ reduces the frequency of catastrophe by increasing the prevalence of interrupted growth

To further test STADIA's utility in analyzing dynamic instability and to examine both the prevalence and the significance of stutters, we compared the control *in vitro* dataset to the *in vitro* dataset in which the MTs were polymerizing in the presence of the MT binding protein CLASP2γ, which has been previously characterized as an anti-catastrophe factor (Lawrence and Zanic 2019; Aher et al. 2018). CLASP2 proteins are of interest to the biomedical community because they have been implicated in functions as diverse as kinetochore attachment (Girão et al. 2019), nervous system development (Dillon et al. 2017), and the insulin response (Kruse et al. 2017).

Recall that the clustering results, including detection of stutters, are similar for the control and CLASP2γ datasets (**Supplemental Figures S1.4 and S1.5**). However, dramatic differences in the transition frequencies between the CLASP2γ data and control *in vitro* data were observed when these data were examined quantitatively by STADIA.

First, the overall frequency of catastrophe in the presence of CLASP2γ was significantly reduced (**Figure 7 C** and **Supplemental Figure S1.10**). This observation itself is not surprising, given that previous work has shown that CLASP2γ reduces the frequency of catastrophe (e.g., (Sousa et al. 2007; Lawrence et al. 2018; Aher et al. 2018; Majumdar et al. 2018)). However, STADIA provides additional insight by distinguishing transitional catastrophes (growth-stutter-shortening) from abrupt catastrophes (growth-shortening). In particular, our results demonstrate that the reduction in overall catastrophe frequency was due to a large decrease in transitional catastrophe frequency, while the abrupt catastrophe frequency actually increased somewhat (**Figure 7 A** and **Supplemental Figure S1.10**).

Second, CLASP2γ slightly reduced the frequency of growth-to-stutter occurrences (i.e., F_TransCat_ + F_IntGrowth_, **Figure 7 C**), but not enough to account for the large decrease in transitional catastrophe frequency.

Third, and most striking, CLASP2γ increased the frequency of interrupted growth (growth-stutter-growth) (**Supplemental Figure S1.10**). More specifically, among transitions that began as growth-to-stutter, CLASP2γ increased the proportion of transitions that resulted in interrupted growth (growth-stutter-growth) and decreased the proportion of transitions that proceeded to transitional catastrophes (growth-stutter-shortening) (**Figure 7 B**). This change in proportions is the factor that accounts for most of the decrease in transitional catastrophe frequency.

Taken together, these data demonstrate that STADIA analysis provides information about CLASP2γ function not supplied by traditional analysis and indicates that CLASP2γ suppresses catastrophe at least in part by enabling stuttering MTs to re-enter growth (i.e., CLASP2γ tends to convert would-be transitional catastrophes into interrupted growths). This idea is supported by recent reports that MTs can withstand greater growth rate variability without undergoing catastrophe in the presence of CLASP2γ (Lawrence et al. 2018; Lawrence and Zanic 2019) and that CLASP2γ can protect against catastrophe in the presence of lagging protofilaments (Aher et al. 2018).

### Robustness of conclusions over varying values of input parameters and data acquisition rates

The data above lead us to conclude that stutters (previously observed but not quantified in detail) are strongly associated with spontaneously occurring catastrophes, both *in silico* and *in vitro*. Our STADIA analysis also indicates that the anti-catastrophe factor CLASP2γ reduces catastrophe by increasing the fraction of stuttering microtubules that return to growth rather than entering shortening phases. An important remaining question is whether these conclusions are robust to variations in the STADIA input parameters.

To address this question, we performed a parameter sweep by analyzing all three datasets (i.e., *in silico* as well as *in vitro* with and without CLASP2γ) using a range of values for each of the key parameters in STADIA, namely the Minimum Segment Duration and the Maximum Error Tolerance. The values of the Minimum Segment Duration and the Maximum Error Tolerance determine how closely STADIA’s piecewise linear approximation matches the raw length-history data inputted into STADIA. As discussed above, the piecewise linear approximation is produced in the segmentation stage and then used by the classification stage, followed by the phase & transition analysis stage (see **Figures 2** and **S1.1** for overview of STADIA stages). The results of the parameter sweep analysis (gap statistic plots, clustering profiles, phase-labeled length-history plots, and transition analysis) are provided in **Supplemental Material Section 2.**

The parameter sweep results indicate that the conclusions above are indeed robust as long as the parameters are kept within ranges relevant to the scale of the dynamics being studied. For example, for the *in silico* data, both the values of the frequencies of abrupt and transitional catastrophe and the ratio between them are relatively insensitive to changes in the Maximum Error Tolerance in the range of 10-40 dimer-lengths (i.e., 80-320 nm) (**Supplemental Figure S2.9**). Moreover, the conclusion that most catastrophes are transitional is robust for Minimum Segment Duration values of 1 second or less. However, for Minimum Segment Duration values of 1.5 seconds or greater, abrupt catastrophes become more common (**Supplemental Figure S2.9**). The observation that catastrophes become more abrupt as the Minimum Segment Duration increases occurs simply because fewer stutters are detected and therefore fewer catastrophes are recognized as transitional. This observation is illustrated in examples of the labeled length-history plots (**Supplemental Figure S2.4**). Importantly, the overall catastrophe frequency (the sum of abrupt and transitional) is less sensitive to changes in Minimum Segment Duration and Maximum Error Tolerance than are the abrupt and transitional catastrophe frequencies. The situation is similar for the *in vitro* data (**Supplemental Figures S2.8, S2.10, S2.11**), though it is a bit noisier, likely because the amount of data is smaller (especially with CLASP2γ). Importantly, the conclusion that CLASP2γ reduces catastrophe by increasing the frequency of interrupted growth events is robust to changes across wide ranges of both Minimum Segment Duration and Maximum Error Tolerance (**Supplemental Figures S2.10, S2.11, S2.12**).

In addition, we also tested the effect of varying the data acquisition rate using the *in silico* dataset (**Supplemental Material Section 3**). Examining the data acquisition rate is significant given that the *in silico* dataset records every dimer-scale biochemical event (bond formation/breaking, hydrolysis; on the scale of >1000 data points per second; see Methods). In contrast, frame rates in experiments vary from more than 10 frames per second (e.g., (Mickolajczyk et al. 2019)) to fewer than 0.3 frames per second (e.g, (Gierke, Kumar, and Wittmann 2010)). To perform these tests, we took the original full resolution *in silico* dataset and extracted the data at the data acquisition time steps indicated in the figures (frame rate is the inverse of the time step, e.g., 2 fps corresponds to a time step of 0.5 sec). These data (figures in **Supplemental Material Section 3**) show that the conclusion that up stutters exist is robust for data acquisition time steps up to 3 seconds, and similarly for down stutters at data acquisition time steps up to 1 second, assuming reasonable choices for Maximum Error Tolerance and Minimum Segment Duration (see above, also discussed more below). However, even when stutters are detected as a distinct cluster, the number of stutter segments detected generally decreases for larger data acquisition time steps (i.e., slower data acquisition rates). This observation is not surprising because some stutters, particularly down stutters for the *in silico* data, have durations on the order of one second (**Supplemental Figure S1.8**), and with frame rates less than one second, such stutters would be undetectable.

In closing, these parameter testing data show that to detect stutters, one must use spatial and temporal parameters that have resolution sufficiently high to allow detection of stutters, but not so high that the analysis is “distracted” by protofilament extensions (as are recorded *in silico* and perhaps *in vitro*) or experimental noise. For our *in silico* dataset these ranges were empirically determined to be from 15-30 dimer-lengths (i.e., 120-240 nm) for the Maximum Error Tolerance and 1 second or less for the Minimum Segment Duration. For our *in vitro* (control) dataset, ranges were determined to be from 10-20 dimer-lengths (80-160 nm) for the Maximum Error Tolerance and 1 second or less for the Minimum Segment Duration. Note that the *in vitro* dataset was obtained using a frame rate of 2 fps, which was determined to be near the slower end of the range of acceptable data acquisition rates for some of the conclusions; it is possible that with a faster frame rate, *in vitro* data may tolerate a wider range of STADIA parameters (similar to the *in silico* dataset). Significantly, short data acquisition time steps do not introduce problems (indeed, they are ideal, as seen with the full resolution *in silico* dataset) because the Maximum Error Tolerance and Minimum Segment Duration parameters prevent the segmentation (piecewise linear approximation) of MT length-history data from containing arbitrarily short segments.

## DISCUSSION

Here we have presented STADIA, a data-driven, automated tool for performing dynamic instability analysis using length-history data as input. Using STADIA, we have quantified stutters and their associated transitions (**Figures 3,5-7, Table 1**, and **Supplemental Figures S1.8-S1.10**). Stutters are a set of dynamic behaviors that can be distinguished from typical growth or shortening; the primary distinguishing factor is that stutters on average have slower rates of MT length change. Stutters are also distinguishable from pauses in that a pause is typically described as a period of time when the MT neither grows nor shortens. Our analysis shows that stutters (previously observed but not quantified in detail) are strongly associated with spontaneously occurring catastrophes, both *in silico* and *in vitro* (**Figures 5,7**). Our STADIA analysis also indicates that the anti-catastrophe factor CLASP2γ reduces catastrophe by increasing the fraction of stutters that return to growth rather than entering shortening phases (**Figure 7**). Importantly, we have shown that these results are robust across a range of parameter values (**Supplemental Material Section 2**) and are compatible with data acquired across a range of temporal resolutions (**Supplemental Material Section 3**).

### Mechanisms of stutters and implications for the process of catastrophe

What causes stutters, especially those that disrupt growth, and why are they associated with catastrophe? A fundamental component to answering this question comes from recognizing that when transitioning from growth to stutter, there is a net decrease in the number of subunits (tubulin dimers) that are incorporated into the MT per unit time. This net decrease could occur because new subunits attach to the tip less frequently than during normal growth, or because bound subunits leave the tip more frequently than during growth, or a combination of these two.

While simple stochastic fluctuations in subunit arrival or departure could potentially contribute to stutters, examination of length-history plots (**Figures 3, 5-7**) demonstrates that stutter segments occur at a more macroscopic scale than the small stochastic fluctuations that occur within growth, shortening and stutter segments. Changes in rates of attachment and detachment could also result from changes in tip structure. However, one could argue that the rate of subunit attachment should not vary with tip structure: assuming that longitudinal bonds form first, there are always 13 landing sites for new subunits (Castle and Odde 2013). Therefore, we suggest that stutters following growth segments likely result from a situation where an unusually large fraction of incoming subunits detach from the tip structure without being fully incorporated into the lattice, e.g., because tip taper or other structural features like the presence of GDP-tubulin make it difficult for lateral bonds to form. In other words, we suggest that stutters occur when the structure of the tip is such that the subunit detachment rate is unusually high compared to the average detachment rate during growth.

This reasoning provides a potential explanation for the correlation between stutter and catastrophe: if the fraction of incoming subunits incorporated into the lattice is smaller than during normal growth periods, the stabilizing cap of GTP-tubulin at the MT end will become smaller, the likelihood of exposing GDP-tubulin subunits will increase, and the possibility of complete cap loss (catastrophe) will rise. At present, these ideas are speculation, but future work may be able to shed light on these hypotheses (see also related discussions in papers including (VanBuren, Cassimeris, and Odde 2005; Howard and Hyman 2009; Gardner et al. 2011; Margolin et al. 2012; Coombes et al. 2013; Zakharov et al. 2015; McIntosh et al. 2018)).

Furthermore, the mechanisms could vary for different types of stutters. As demonstrated in the Results, STADIA distinguishes up, down, and flat stutters, and distinguishes stutters that occur as part of interrupted growth, interrupted shortening, transitional catastrophe, and transitional rescue. Thus, as a tool for comprehensively identifying multiple types of stutters, STADIA lays the groundwork for future mechanistic studies.

### Comparison of the *in silico* and *in vitro* results

The behaviors observed in the dimer-scale simulation data and the experimental data are qualitatively similar. In particular, both types of data support the prevalence of stutters throughout length histories and the role of stutters in catastrophes. The differences in the particular numerical values of measured quantities are not surprising, because the simulation parameters were tuned in (Margolin et al. 2012) based on a different experimental dataset (Walker et al. 1988) from the experimental datasets used here (a subset of which was used in (Lawrence et al. 2018)). The qualitative similarities between the results from the different datasets provide evidence that the observed trends are not specific to one experimental preparation or one type of tubulin (e.g., 10 μM pure porcine tubulin in (Walker et al. 1988), versus 12 μM bovine tubulin with EB1 and with or without CLASP2γ here and in (Lawrence et al. 2018)). Furthermore, comparison with the negative control (two-state growth-shortening model, **Supplemental Material Section 4**) demonstrates that the existence of stutters in the dimer-scale simulations and the *in vitro* data is not manufactured by STADIA.

### Relationship to previous work

#### Distinguishing stutters and previously identified pauses

Pauses have most frequently been observed *in vivo* (see citations in the Introduction), and are likely caused by MT binding proteins (Moriwaki and Goshima 2016) and other factors external to the MTs themselves (e.g., reaching the cell edge (Rusan et al. 2001; Komarova, Vorobjev, and Borisy 2002)). In contrast, *in vitro* pauses in the absence of drugs or MTBPs are rare (Walker et al. 1988). The observation that stutters are prevalent in both our *in silico* and *in vitro* datasets suggests that stutters are an intrinsic component of DI itself.

Gierke and colleagues have previously described “*bona fide* pauses” as phases “during which no polymerization or depolymerization occurs.” Due to physical detection limits, true pauses would be indistinguishable from periods of very slow polymerization or depolymerization that do not meet the detection threshold (Gierke, Kumar, and Wittmann 2010). Particularly in older datasets with large thresholds (e.g., a length-change threshold of 0.5 microns), most stutters either would have been considered pauses or would not have been separated out from larger growth or shortening phases at all. With newer imaging technology, data can be obtained at higher temporal and spatial resolution (e.g., (Maurer et al. 2014; Duellberg et al. 2016; Rickman et al. 2017; Guo et al. 2018)), which can enable the distinction of stutters and pauses.

For most stutters, a measurable net length change does occur over the course of the stutter segment: up and down stutters occur much more often than flat stutters. Using this information about stutters from our results and definitions of pauses already existing in the literature, we propose the following operational criteria for distinguishing pauses and stutters: pauses are periods during which no detectable length change occurs, whereas stutters are periods during which the MT structure changes but with slower net rates of length change than typical growth and shortening phases. In datasets that contain both stutters and pauses, the current version of STADIA would classify *“bona fide* pauses” as flat stutters. Future work will be needed to determine if it would be meaningful to apply criteria to distinguish flat stutters, which generally have very short time durations, from pauses.

#### Previously observed behaviors that are similar to particular types of stutters

Maurer et al. observed short episodes of pause or slow growth prior to catastrophes in experiments with EB1 (Maurer et al. 2014). These pre-catastrophe slowdowns are analogous to transitional catastrophes in our terminology. Pauses or slowdowns prior to catastrophe have also been observed in cases where the catastrophe is induced by outside factors such as mechanical force (Janson and Dogterom 2004) or reduction in tubulin concentration ((Duellberg et al. 2016; Duellberg, Cade, and Surrey 2016), similar to predictions based on simulations in (Margolin et al. 2012)). In contrast, the catastrophes in our datasets occur spontaneously as part of DI; in the *in vitro* datasets, EB1 and CLASP2γ affect the frequency of catastrophe, but the catastrophes still occur stochastically over time, as opposed to being induced by an experimenter at a particular moment.

The episodes of slow growth in Rickman et al. (Rickman et al. 2017) bear some similarity to stutters interrupting growth as identified by STADIA. However, the slow growth episodes of Rickman et al. occurred rarely (2 to 5 occurrences, ~0.26% to ~6.1% of the time analyzed, depending on tubulin concentration). These episodes appear to correspond to the most extreme of our stutters, meaning the stutters with the longest time durations or with the most variability in length during the stutter.

Based on analysis of variability in growth rates in experimental data, Odde et al. proposed a model with “near catastrophes”, which are similar to stutters interrupting growth (David J. Odde, Buettner, and Cassimeris 1996). They suggested that the largest of the “near catastrophes” may correspond to previously observed pauses and that the smaller “near catastrophes” would not be easily detected by eye (the time between data points in their analysis was ~ 3 seconds).

Building beyond this previous work, STADIA provides a comprehensive method for detecting and quantifying multiple types of stutters, and distinguishing phase transitions that include stutters from those that do not.

#### Differences between STADIA and previous segmentation/classification methods

Although STADIA identifies DI phases at a finer scale than many existing DI analysis methods, STADIA differs from methods that simply consider individual displacements between frames and label them as growth, shortening, or pause using thresholds on the length change (e.g., (Komarova, Vorobjev, and Borisy 2002; Guo et al. 2018)). In contrast to such methods, STADIA identifies larger-scale segments during which a MT exhibits a consistent behavior.

Similar to STADIA, many existing time-series analysis methods that have been used in other applications (e.g., identifying runs and pauses in the transport of organelles along MTs by motor proteins (Zaliapin et al. 2005)) involve a segmentation step (e.g., (Zaliapin, Gabrielov, and Keilis-Borok 2003)) that is often followed by a classification step (e.g., (Fu 2011)). To our knowledge, such methods have not been previously applied to MT dynamic instability data. In contrast, many DI analysis methods essentially perform classification before segmentation, by setting thresholds for classifying growth, shortening, and possibly pause or slowdown periods, and then applying the thresholds to identify segments in the data (e.g., (Dhamodharan and Wadsworth 1995; Fees, Estrem, and Moore 2017; Kiris, Ventimiglia, and Feinstein 2010)). Additionally, unlike existing methods that use arbitrary pre-defined thresholds on segment features (length change, time duration, and/or slope), STADIA uses a data-driven approach to identify emergent clusters in the segment feature data.

#### Differences between STADIA and previous phase transition analysis

In regard to phase transition analysis, several previous articles grouped their pauses with growth when defining catastrophe and rescue; by their definitions, a catastrophe is a transition from growth or pause to shortening, and a rescue is a transition from shortening to growth or pause (Panda et al. 1996; Dhamodharan et al. 1995; Dhamodharan and Wadsworth 1995; Rusan et al. 2001; Kamath, Oroudjev, and Jordan 2010; Yenjerla, Lopus, and Wilson 2010; Kiris, Ventimiglia, and Feinstein 2010; Moriwaki and Goshima 2016). By these definitions or analogous definitions with stutter in place of pause, an episode of interrupted shortening would be labeled as a rescue followed by a catastrophe, whereas an interrupted growth would not be distinguished from *un*interrupted growth. STADIA improves upon typical transition analysis by considering all possible transitions between growth, shortening, and stutter (similar to the transitions among growth, shortening, and pause considered in (Keller et al. 2008)). Such transition analysis enables more in-depth investigation of the mechanisms of DI and DI-regulating proteins. For example, the observation that CLASP2γ tends to convert would-be transitional catastrophes into interrupted growths would not have been possible without a method that is able to identify transitional catastrophes and interrupted growths.

### Conclusions

The key results of this work are four-fold: (1) the use of STADIA to more thoroughly quantify and examine ‘stutters’, a previously observed category of behaviors during which MTs undergo slow rates of overall length change compared to growth or shortening phases; (2) the observation that stutters are strongly associated with catastrophe in dimer-scale *in silico* and TIRF-imaged *in vitro* data; (3) the evidence that the anti-catastrophe factor CLASP2γ reduces catastrophe by increasing the fraction of stutters that return to growth rather than enter shortening phases; (4) the development of STADIA as an improved analytical tool for quantification of MT behavior, as exemplified by the first three key results. Our results concerning the detection of stutters differ from previous work in that STADIA comprehensively and systematically identifies all types of stutters (up stutter, flat stutter, down stutter) across length-history data and considers all possible transitions among growth, shortening, and stutters. We suggest that quantification of stutters in future DI analysis through STADIA or similar tools will enable improved analysis of MT dynamics that is more complete, precise, and reproducible. The clearer picture that results from this analysis will facilitate investigation of the mechanisms of catastrophe and rescue as well as the activities of the MT binding proteins that regulate these transitions.

## METHODS

### CLASSICAL DI ANALYSIS

For purposes of comparison to STADIA, we use the custom DI analysis program written in MATLAB and described in the Supplemental Methods of (Jonasson et al. 2020), which we refer to here as the “peak-valley method” (our implementation of classical DI analysis). Briefly, this method segments growth and shortening phases by first identifying major peaks and valleys in the length-history data using the MATLAB function *findpeaks()*. Then the ascent to each major peak is classified as a growth segment and the descent from the peak is classified as a shortening segment. Each major peak is classified as a catastrophe, where the end of growth and the start of shortening are identified as occurring at the same time point. A major valley is classified as a rescue only if the MT length at the time of the major valley is greater than or equal to a user-defined value called the ‘minimum rescue length’, in which case the end of shortening and the start of growth are identified as the same point. For a major valley that occurs below the minimum rescue length, the end of shortening can be identified as an earlier point in time than the start of growth, in which case the time between these points would correspond to a nucleation period near the MT seed (see Supplemental Methods of (Jonasson et al. 2020) for additional details). For the classical DI analysis in this paper, the minimum prominence for major peaks (i.e., minimum height change between a major peak and the nearest major valley) in the peak-valley method was set equal to the Maximum Error Tolerance in STADIA. The minimum peak height and the minimum rescue length in the peak-valley method were set equal to the sum of the Nucleation Threshold plus the Maximum Error Tolerance in STADIA (see **Supplemental Table S1.1**).

In the peak-valley method results shown in **Table 1**, the V_growth_ and V_short_ calculations relied on linear regressions fitted to each growth or shortening segment. V_growth_ was calculated as the arithmetic mean of the slopes of the regression lines for all growth segments whose linear regression had an R^2^ value of at least 95%. V_short_ was calculated in the same manner using the shortening segments. F_cat_ was calculated as the total number of catastrophes divided by the total time spent in growth phases. Similarly, F_res_ was calculated as the total number of rescues divided by the total time spent in shortening phases. See **Table 1** for comparison of the DI parameters measured from the peak-valley method and from STADIA.

### STATISTICAL TOOL FOR AUTOMATED DYNAMIC INSTABILITY ANALYSIS: <u>STADIA</u>

This section outlines the three major stages of STADIA analysis (Segmentation, Classification, and Phase and Transition Analysis; **Figure 2, Supplemental Figure S1.1**) as well as the parameters used for the inspection of *in silico* and *in vitro* data using STADIA. Refer to **Supplemental Table S1.1** for a complete list of all STADIA user-defined parameters used for analysis of both *in silico* and *in vitro* MTs.

#### Segmentation Stage

In the segmentation stage, STADIA takes MT length-history data and generates a continuous piecewise linear approximation of the MT length-history plot. Segmentation also includes a preprocessing step that prepares the user's length-history data for input into STADIA and a post-processing step that prepares the results of the segmentation step for classification.

##### Preprocessing

As an initial step, STADIA automatically formats the inputted MT length-history data into a single time series of length-history data points. MT length-history data can be provided either as a long-time observation of a single MT (possible with simulations) or as a series of length histories of multiple MTs (common with experimental data). In the latter case, STADIA automatically connects, or ‘stitches’, the data from multiple MTs (with separators in between) into a single time-series representation of MT length-history data (e.g., **Figure 1 D**). Note that special treatment of the stitching separator between observations allows the segmentation to avoid misclassification of stitch boundaries as transitions. This preprocessing step allows STADIA to conduct analysis for both simulation data and experimental data in a similar and consistent manner.

In this manuscript, the simulation data were provided as one long time series from an individual MT (no stitching required), while the *in vitro* data (both with and without CLASP2γ) were obtained from multiple MTs over a shorter period of observation (for details, see Data Acquisition: *In Vitro* Microtubule Experiments below). Because long shortening phases were not captured (for technical reasons), the data from a specific MT were broken into samples that typically consisted of a growth phase followed by an initial depolymerization and then termination of that observation. STADIA first placed individual length-history samples for a given MT consecutively into the same time-series plot, and then stitched all of the data for all of the MTs within each experiment. Note that the clustering methods used in STADIA require a dataset large enough to display a rich variety of possible DI behaviors. Therefore, instead of analyzing each individual *in vitro* MT for various behaviors, it is necessary to collectively consider multiple MTs from the same experiment so there is enough length-history data to classify. Thus, separately for the control and CLASP2γ *in vitro* datasets, this stitching procedure combined all the available behavior into a dedicated single time-series representation (one with CLASP2γ and one without).

##### Segmentation

STADIA takes the single time-series graph produced by the preprocessing step and performs segmentation as an adaptive, iterative process according to restrictions provided by user-defined thresholds. The segmentation process begins by identifying major peaks and valleys (i.e., local extrema) in MT length-history data using the *findpeaks()* function in MATLAB. Consecutive extrema are connected by line segments to form an initial linear approximation of the length-history data (**Figure 2 C**). An initial list of vertices is defined by these peaks and valleys. New vertices are added to mark the boundaries between which the MT lengths are below a ‘nucleation’ threshold length, generally chosen to be near the limits of observation in experimental conditions. Then, the iterative process seeks to include new vertices to define line segment endpoints. This improves the approximation accuracy by constructing a continuous piecewise linear approximation that satisfies the user-defined parameters of Maximum Error Tolerance and Minimum Segment Duration (**Figure 2 D**). Note that the segmentation algorithm implemented in STADIA is similar to the ‘Top-down algorithm’ described in (Keogh et al. 2001). The segmentation algorithm can be explained in the following steps:

1. Let {*x*^1^, *x*^2^,…, *x*^*N*^} represent the peaks, valleys, and nucleation threshold points that serve as the initial list of vertices (i.e., segment endpoints), where *x*^1^ and *x*^*N*^ are the first and last points of the length-history data, respectively.
2. For any *i* = 1,…, *N* − 1, define the *i*^th^ region as the interval between the consecutive pair of initial vertices, *x*^*i*^ and *x*^*i*+1^. Construct a line segment with end points as 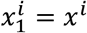 and 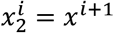 such that the vertices corresponding to the *i*^th^ region are 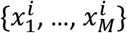, initially *M* = 2, but we seek to grow this list in the following steps.
3. For *j* = 1,…, *M* − 1, consider the *j*^th^ line segment in the *i*^th^ region defined by 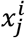 and 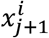. Calculate the error between this line segment and the corresponding points on the original length-history data.
  ○ If the maximum error is greater than the user-defined Maximum Error Tolerance, then the error condition is not satisfied, and a new data point needs to be included in the vertex list. Proceed to step 4.
  ○ If the maximum error from this segment is less than the user-defined Maximum Error Tolerance, then the error condition is satisfied for the *j*^th^ line segment in the *i*^th^ region. Proceed to step 6.
4. Define the data point where the greatest error occurs found in step 3 as 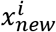
  ○ If 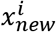 violates the user-defined Minimum Segment Duration, attempt to choose the closest point in the length-history data that would satisfy both the Maximum Error Tolerance and Minimum Segment Duration.
5. Include the newly identified vertex into the list of vertices for the *i*^th^ region. This will require redefining indices to preserve ordering. For example, for the first new vertex added to the *i*^th^ region, the original single segment in the *i*^th^ region is broken into 2 segments, and the list of vertices corresponding to the *i*^th^ region is now defined as

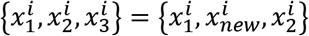

where the vertex list on the right side is indexed according to the previous iteration, and the updated vertex list on the left side replaces the list defined in step 2, such that 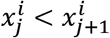 for all *j* = 1,…, *M* − 1.
6. Repeat steps 3 through 5 until the error condition is satisfied without adding new vertices into the *i*^th^ region.
7. Repeat steps 2 through 6 for all *i* ≤ *N* − 1.

The final result is a continuous, piecewise linear approximation that fits the entire length-history dataset according to a user-defined Maximum Error Tolerance (excerpts of the full length-history approximation are illustrated in **Figure 2 D**, and the black lines plotted in **Figure 3 G,H**). The vertices of the piecewise linear approximations provide line segments with endpoints at moments where significant changes in slope occur in length-history plots. Thus, the activity covered by each segment between endpoints represents a consistent period of MT length-history behavior that can be identified as belonging to a DI phase in the classification stage.

The following user-defined parameter values set the accuracy of the piecewise linear approximations for all analysis in the main text and **Supplemental Material Section 1**: Minimum Segment Duration = 0.5 seconds; Maximum Error Tolerance = 20 dimer-lengths (**Supplemental Table S1.1**). These parameters are varied over a range of values in **Supplemental Material Sections 2** and **3**.

##### Justification for segmentation method

To create a more accurate approximation of MT length-history data compared to more classical ‘peak-valley’ methods, STADIA employs the iterative approach described above to create a piecewise linear approximation of the MT length-history data satisfying the user-defined Maximum Error Tolerance. We chose this approach because it provides a simple method for identifying points that may not necessary be peaks or valleys, but where a change from one sustained MT behavior to another occurs. Through the Maximum Error Tolerance choice, the user is able to regulate the accuracy of the linear approximation. Through the Minimum Segment Duration choice, the user is able to perform the analysis of MT length-history data at the desired timescale. An assumption of performing segmentation in this manner is that MT behavior follows a linear trend at the timescale being analyzed. Finally, a difference we note is that this method results in a continuous piecewise linear approximation whereas some other segmentation methods produce discontinuous approximations (e.g., (Zaliapin, Gabrielov, and Keilis-Borok 2003)).

##### Post-processing to prepare for classification

Line segments from the piecewise linear approximation each have measured slopes, time durations, and height changes (**Figure 2 E**); this set of measurements provides a 3-D feature space where the segments reside (see below for *Justification for classification feature space*). However, some post-processing is needed before submitting this dataset to the classification process. First, the periods of time when the MT length is less than the ‘nucleation’ threshold described above are excluded from further analysis. Next, line segments are identified as flat stutters if either their total height change or slope magnitude are below user-defined thresholds. Flat stutters, which are not characterized as growth or shortening, are set aside until the end of the classification procedures. The remaining positive and negative sloped segments are considered separately during the next stage (classification).

In this work, line segments containing MT lengths less than 75 dimer-lengths were considered nucleation segments. A segment was identified as a flat stutter if the absolute value of its total height change was less than 3 dimer-lengths or the absolute value of its slope was less than 0.5 dimer-lengths/second (**Supplemental Table S1.1**).

##### Justification for classification feature space

Mathematically, knowing the values of only two of time duration, height change, and slope provides sufficient information to calculate the value of the third variable. However, we use all three variables in the clustering step because some data points that are well separated in the three-dimensional space would become indistinguishable for all practical purposes if only two of the variables were used (**Supplemental Figures S1.2** and **S1.3**). Additionally, which data points become indistinguishable would depend on which pair of variables was used (time duration and height change, time duration and slope, or height change and slope).

The slope = height/time surface (**Supplemental Figures S1.2A** and **S1.3E**) could be parameterized with only two variables in a way that would maintain the separation from the three-dimensional space. However, these two new variables would be some combination of the original three variables, and these combinations would not necessarily have clear physical meanings. We therefore chose to use all three of the basic variables (time duration, height change, and slope) to maintain a more direct connection to the biology.

The use of nonlinear combinations of variables is not uncommon in statistics. Additional combinations of our three basic variables as well as other variables may be worth exploring in future work that aims to further dissect MT length-history behaviors. For the purposes of the present work, the three basic variables are sufficient for detecting distinct clusters within the positive and negative slope groups.

#### Classification Stage

To begin the classification stage, STADIA takes the results of the segmentation stage (segregated positive and negative slope line segments that approximate the MT length-history data) and analyzes them using *k*-means clustering (Lloyd 1982; Macqueen 1967), where the number of clusters, the *k*-value, is suggested by the gap statistic (Tibshirani, Walther, and Hastie 2001).

##### Justification for using k-means clustering

As an unsupervised clustering algorithm, *k*-means does not require prior knowledge of the characteristics of the clusters to be found. Rather, *k*-means groups together data points that share similar characteristics (i.e., data points that are near each other in a relevant feature space). *K*-means also has advantages of its ease of use and interpretability. Ideal datasets for *k*-means have globularly shaped clusters (i.e., each cluster would follow a Gaussian distribution). Although our data is not Gaussian, *k*-means still provides an objective methodology to find substructures in the overall data structure. The observation that *k*-means enables us to identify and quantify stutters (behaviors that have been noted previously but not quantified in detail) indicates that it provides a useful methodology for categorization and quantification of MT behavior.

##### Pre-processing

*K*-means clustering uses Euclidean distance (i.e., straight-line distance between two points in 3-D space) as the primary measurement in its algorithm to classify data. Therefore, all features should exist on the same scale so as to give each feature equal weight in the *k*-means classification process. To meet this requirement, the segment features (slope, height change, and time duration values) were transformed by first being log-scaled and then standardized with respect to each feature’s statistics (i.e., by subtracting the mean and dividing by the standard deviation). Scaling and standardizing the data in this way is a common practice for analysis utilizing *k*-means clustering and allows for all features to be considered on the same scale to better suit the Euclidean distance used in *k*-means clustering (Hastie, Tibshirani, and Friedman 2009).

##### Determining appropriate number of clusters for each dataset

As noted at the beginning of the Results and Discussion section, the second goal for STADIA development was that it be impartial to the number of behaviors exhibited by MTs, thus removing any assumptions about MT dynamics being restricted to two behaviors (i.e., only growth and shortening). The *k*-value (i.e., number of clusters to use in *k*-means) is determined for positive and negative slope segments separately in the **Diagnostic Mode** of STADIA. This process utilizes the ‘gap statistic’, which compares the within-cluster dispersion to a null reference distribution when seeking the optimal number of clusters that best separates the data during *k*-means clustering (i.e., the gap statistic answers the question: ‘what number of clusters results in the best separation between the clusters?’) (Tibshirani, Walther, and Hastie 2001). The gap statistic is the primary driver in determining the proper *k*-value for clustering the line segment data. However, it is also recommended for the user to check how well the number of clusters suggested by the gap statistic describes the dataset qualitatively. Typically, the optimal *k*-value corresponds to the first local maximum of the gap-statistic plot. In some cases, however, qualitative inspection of the data may suggest that the first local maximum is not optimal, in which case the next local maximum should be used.

For the purposes of informing the optimal *k*-value for use in *k*-means clustering, STADIA repeats clustering procedures for different *k*-values, ranging from 1 through 12, using 100 random starts for each value (*k*-means clustering does not necessarily converge to a global maximum so multiple starts are required to determine optimal centroid locations). Simultaneously, STADIA measures the corresponding gap statistic for each value of *k*. As explained above, in our analyses the *k*-value corresponding to the first local maximum of the gap statistic plot was usually chosen as the optimal number of clusters (**Figure 3 A,B,D,E, Supplemental Figures S1.4,S1.5**). However, for example, qualitative inspection of the clustering for the positive slope segments of the *in vitro* control MTs (i.e., without CLASP2γ) and comparison to the other datasets suggested that the first local maximum greater than *k*=1 (i.e., *k*=3) described the data more appropriately (**Figure 3 B, Supplemental Figure S1.4**).

##### K-means clustering

Once the optimal number of clusters is determined for both positive and negative slope segments using the **Diagnostic Mode** of STADIA, the user inputs these *k*-values, and *k*-means clustering is performed in the **Automated Mode** on the positive and negative slope segments separately. Segments from the simulation data were clustered using *k*=3 for each of the positive and negative slope groups. Similarly, the experimental data (either with or without CLASP2γ) was clustered using *k*=3 for positive slope segments and *k*=2 for negative slope segments (as discussed in the Results, *k*=2 for negative slope segments was appropriate for these datasets because the full depolymerizations were not captured for technical reasons). The final clustering results were obtained from using 500 random starts (again, multiple trials must be performed to determine optimal centroid locations as *k*-means is not guaranteed to converge to a global optimum); centroid locations that attained the lowest sum of squared distances between the centroids and each point in their respective clusters were chosen for further analysis.

##### Phase/behavior bundling

After *k*-means clustering, the clustering results are applied to the original un-standardized and un-log-transformed segment data. The multiple clusters of the positive and negative slope segments are collected, along with the ‘flat stutters’ that were removed prior to clustering, and all are considered together in the 3-D space defined by the segment features (**Supplemental Figure S1.7**). Statistics such as average slopes, average time duration, and average height change are calculated for each cluster (slopes in **Figure 3 C,F**; slopes and time durations in **Supplemental Figure S1.8**). Clusters with similar average slopes are bundled together to form larger groups, which we refer to as ‘phase classes’ or ‘behavior classes’ (**Figure 3 I**). Clusters with slopes considerably less in magnitude (flatter) are grouped into the category of behaviors called ‘stutters’ (along with the ‘flat stutters’ not considered during the clustering process). The remaining clusters with segment slopes larger in magnitude (i.e., the higher positive valued and lower negative valued slopes) more closely resembled the classically understood growth and shortening phases.

At this point, every segment identified during the segmentation stage has been classified as belonging to one of the following DI phase/behavior classes: nucleation, growth, shortening, or stutter. Applying these phase class labels to each segment in the length-history plot is illustrated in **Figure 3 G,H** and **Figure 6**. Using the chronological ordering of phases, the analysis of phase transitions can now be performed.

#### Phase and Transition Analysis Stage

After classifying segments into phase/behavior classes as described above, classical methods of calculating DI metrics are adapted to account for stutters in addition to growth and shortening.

##### Phase/behavior class analysis

The following attributes of each phase class are calculated: percent time spent in each phase class 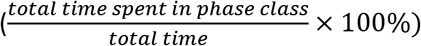, total number of segments (counts obtained from the piecewise linear approximation) for each phase class, and percent height change corresponding to each phase class (**Figure 5 A** and **Supplemental Figure S1.9**).

##### Transition analysis

Transition frequencies are calculated in a manner similar to what has been done classically. However, when considering stutters in addition to growth and shortening, there are additional transitions to quantify (**Figure 4**). In particular, it is necessary to determine whether catastrophes and rescues are (or are not) directly preceded by stutters. Catastrophes and rescues are identified as either abrupt (occurring without detectable stutters) or transitional (occurring via a stutter) (**Figures 5 D,E,F,G, 6, 7 D,E,F,G**). Additionally, our analysis quantifies interrupted growth (growth → stutter → growth) (**Figures 5 H, 6, 7 H,I**) and interrupted shortening (shortening → stutter → shortening) (**Figure 5 I**).

As mentioned above, MTs shorter than a user-defined threshold are considered to be in ‘nucleation’ phases (here the threshold used was 75 dimer-lengths). Transitions into or out of nucleation phases are not considered here because such MTs would be difficult to detect in experiments, and their behavior might be influenced by the seed.

In agreement with what has been done in classic DI analyses, frequencies of catastrophe and rescue are calculated as the ratio of the number of catastrophe or rescue events to the total time spent in growth or shortening, respectively (**Table 1; Supplemental Figure S1.10**). Similarly, interruptions are calculated as the ratio of the number of interrupted growths or interrupted shortenings to the total time spent in growth or shortening, respectively. Calculations can be done separately for abrupt and transitional types 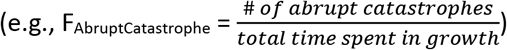 or collectively by simply adding the frequencies for each type together (F_cat_ = F_AbruptCatastrophe_ + F_TransitionalCatastrophe_) (**Supplemental Figure S1.10**).

Similar logic can be used to calculate the frequency of additional stutter-related transitions:

- (frequency of growth-to-stutter) = (frequency of interrupted growth) + (frequency of transitional catastrophe)
- (frequency of shortening-to-stutter) = (frequency of interrupted shortening) + (frequency of transitional rescue)

###### Guidance for users: Expectations for input data and effect of thresholds

STADIA is ideally intended for use on datasets with high temporal resolution, e.g., 2 or more frames per second. For lower resolution datasets, we suggest that STADIA will provide more systematic analysis than manual methods, but the resolution of the data itself will be a limiting factor in what conclusions can be supported.

We expect that the most common difficulty will be obtaining a total amount of data that is large enough for the clustering step of the classification stage. The clustering step performs better as the amount of data increases. To determine if one has a sufficient quantity of data, we suggest that users run STADIA on their entire dataset and on half of their dataset; if both cases yield similar results, this indicates that the user has a sufficient quantity of data. If one has only a small amount of data, STADIA can still be used to perform segmentation, detection of flat stutters with user-defined parameters, and classification with *k*=1 (one cluster each for positive and negative slope segments), but calculation of resulting DI metrics should be treated with caution.

It is important for users to be aware that the values of inputted thresholds will affect the numerical values of results of STADIA (as well as any other DI analysis method, e.g., (David J. Odde, Buettner, and Cassimeris 1996; Smal et al. 2010; Matov et al. 2010; Gierke, Kumar, and Wittmann 2010; Prahl et al. 2014; Guo et al. 2018)). For examples of the effects of changing these values, see the analyses with varying values of the Minimum Segment Duration and Maximum Error Tolerance in STADIA and the data acquisition rate of the length-history data as shown in **Supplemental Sections 2** and **3**. We recommend that users try at least a few different values of thresholds to test the strength of any conclusions they draw. In articles using STADIA, users should report the values of the input parameters that they use in STADIA, in addition to reporting the resolution of their measurements, quantity of data, and values of any other relevant quantities.

At the segmentation stage, users should examine the piecewise linear approximation to ensure that the approximation is not overfitting or underfitting the raw data. The user’s choices for the values of the Minimum Segment Duration and Maximum Error Tolerance determine how closely the piecewise linear approximation will fit the raw length-history data. When choosing the values of these thresholds, the user should take into account the resolution and noise level of their data as well as the timescale of the dynamics that the user wishes to study. For example, there are small-amplitude stochastic fluctuations that occur within growth, shortening, and stutter segments; if the user is studying phases at a similar scale to what we study in this article, which is a larger scale than the small-amplitude fluctuations, then the Minimum Segment Duration and Maximum Error Tolerance should not be so small as to pick up these fluctuations.

Note that for certain combinations of the Minimum Segment Duration and Maximum Error Tolerance, STADIA will produce ‘irreconcilable errors.’ These errors occur because it not always possible to satisfy both the Minimum Segment Duration and the Maximum Error Tolerance. In such cases, STADIA produces ‘irreconcilable errors’ and outputs a warning to the user for each error. Such errors are most likely to occur if the user has chosen a long Minimum Segment Duration with a small Maximum Error Tolerance. The specific values of Minimum Segment Duration and Maximum Error Tolerance that result in irreconcilable errors will depend on the particular dataset being analyzed. If such errors occur, the user should either change the parameter values or recognize that some segments of the piecewise linear approximation will not meet the input criteria. One useful approach to determining the significance of such errors is to examine the labeled length-history plots (e.g., **Figure 3 G,H**).

If the user is aiming to identify one set of input parameter values or a small number of parameter sets that are ideal for their particular dataset, then we recommend that the user choose input parameters values that minimize the number of irreconcilable errors. Our parameter sweep analysis (**Supplemental Material Sections 2 and 3**) indicates that the number of irreconcilable errors is more sensitive to the Maximum Error Tolerance than to the Minimum Segment Duration. For our *in silico* dataset, the percentage of segments that have irreconcilable errors has a local minimum at Maximum Error Tolerance = 20 dimer-lengths. The percentage of segments that have irreconcilable errors is also low for Maximum Error Tolerance > 40 dimer-lengths and is very high for Maximum Error Tolerance < 15 dimer-lengths. If the user is performing a parameter sweep with a large range of parameter values (similar to the parameter sweep done in **Supplemental Material Sections 2 and 3**), then the user should be aware that some parameter combinations may result in a large number of irreconcilable errors.

For further instructions on how to use the STADIA MATLAB code we refer readers to (Patel et al. 2020). The code as well as updated tutorials are available in our repository on GitHub (https://github.com/GoodsonLab/STADIA/). Note that the input parameter called the “Minimum Segment Duration” here was referred to as the “minimum time step” in (Patel et al. 2020).

### DATA ACQUISITION: IN SILICO MICROTUBULE EXPERIMENTS

This section outlines the details regarding the acquisition of the dimer-scale simulation MT data, analyzed in the main text Results and in **Supplemental Sections 1, 2, and 3**, including information about both the model and the parameters used.

#### The computational model: Stochastic model for simulating 13-protofilament (13-PF) MTs

The computational MT model used in this paper to generate the *in silico* length-history data is an updated version of the detailed, stochastic 13-PF MT model published in (Margolin et al. 2012) and utilized in (Margolin, Goodson, and Alber 2011; Gupta et al. 2013; Li et al. 2014; Duan et al. 2017; Jonasson et al. 2020; Mauro, Jonasson, and Goodson 2019). The model tracks the state of individual subunits (representing tubulin dimers bound to either GTP or GDP) in the entire 13-PF MT structure. The events that occur in the model are attachment/detachment of subunits to/from a PF, formation/breaking of lateral bonds between subunits in neighboring PFs, and hydrolysis of GTP-subunits to GDP-subunits. The values of the biochemical kinetic rate constants for each type of event are user inputs and depend on the state (GTP-bound or GDP-bound) of the subunits involved in the event. To carry out the simulation, the event that occurs at each step and the times between events are sampled using the Gillespie algorithm (Gillespie 1976, 1977), which is a kinetic Monte Carlo algorithm. At each event, the simulation outputs the time of the event and the length of the MT. The DI behavior, including stutters, and the values of DI parameters are emergent properties that arise as a consequence of the subunit-scale events. This feature is in contrast to two-state growth-shortening DI models, where the four traditional DI parameters are inputs (e.g., (Verde et al. 1992; Dogterom and Leibler 1993).

A key difference between the previous versions of our 13-PF MT computational model and the current implementation is strict adherence to the assumption that only one of the many possible biochemical events occurs at a time. The previous detailed level 13-PF MT model approximated hydrolysis events by allowing several subunits to hydrolyze simultaneously after one of the other four reaction events (lateral bonding/breaking or subunit gain/loss) have occurred. Hydrolysis events are now considered as a possible event in the same way that the other events are handled. This modification resulted in very little change in macro-level behavior of *in silico* MTs, but the ability to output dedicated observations of each dimer-level event provides a more accurate representation of MT biochemistry. The overall result of the simulation is *in silico* MTs that exhibit macro-level DI behaviors in agreement with those observed previously (Margolin et al. 2012).

#### Simulation setup and parameters

The dimer-scale kinetic parameters used in this study to simulate a 13-protofilament MT using the model described above were tuned in (Margolin et al. 2012) based on *in vitro* DI measurements from (Walker et al. 1988); a detailed list of parameters can be found in **Supplemental Table S1.2**. For the purposes of this analysis, a single MT was simulated at a constant [free tubulin] of 10 μM for 10 hours of simulation time. For the kinetic parameters and tubulin concentration used here, approximately 1650 subunit-scale reaction events occurred per second. To generate the length-history data passed into STADIA, we used either the max PF length (i.e., the length of the longest of the 13 protofilaments) or the mean PF length (i.e., the mean of the 13 protofilament lengths) as the length of the MT. Comparisons of results using the mean or max PF length are shown in **Supplemental Figures S1.4, S1.5, S1.6, S1.9, and S1.10**. The clustering profiles in **Supplemental Figure S1.4** show better agreement with the *in vitro* data used here when using the max PF length instead of the mean PF length. Thus, all the *in silico* results are presented for the max PF length unless otherwise indicated. Each dimer has a length of 8 nm. The max and mean PF lengths are both reported in units of dimer-lengths; this is not the same as the number of dimers in the MT, which would be 13 times the mean PF length.

### DATA ACQUISITION: *IN VITRO* MICROTUBULE EXPERIMENTS

This section outlines the details regarding capture of experimental MT data including conditions for a control group (tubulin + EB1) and a group with MTBPs (tubulin + EB1 + CLASP2γ).

#### Protein preparation

His-CLASP2γ and His-EB1 were purified as previously described (Zanic et al. 2013; Lawrence et al. 2018). Bovine brain tubulin was purified using the high-molarity method (Castoldi and Popov 2003). Tubulin was labeled with TAMRA and Alexa Fluor 488 (Invitrogen) according to the standard protocols, as previously described (Hyman et al. 1991).

#### TIRF microscopy

Imaging was performed using a Nikon Eclipse Ti microscope with a 100×/1.49 n.a. TIRF objective, NEO sCMOS (complementary metal–oxide–semiconductor) camera; 488- and 561-solid-state lasers (Nikon Lu-NA); Finger Lakes Instruments HS-625 high speed emission filter wheel; and standard filter sets. An objective heater was used to maintain the sample at 35°C. Microscope chambers were constructed as previously described (Gell et al. 2010). In brief, 22 × 22 mm and 18 × 18 mm silanized coverslips were separated by strips of Parafilm to create a narrow channel for the exchange of solution (Gell et al. 2010). Images were acquired using NIS-Elements (Nikon).

#### Dynamic MT Assay

GMPCPP-stabilized MTs were prepared according to standard protocols (Hyman et al. 1992; Gell et al. 2010). Dynamic MT extensions were polymerized from surface-immobilized GMPCPP-stabilized templates as described previously (Gell et al. 2010). The imaging buffer consisted of BRB80 supplemented with 40 mM glucose, 40 μg/ml glucose oxidase, 16 μg/ml catalase, 0.5 mg/ml casein, 100 mM KCl, 10 mM DTT, and 0.1% methylcellulose. The imaging buffer containing 1 mM GTP and purified proteins was introduced into the imaging chamber. Dynamic MTs were grown with 12 μM Alexa 488-labeled tubulin and 200 nM EB1 with or without 400 nM CLASP2γ and imaged at 2 fps using a 100× objective and an Andor Neo camera (pixel size of 70 nm). Alexa-488-labeled tubulin was used at a ratio of 23% of the total tubulin.

#### *In vitro* MT length-history data

Length-history data for *in vitro* MTs was obtained from 30 minute-long experiments using both a control group and a group with the stabilizing MTBP, CLASP2γ. Kymographs of dynamic microtubules (examples in **Figure 6**) were generated using the KymographClear macro for ImageJ, and the dynamic MT tip positions as a function of time were determined in KymographClear, using a thresholding-based, edge-detection method that can trace the microtubule tip position in kymographs with subpixel accuracy (Mangeol, Prevo, and Peterman 2016). A subset of this experimental dataset was previously published in (Lawrence et al. 2018). The control group data was acquired from 68 MT seeds, from which 776 individual traces were observed. The group with CLASP2γ was acquired from 29 MT seeds, from which 85 individual traces were observed. After applying the stitching preprocessing step during the STADIA segmentation stage, the control group and the group with CLASP2γ each generated a single stream time series representing length-history data with total time duration over 21 hours and 3.5 hours, respectively. The collective consideration of all experimental data samples together meets the needs of the machine learning requirements for reliable clustering results (i.e., the lifetime of a single MT alone would not be a sufficient amount of data for *k*-means clustering). The *in vitro* MT lengths were measured in units of nm, and then divided by 8 nm per dimer-length to convert to units of dimer-lengths.

#### Software and data availability

STADIA software and tutorials can be downloaded from GitHub (https://github.com/GoodsonLab/STADIA/). Data analyzed in this paper are available from the authors upon request.

## ACKNOWLEDGEMENTS

This work was supported by NSF grants MCB-1244593 to HVG and MSA, MCB-1817966 to HVG, and MCB-1817632 to EMJ, and NIH grants R35GM119552 to MZ and IBSTO training grant T32CA119925 to EJL. MZ also acknowledges the Searle Scholars Program. Portions of the work were also supported by funding from the University of Massachusetts Amherst (AJM), NSF-GFRP DGE-1313583 (KSM), and a fellowship from the Dolores Zohrab Liebmann Fund (SMM). We thank the members of the Goodson laboratory and the Chicago Cytoskeleton community for their insightful discussions.

