## supplement for "Quantification of Microtubule Stutters: Dynamic Instability Behaviors that are Strongly Associated with Catastrophe"

#### Supplement Table of Contents


| Section | Topic | Pages |
| --- | --- | --- |
| **Section 1** | Main supplemental figures and tables | pp. 2-14 |
| **Section 2** | STADIA parameter sweep analysis | pp. 15-36 |
| **Section 3** | Data acquisition rate sensitivity analysis | pp. 37-57 |
| **Section 4** | Negative control: two-state model | pp. 58-64 |


**Note**: The simulation data used in the main text and in **Supplemental Sections 1, 2, and 3** was generated from the stochastic dimer-scale 13-PF model described in the Methods. In contrast, **Supplemental Section 4** presents and analyzes simulation data from a different model, which was designed to only have two states: growth and shortening.

**MOVIES:** Separate supplemental files (**Movies 1-4**) contain movies from the *in vitro* control data corresponding to the kymographs in **Figure 6** of the main text. Note that the images were acquired at 2 fps, while the movies are presented at 7 fps, meaning that the movies are presented at 3.5 x the time labeled on the kymographs and length-history plots.

**Movie 1**: Corresponds to the kymograph in **Figure 6A** – Example of **Abrupt Catastrophe**
**Movie 2**: Corresponds to the kymograph in **Figure 6B** – Example of **Abrupt Catastrophe**
**Movie 3**: Corresponds to the kymograph in **Figure 6C** – Example of **Transitional Catastrophe**
**Movie 4**: Corresponds to the kymograph in **Figure 6D** – Example of **Transitional Catastrophe**

Each kymograph and corresponding movie also contains at least one example of **Interrupted Growth**, as indicated in **Figure 6**.

#### **Supplemental Section 1:**
Main supplemental figures and tables

**Table of Contents**


**Supplemental Figures S1.1-S1.10** Pages 3-12

**Supplemental Tables S1.1-S1.2** Pages 13-14

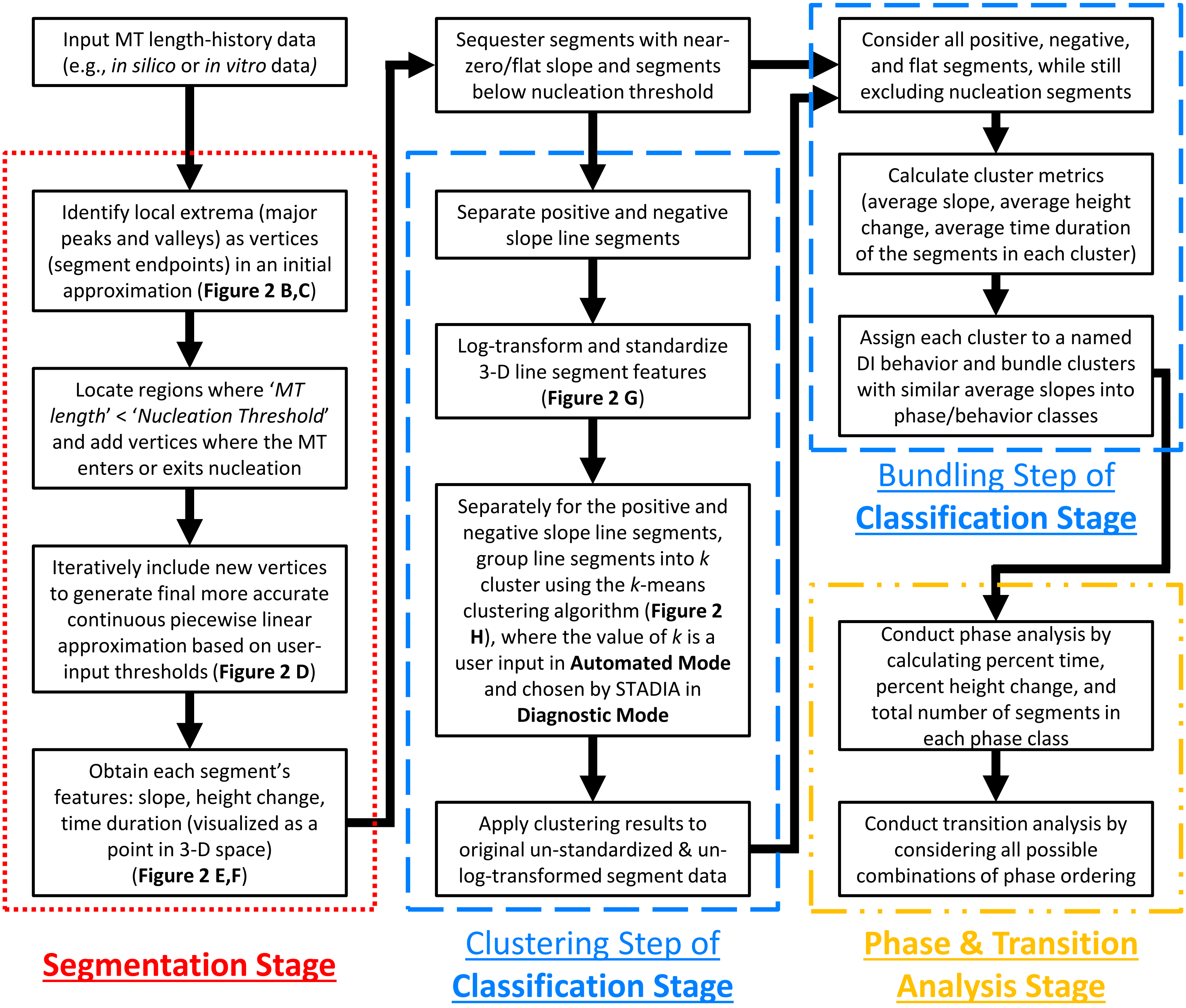


**Figure S1.1. Workflow diagram outlining main steps in each stage of STADIA.** Note that **Automated Mode** performs all the steps shown in the workflow diagram. **Diagnostic Mode** performs the steps through the end of the clustering step of the classification stage. Running STADIA in **Diagnostic Mode** before **Automated Mode** provides information to aid the user in choosing the optimal number of clusters (*k*-values) to input into **Automated Mode**. Additional details regarding the technical details of STADIA can be found in the Methods section of the main text.

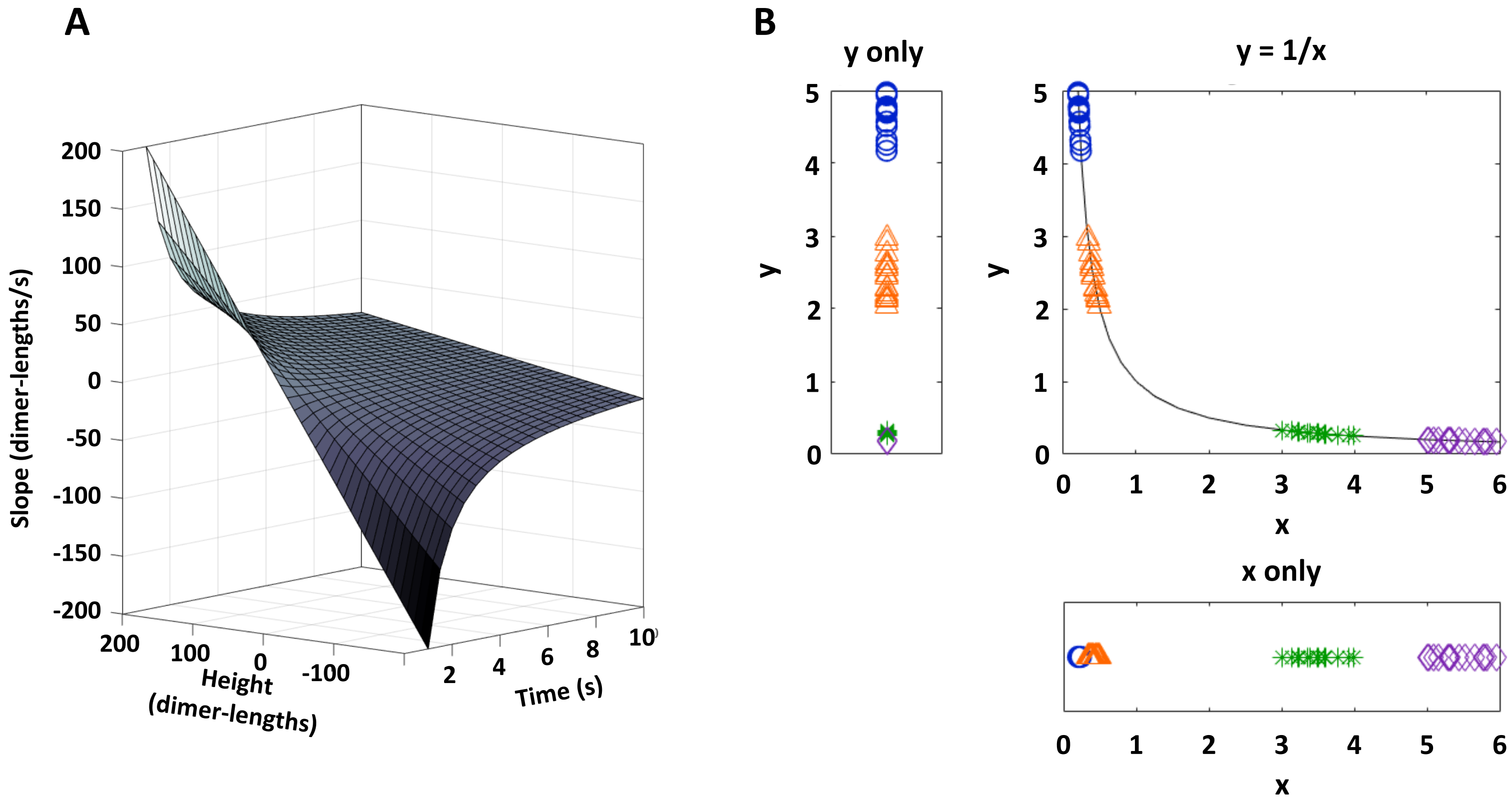


**Figure S1.2. Slope=height/time surface in 3-D space, and analogous example in 2-D. (A)** Data points representing each line segment reside on this Z(X,Y) = Y/X manifold (surface), where Z = slope, X = time, and Y = height. See the subsection "Justification for classification feature space" in the main text Methods section regarding the use of all three variables in the Classification Stage. **(B)** Justification for using height, time, and slope is demonstrated using a parallel example in two dimensions: a dataset containing four clusters (groups) of points that fall on the curve y = 1/x. Plotting only y (left of y=1/x plot) or only x (below y=1/x plot) creates the appearance that this dataset contains only three clusters of points. In contrast, when the data are plotted in two dimensions (y=1/x plot), the data are separated sufficiently to reveal that the dataset actually contains four clusters of points. For similar reasons, we need to consider all three variables in our line segment data to properly identify the clusters in our dataset.

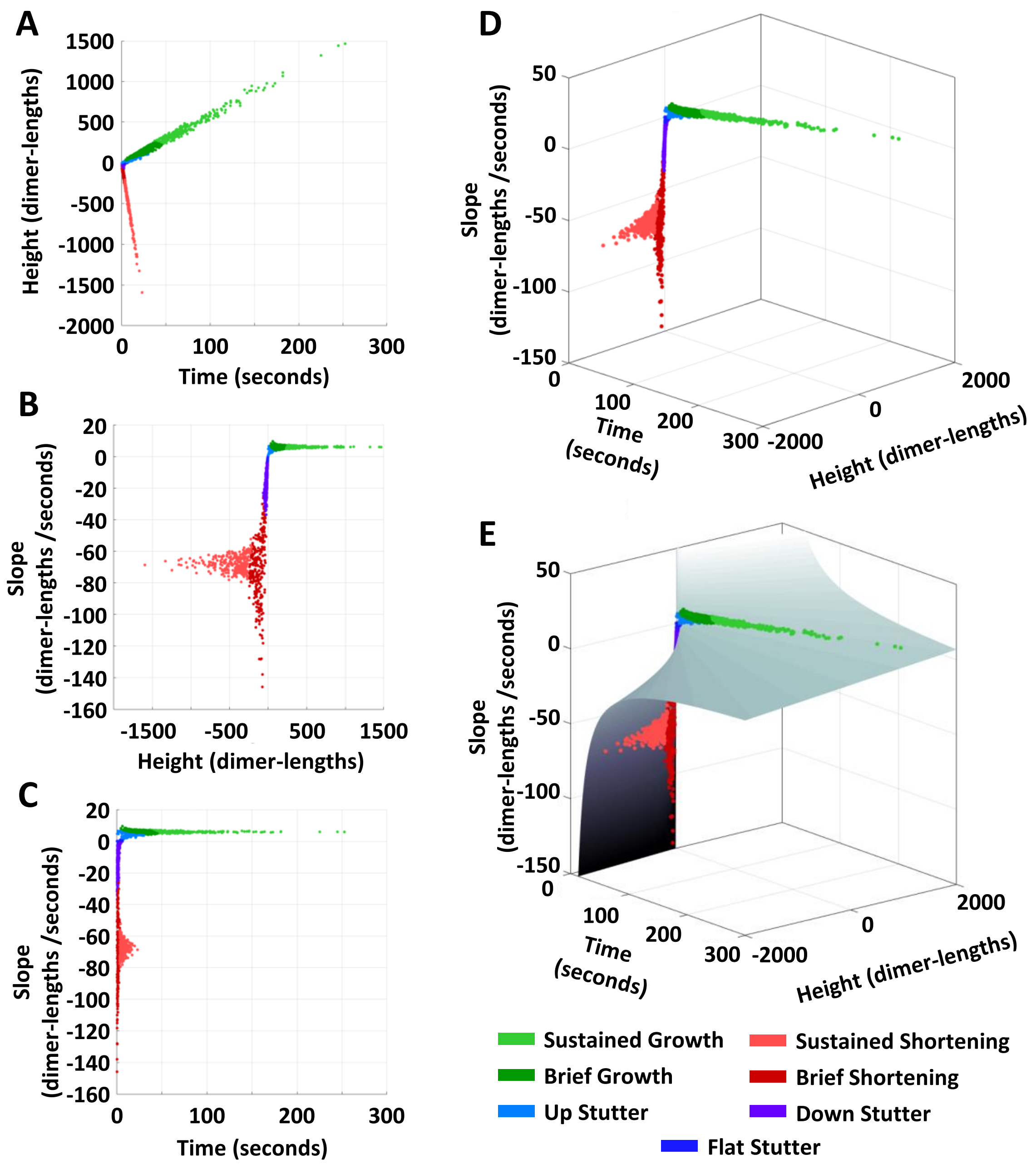


**Figure S1.3. The segment features (slope, height change, and time duration) for segments identified from the piecewise linear approximation of the dimer-scale *in silico* length-history data.** Each point corresponds to one line segment from the length-history approximation and is colored according to the cluster identified by STADIA. **(A-C)** Multiple perspectives of the segment feature data represented in two dimensions (Height and Time (A), Slope and Height (B), Slope and Time (C)) demonstrate the lack of separability between points when only two dimensions are considered (similar to the example in **Supplemental Figure S1.2 B**). **(D)** Final clustering profile of all un-standardized and un-log-transformed segment data following the Classification Stage, provided to help visualize the 3-D data. **(E)** An illustration of how the segment points lie on the Z=Y/X manifold described in **Supplemental Figure S1.2 A**.

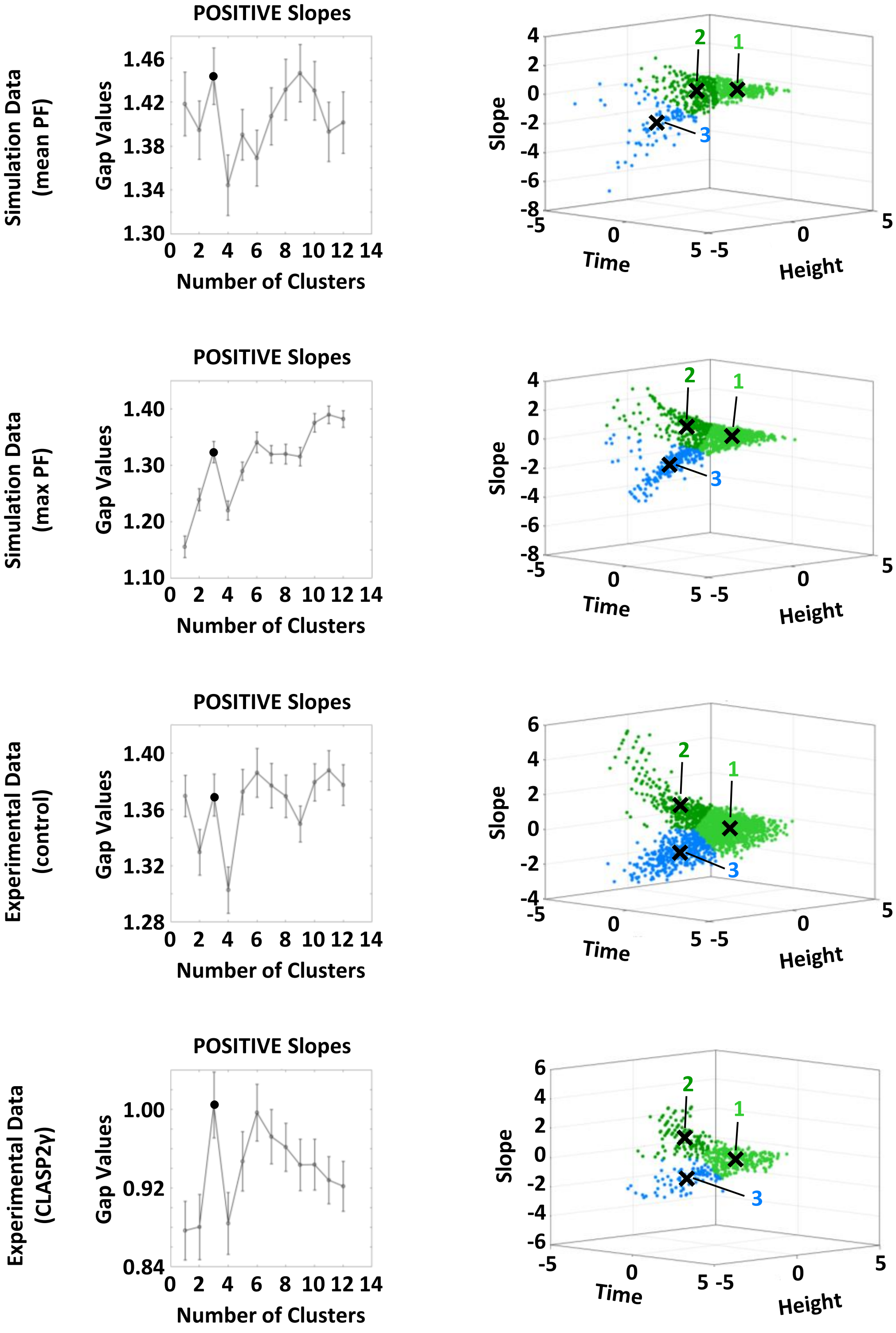


**Figure S1.4. Gap statistic plots (left column) and corresponding clustering profiles (right column) for positive slope segments in each log-transformed and standardized dataset**. When using the gap statistic to suggest the best number of clusters to use in *k*-means clustering, a rule of thumb is to use the first *k*-value where the gap statistic plot shows a local maximum. In practice here, we expect the number of clusters to be greater than 1, because the 3-D data structure (right column) shows multiple appendages. Thus, for the cases of the dimer-scale simulation data using the mean PF length and the control experimental data, the local maximum at $k$=1 is rejected. Taking this into consideration, all datasets indicate that the gap statistic attains the first local maximum greater than one at $k$=3. Thus, for all positive slope segment data, *k*-means clustering is performed by separating the data into 3 clusters. Furthermore, the clustering profile of the simulation data using the max PF length, rather than the mean PF length, more closely resembles the clustering profile of the experimental (note that for the mean PF data, there are fewer rapid, short duration segments in cluster 2). Therefore, we chose to use the max PF data instead of the mean PF data for presenting the STADIA results in the main text.

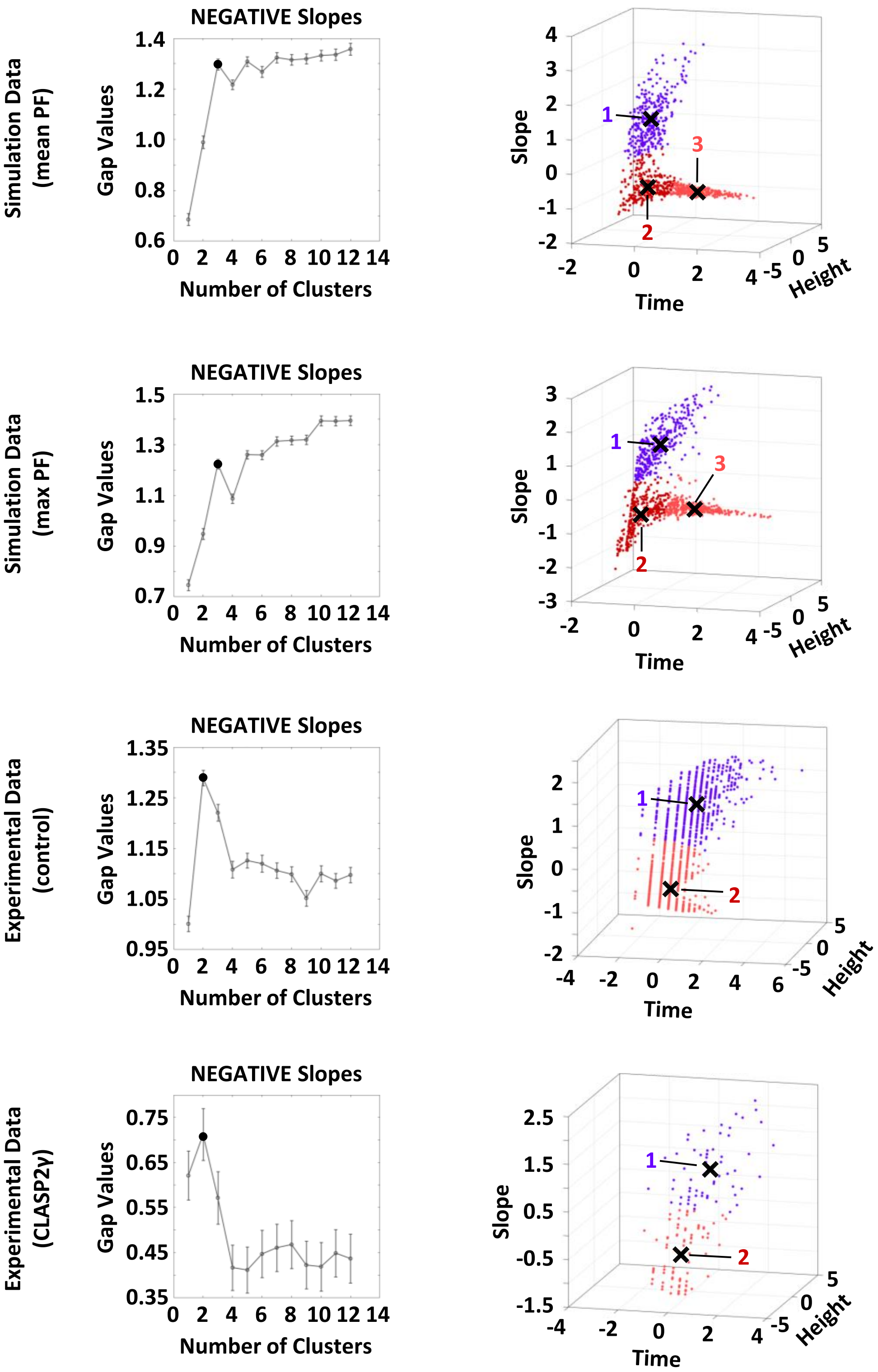


**Figure S1.5. Gap statistic plots (left column) and corresponding clustering profiles (right column) for negative slope segments in each log-transformed and standardized dataset.** When using the gap statistic to suggest the best number of clusters to use in *k*-means clustering, a rule of thumb is to use the first *k*-value where the gap statistic plot shows a local maximum. The two simulation datasets indicate that the gap statistic attains the first local maximum at $k$=3, whereas the experimental datasets indicate $k$=2. We attribute this difference to the fact that in these experimental datasets, only the beginnings of depolymerization phases were captured, thus omitting long time duration shortening segments from the dataset. Thus, for negative slope segments, we performed *k*-means clustering separating the dimer-scale simulation data into 3 clusters and the experimental data into 2 clusters.

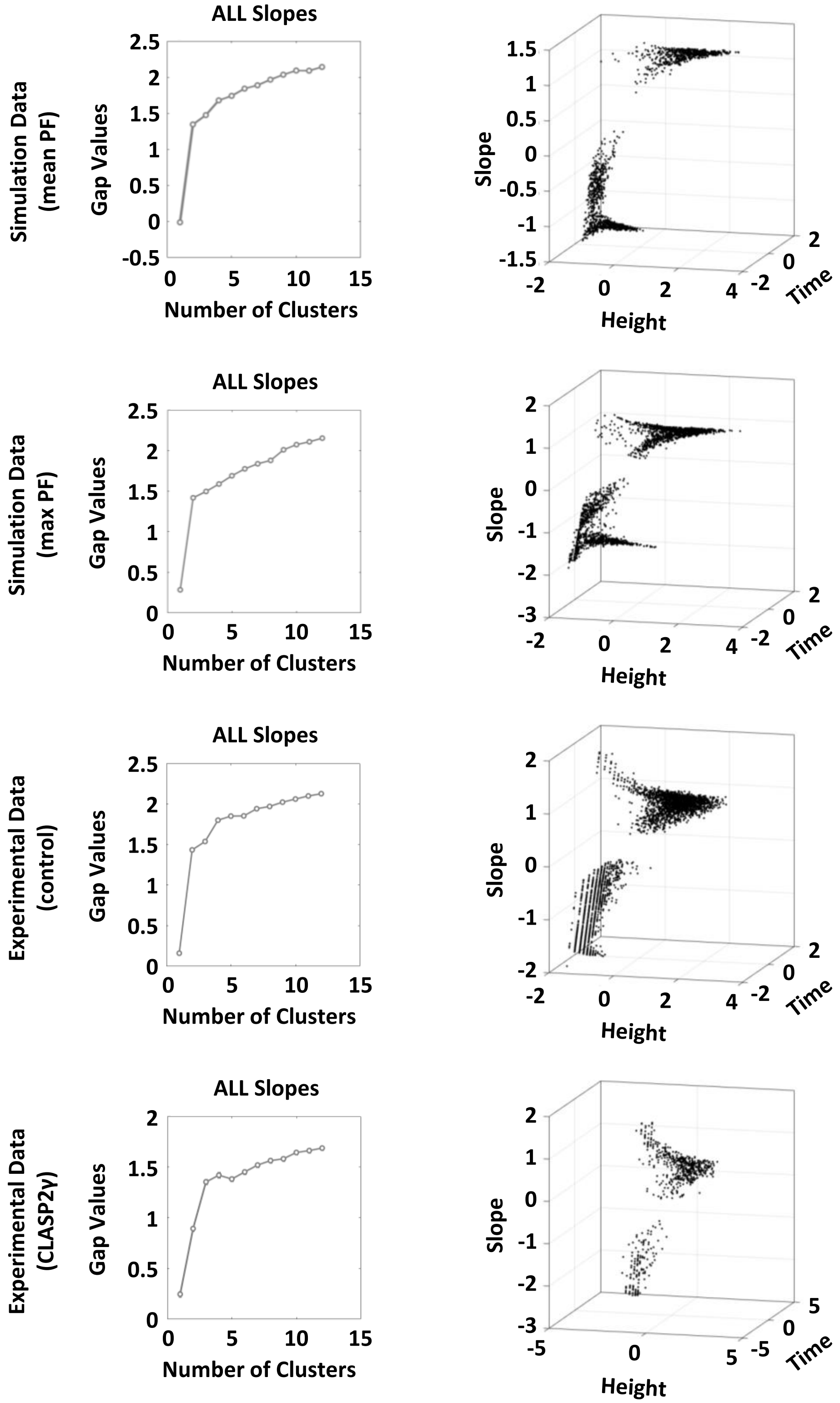


**Figure S1.6. Gap statistic plots (left column) and segment feature plots (right column) for an analysis where all slope segments in each dataset were considered together (excluding flat segments), not separated into positive and negative slopes as in Supplemental Figures S1.4 and S1.5**. The gap statistic plots are generally increasing with no clear local maxima, indicating that the initial dataset was too complex for effective calculation of the gap statistic and that we needed to subdivide it before further analysis. For this reason, the data are not color-coded as in the previous two figures. Note that these plots are simply for demonstrating that consideration of all segments together is not conclusive, thus providing justification for analyzing positive and negative slope segments separately.

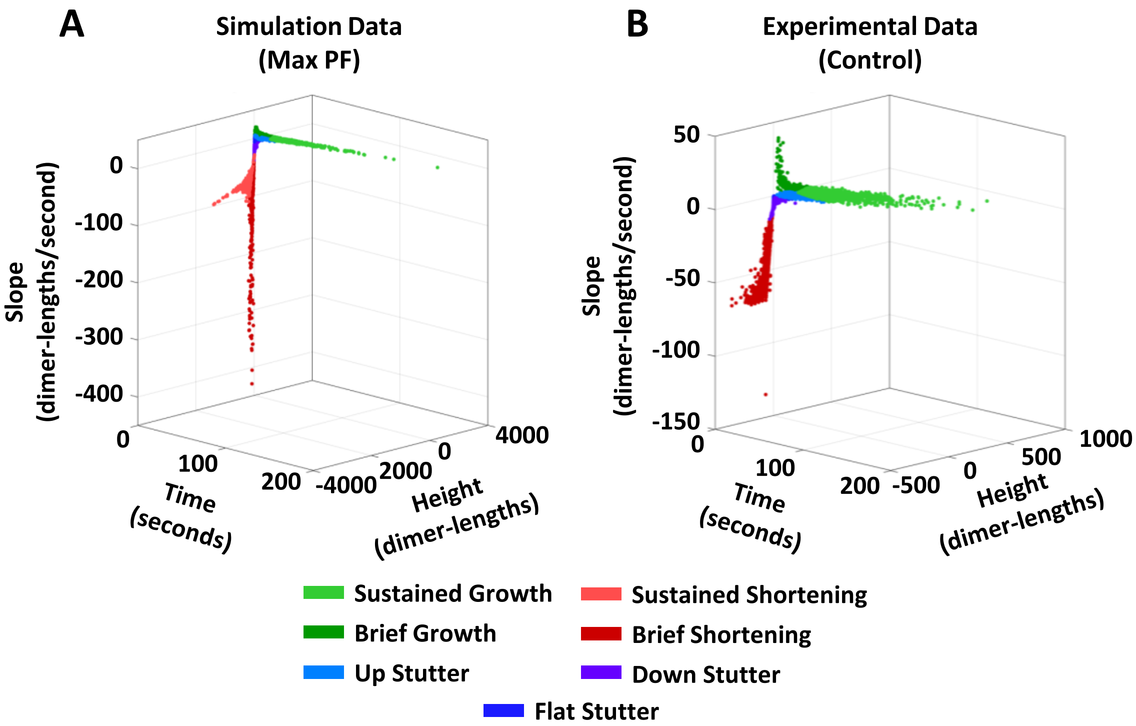


**Figure S1.7. Clustering profiles for ALL segments of *in silico* and *in vitro* data.** Following separate classification of the positive and negative slope segments (see **Supplemental Figures S1.4** and **S1.5**), cluster assignments were applied to the original un-log-transformed and un-standardized segment data and plotted for full view of clustered line segment data in 3-D space. Note that the classification step has already taken place, and these figures are simply for visualizing how the clusters exist in relation to each other in 3-D space.

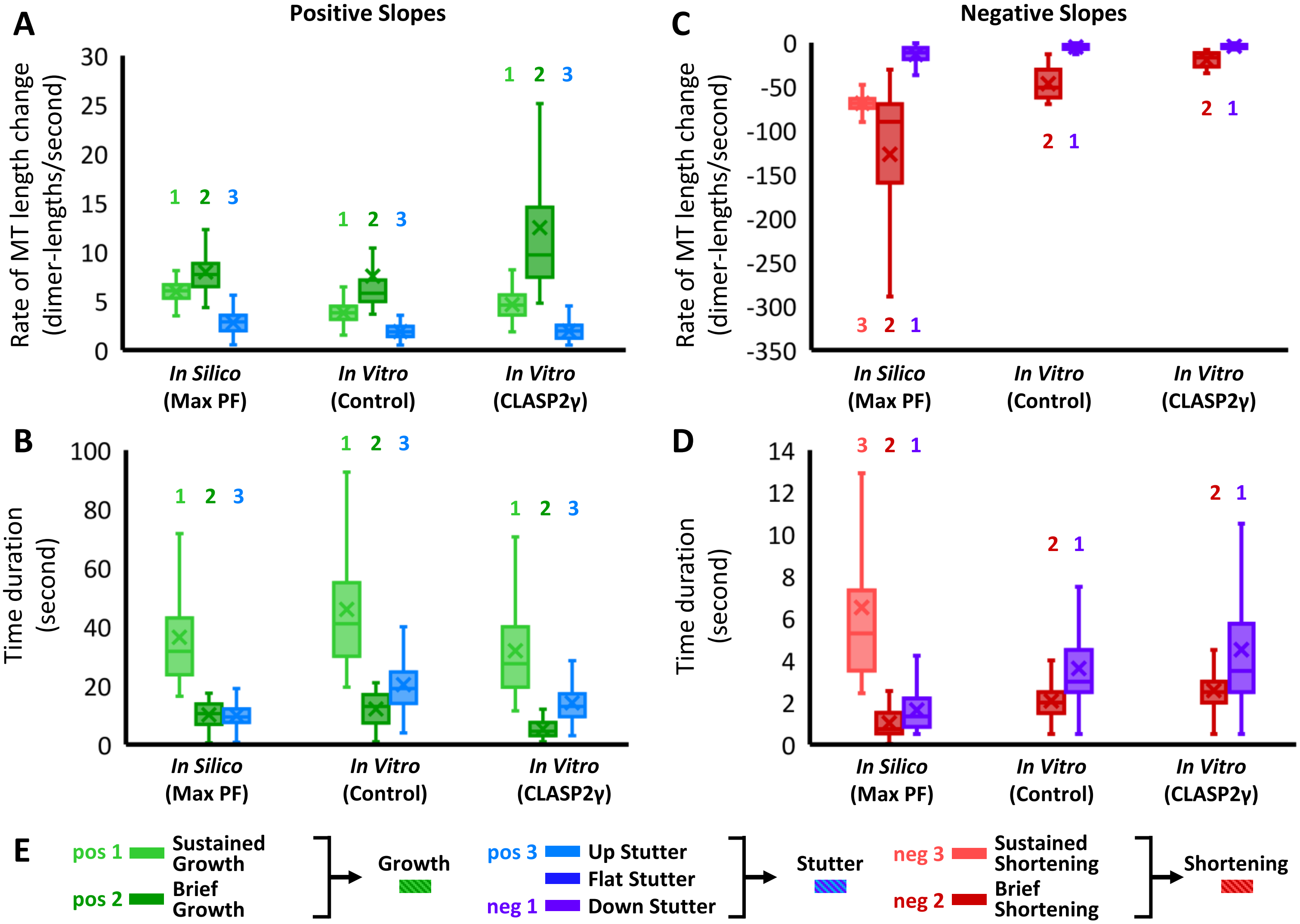


**Figure S1.8. Justification for "phase/behavior bundling," i.e., grouping together multiple clusters as identified by STADIA into growth, shortening, and stutter classes.** Left column: positive slope segments. Right column: negative slope segments. The *in silico* max PF and *in vitro* control data in **A** and **C** are the same as the data presented in **Figure 3 C** and **F**, respectively. The box and whisker plots cover the four quartiles of each cluster of segments, and the 'X' marks the mean value in the plots. Outliers were excluded from these plots using the default definition in MATLAB (i.e., any value that is more than 1.5 times the interquartile range away from the bottom or top of the box is considered an outlier). **(A)** Growth rates for the positive slope segment clusters. Note that in each dataset, clusters 1 and 2 (light and dark green) have average growth rates relatively large in magnitude compared to cluster 3 (light blue). **(B)** Time durations for the positive slope segment clusters show that cluster 1 (light green) represents longer, more sustained periods of consistent behavior than clusters 2 and 3 (dark green and light blue) for all datasets. The combination the observations from (A) and (B) indicates that clusters 1 and 2 (green) differ primarily by time duration and justifies grouping them into Growth. In contrast, cluster 3 (light blue) has much shallower slopes than clusters 1 and 2 (green), justifying the classification of cluster 3 as Stutter. **(C)** Shortening rates for the negative slope segment clusters. Note that in each dataset, the shortening rates of cluster 2 (dark red) and, when present, cluster 3 (light red) were on average relatively large in magnitude compared to cluster 1 (purple). **(D)** Time durations for the negative slope segment clusters from the simulation data indicates that clusters 2 and 3 (dark and light red) differ in time duration. The combination of the observations from (C) and (D) indicates that clusters 2 and 3 (red) differ primarily by time duration and justifies grouping them into Shortening. In contrast, the cluster 1 (purple) has shallower slopes than clusters 2 and 3 (red), justifying the classification of cluster 1 as Stutter. Note that since the experimental data did not capture most of the shortening behavior, analysis of longer time duration shortening segments was not possible for the *in vitro* datasets. **(E)** Bundling clusters together into larger phase/behavior classes based on the observations described above. The 'Brief' and 'Sustained' clusters within the Growth and Shortening classes are characterized by their time durations. The 'Up', 'Flat', and 'Down' clusters in the Stutters category are characterized by the segment slopes being positive, near-zero, or negative respectively.

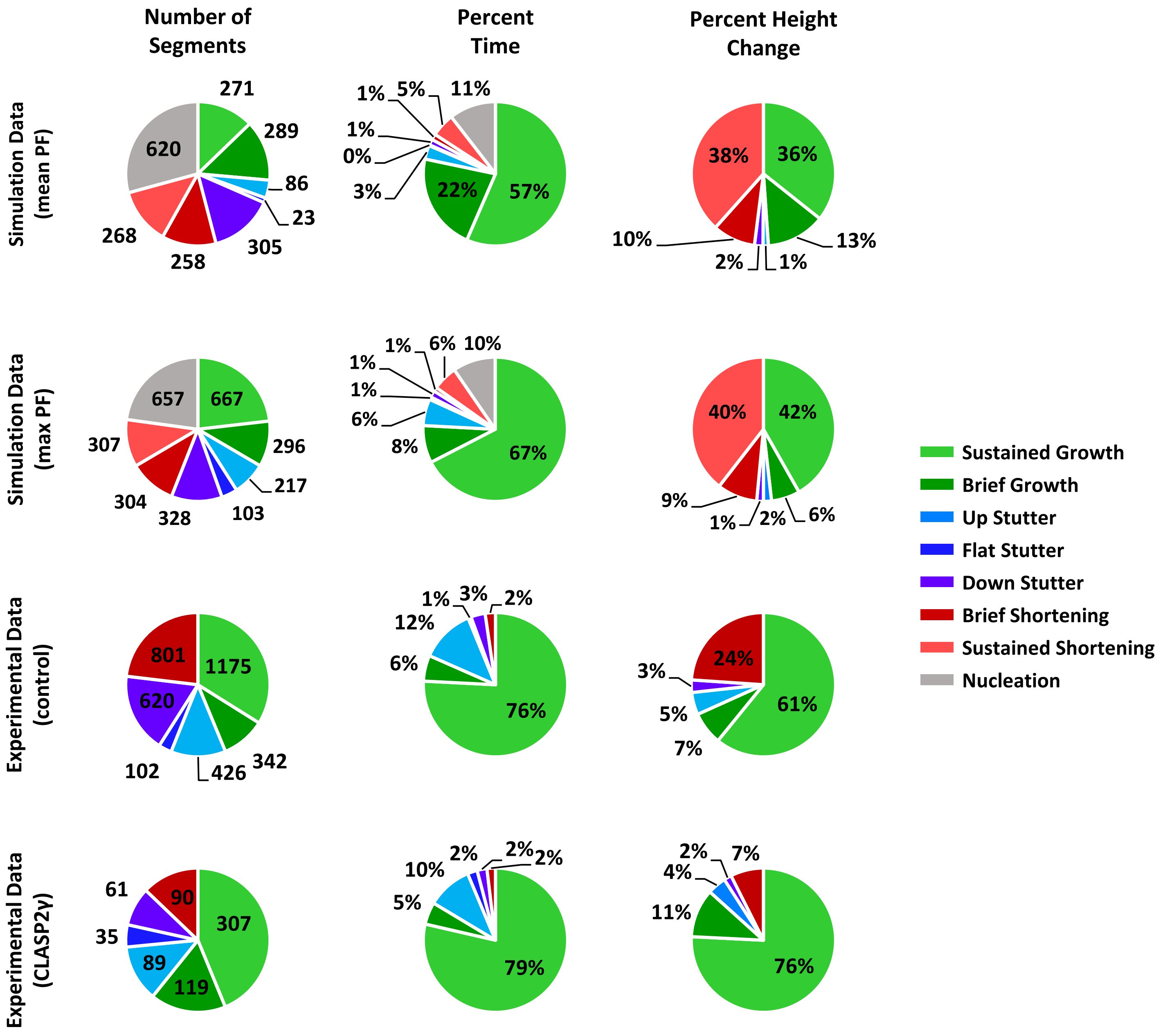


**Figure S1.9. Segment statistics for all *in silico* and *in vitro* datasets.** *Number of segments*, *percent time*, and *percent height change* for each cluster are recorded for each dataset. For the *in silico* datasets, where depolymerizations were fully captured, the breakdown of the number of segments, time duration, and height change are representative of the actual time the simulated MT spent in the various phases. As noted throughout the paper, the *in vitro* depolymerizations were not captured in their entirety, and so the number of negative slope segments, percent time, and percent height change attributed to negatively sloped MT behavior is largely underreported.

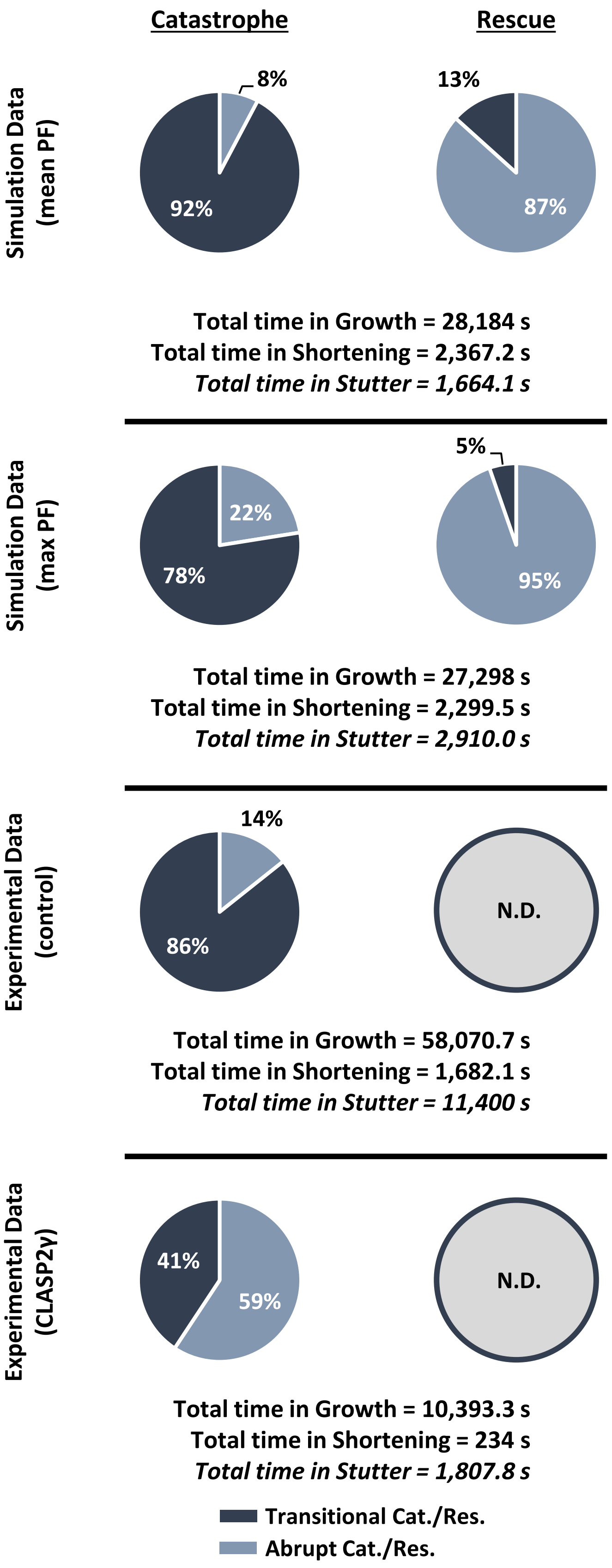


| Transition Statistics | | |
|---|---|---|
| | Count | Frequency ($s^{-1}$) |
| **Simulation Data (mean PF)** | | |
| Abrupt Catastrophe | 20 | 0.00071 |
| Transitional Catastrophe | 238 | 0.0084 |
| **Total Catastrophe** | **258** | **0.00911** |
| Abrupt Rescue | 39 | 0.0164 |
| Transitional Rescue | 6 | 0.0025 |
| **Total Rescue** | **45** | **0.0189** |
| Interrupted Growth | 75 | 0.0027 |
| Interrupted Shortening | 32 | 0.0135 |
| **Simulation Data (max PF)** | | |
| Abrupt Catastrophe | 67 | 0.0025 |
| Transitional Catastrophe | 231 | 0.0085 |
| **Total Catastrophe** | **298** | **0.0110** |
| Abrupt Rescue | 71 | 0.0307 |
| Transitional Rescue | 4 | 0.0017 |
| **Total Rescue** | **75** | **0.0324** |
| Interrupted Growth | 293 | 0.0107 |
| Interrupted Shortening | 48 | 0.0207 |
| **Experimental Data (control)** | | |
| Abrupt Catastrophe | 105 | 0.0018 |
| Transitional Catastrophe | 629 | 0.0108 |
| **Total Catastrophe** | **734** | **0.0126** |
| Abrupt Rescue | 18 | **N.D.** |
| Transitional Rescue | 0 | |
| **Total Rescue** | **18** | |
| Interrupted Growth | 211 | 0.0036 |
| Interrupted Shortening | 0 | **N.D.** |
| **Experimental Data (CLASP2γ)** | | |
| Abrupt Catastrophe | 51 | 0.0049 |
| Transitional Catastrophe | 35 | 0.0034 |
| **Total Catastrophe** | **86** | **0.0083** |
| Abrupt Rescue | 45 | **N.D.** |
| Transitional Rescue | 7 | |
| **Total Rescue** | **52** | |
| Interrupted Growth | 92 | 0.0088 |
| Interrupted Shortening | 1 | **N.D.** |

**Figure S1.10. Detailed transition statistics for each dataset.** Both *in silico* datasets as well as the control experimental dataset demonstrate that a significant majority of catastrophes occur via stutter (i.e., transitional catastrophe), while the CLASP2γ dataset shows a shift to MTs exhibiting abrupt catastrophes along with a decrease in total catastrophe frequency. The presence of CLASP2γ markedly reduces the frequency of transitional catastrophe and increases the frequency of interrupted growth (see also **Figure 7** and the Results section). Rescue data for *in silico* MTs indicate that most rescues occur abruptly. Note that frequencies of rescue and interrupted shortening were not determined (N.D.) for the *in vitro* data because depolymerizations were not captured in their entirety for the *in vitro* MTs.

| STADIA: User-defined Parameters | |
|---|---|
| Nucleation height threshold | 75 dimer-lengths |
| Minimum Segment Duration | 500 ms |
| Maximum Error Tolerance | 20 dimer-lengths |
| Maximum height change for near-zero slope segments | 3 dimer-lengths |
| Maximum slope magnitude for near-zero slope segments | 0.5 dimer-lengths/sec |
| Number of centroids for positive slope segments | 3 |
| Number of centroids for negative slope segments (*in silico* data) | 3 |
| Number of centroids for negative slope segments (*in vitro* data) | 2 |

| Classical Peak-Valley Analysis: User-defined Parameters | |
|---|---|
| Minimum peak height | 95 dimer-lengths |
| Minimum rescue length | 95 dimer-lengths |
| Minimum Prominence For Major Peaks | 20 dimer-lengths |
| Minimum Prominence For Minor Peaks | 0.1 dimer-lengths |
| Minimum Regression $R^2$ | 0.95 |

**Table S1.1. User-defined parameters for STADIA and classical Peak-Valley analysis.**
See **Supplemental Sections 2 & 3** for parameter sensitivity analysis.

| Stochastic Dimer-Scale 13-PF MT Model Parameters | |
|---|---|
| Number of protofilaments | 13 |
| Tubulin concentration | 10 µM |
| Simulation time | 10 hours |
| Seam shift | 1.5 dimer-lengths |
| Compete for tubulin | No |
| Hydrolysis rate | 0.7 dimers/sec |
| HalfMax | 200 |
| kgrowT | 250 |
| kgrowD | 250 |
| kshortT | 0.02 |
| kshortD | 20 |
| kbondTT | 100 |
| kbondTD | 100 |
| kbondDT | 100 |
| kbondDD | 100 |
| kbreakTT | 70 |
| kbreakTD | 90 |
| kbreakDT | 90 |
| kbreakDD | 400 |
| SkbondTT | 200 |
| SkbondTD | 200 |
| SkbondDT | 200 |
| SkbondDD | 200 |
| SkbreakTT | 140 |
| SkbreakTD | 180 |
| SkbreakDT | 180 |
| SkbreakDD | 800 |

**Table S1.2. Computational model parameters used to generate the dimer-scale simulation data.** Parameter values used are from Margolin et al. 2012. Please see Methods section for more information about the model.

### Supplemental Section 2:
#### STADIA parameter sweep analysis

**Overview:** The goal of the analyses presented here is to test the robustness of the main conclusions of the manuscript to variation in the values of the user-input STADIA segmentation parameters "Minimum Segment Duration" and "Maximum Error Tolerance". In the next section (3), we examine sensitivity to the temporal resolution of the inputted length-history data itself.

In brief, the conclusions being considered are as follows: **(1)** MTs exhibit more behaviors than just growth and shortening, with stutters being distinguishable behaviors that are prevalent throughout length-history data, **(2)** transitional catastrophes are more frequent than abrupt catastrophes, and **(3)** the anti-catastrophe factor CLASP2γ reduces catastrophe frequency by promoting stuttering MTs to return to growth. The table of contents below directs readers to the figures related to each of the above conclusions. In addition, the next page contains information to assist readers in understanding and interpreting the analyses performed.

**Table of Contents**

**Supplemental Section 2: Parameter sweep analysis of dimer-scale *in silico* and TIRF-imaged *in vitro* datasets**

**1.** Analysis of *in silico* data with varying segmentation parameters
- **a.** Gap statistic plots – **Figures S2.1** (positive slope segments), **S2.2** (negative slope segments)
  *Useful for conclusion (1)*
- **b.** Cluster profiles of positive and negative slope segment data – **Figure S2.3**
  *Useful for conclusion (1)*
- **c.** Labeled length-history plots – **Figure S2.4**
  *Useful for conclusions (1) and (2)*
- **d.** Frequencies of abrupt and transitional catastrophe and interrupted growth – **Figure S2.9**
  *Useful for conclusions (1) and (2)*

**2.** Analysis of *in vitro* (control) data with varying segmentation parameters
- **a.** Gap statistic plots – **Figures S2.5** (positive slope segments), **S2.6** (negative slope segments)
  *Useful for conclusion (1)*
- **b.** Cluster profiles of positive and negative slope segment data – **Figure S2.7**
  *Useful for conclusion (1)*
- **c.** Labeled length-history plots – **Figure S2.8**
  *Useful for conclusions (1) and (2)*
- **d.** Frequencies of abrupt and transitional catastrophe and interrupted growth – **Figure S2.10**
  *Useful for conclusions (1), (2), and (3)*

**3.** Analysis of *in vitro* (CLASP2γ) data with varying segmentation parameters
- **a.** Frequencies of abrupt and transitional catastrophe and interrupted growth – **Figure S2.11**
  *Useful for conclusion (3)*

**4.** Comparison of *in vitro* control and CLASP2γ data with varying segmentation parameters
- **a.** Frequencies of transitional catastrophe and interrupted growth – **Figure S2.12**
  *Useful for conclusion (3)*

**Note to Readers:** Each figure is accompanied by a legend as well as explanatory text that contains observations, interpretations, and in some cases additional discussion. For most of the figures, some of the accompanying text is on the page following the figure. Since parameter sweeps are repetitive by nature, we use similar language in the text accompanying many figures. Despite some apparent redundancy, we use this approach to make each figure independently interpretable.

#### **Introduction to Section 2,** STADIA parameter sweep analysis

STADIA has two parameters that directly affect segmentation of MT length-history data and are determined entirely by the user: 1) Minimum Segment Duration (i.e., how short can the time duration of a segment used to approximate MT length-history data be?), and 2) Maximum Error Tolerance (i.e., how close do data points need to be approximated by a corresponding line segment?). In this parameter sweep, we tested the effect of changing these parameters across a range of potential values by using STADIA with these varying parameters to (re)analyze the same datasets considered in the main text.

For the **full resolution**† *in silico* dataset, the parameter space considered is as follows: 1) Minimum Segment Duration = 0.3, 0.5, 1.0, 1.5, 2.0, 3.0 seconds, and 2) Maximum Error Tolerance = 5, 10, 15, 20, 25, 30, 35, 40, 63 tubulin dimers (i.e., 40-500 nm). The parameter space considered for the *in vitro* datasets is identical except that because the Data Acquisition Time Step was 0.5 seconds (i.e., 2 fps) for the *in vitro* data, these datasets were not analyzed using a Minimum Segment Duration of 0.3 seconds (Minimum Segment Duration must be ≥ Data Acquisition Time Step).

Throughout the range of values considered for each parameter, analysis was conducted for every possible combination (e.g., every Minimum Segment Duration is used with every Maximum Error Tolerance). Therefore, the layout of each figure is a grid where columns correspond to specific values of the Minimum Segment Duration and rows correspond to specific values of the Maximum Error Tolerance.

For each dataset (*in silico* and *in vitro* with/without CLASP), the parameter sweep analysis figures are presented in the order used in a regular STADIA analysis: the figures start with gap statistic plots followed by cluster profiles of the positive and negative slope segments, and then the labeled length-history plots that result from the earlier stages of the analysis. At the end of **Supplemental Section 2**, we provide a series of tables that summarize the effects of parameter variation on the frequencies of abrupt catastrophe, transitional catastrophe, and interrupted growth. **Readers more interested in the transition frequency results than the process may want to go directly to the final figures in this section** (**Figures S2.9-S2.12**, pages 30-36).

*A note for users of the STADIA code:* it is not always possible to satisfy both the Minimum Segment Duration and the Maximum Error Tolerance input parameters. Such ‘irreconcilable errors’ are especially likely in cases where user has chosen a long Minimum Segment Duration with a small Maximum Error Tolerance. For additional information about irreconcilable errors and how to interpret them, please see the main text Methods section.

† The **full resolution** *in silico* data has dimer-scale spatial resolution, and time resolution of one output per dimer-scale biochemical event (see Methods).

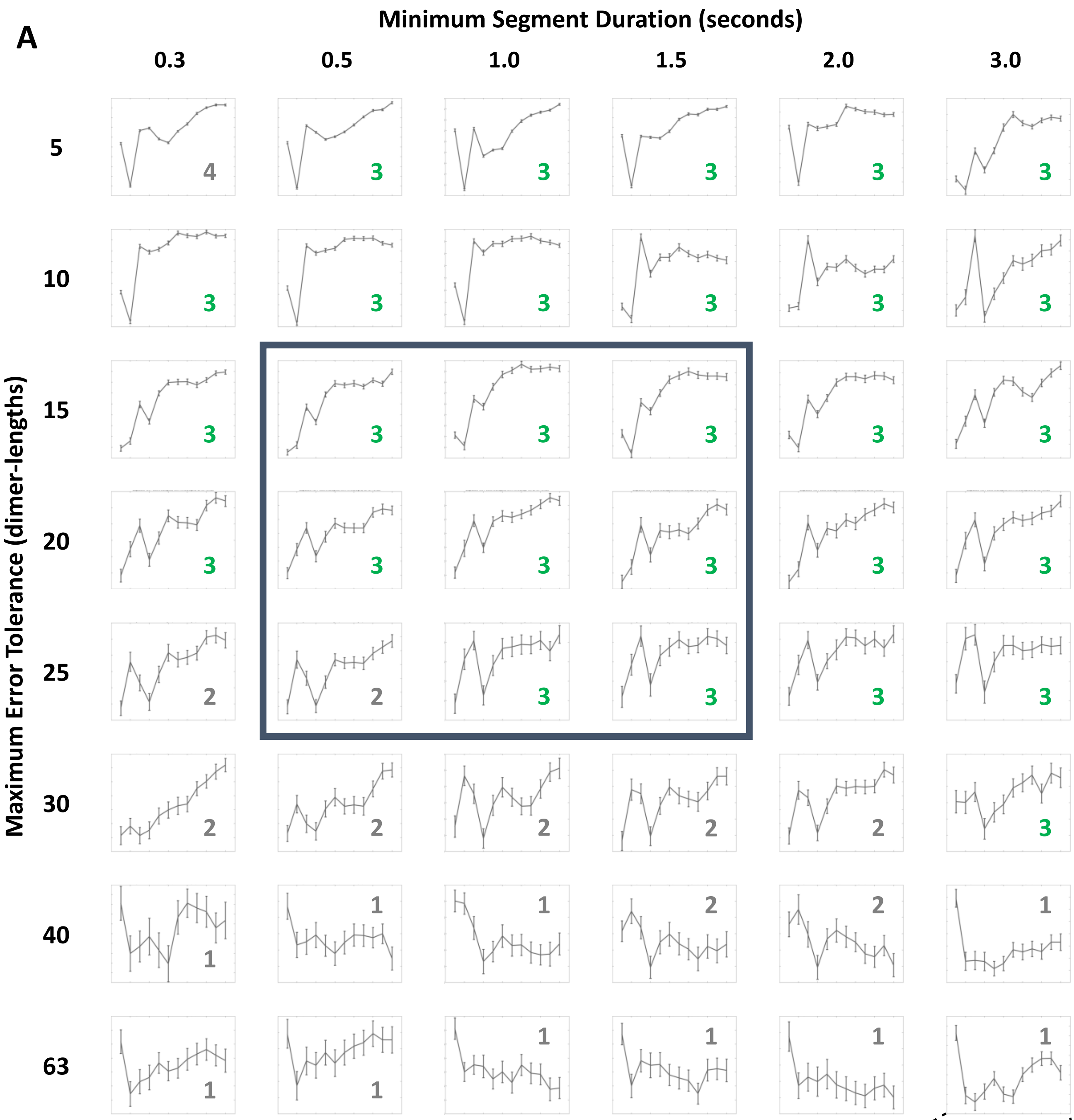


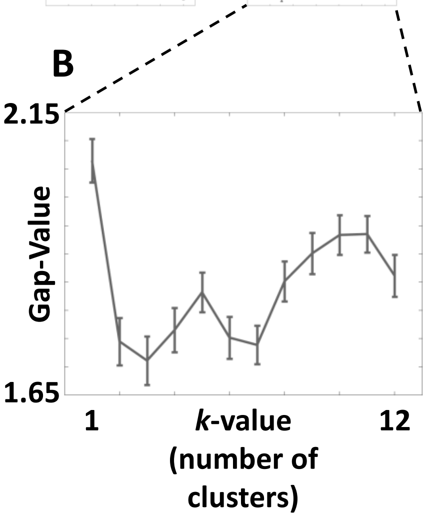


**Figure S2.1. Gap statistic plots for *in silico* data (Positive Slope Segments) support the conclusion of multiple growth behaviors when segmentation is conducted at relevant spatio-temporal scales. (A)** The gap statistic plots presented are for the positive slope segments and are the result of analyzing the **full resolution†** *in silico* dataset using the Diagnostic Mode of STADIA with the Minimum Segment Duration and Maximum Error Tolerance indicated by the column and row headings, respectively. Each gap statistic plot is labeled with the optimal *k*-value suggested (green is used to indicate agreement with the results for this dataset in the main text; gray indicates parameter combinations resulting in different *k*-values.). The **dark blue** box indicates the parameter space for which cluster profiles and labeled length-history plots can be found in **Figures S2.3** and **S2.4**, respectively. **(B)** A representative gap statistic plot (bottom right) shows the axes for each plot in (A). The x-axis (*k*-value) range is the same for all plots. The y-axis (Gap-Value) has differing ranges (not shown) for each plot, but the specific numerical values of the gap statistic are not relevant to interpreting the plots because identification of the optimal *k*-value is based on local maxima within each gap statistic plot. In other words, the pertinent information is the relationship between the values of the gap statistic at different *k*-values within each plot, not the values themselves. **†** The **full resolution** *in silico* data have temporal resolution of one output per dimer-scale biochemical event (see Methods).

**Figure S2.1 – Observations and Interpretations**

**Observations:** Within the full resolution *in silico* dataset, the *k*-value suggested by the gap statistic is relatively unchanged for the positive slope segments across the range of Minimum Segment Durations analyzed (e.g., for a Maximum Error Tolerance of 20 dimer-lengths (160 nm), the gap statistic suggests *k*=3 as the optimal number of clusters for the positive slope segment data regardless of the Minimum Segment Duration). Changing the Maximum Error Tolerance has a markedly higher impact on the *k*-value suggested by the gap statistic: there is a clear trend towards lower *k*-values at higher Maximum Error Tolerances.

**Interpretations:** Recall that the Minimum Segment Duration places a lower limit on the timescale of behaviors being analyzed, while the Maximum Error Tolerance places an upper limit on the difference between the piecewise linear approximation and the inputted length-history data. Limiting the timescale accuracy of the segmentation, even with a Minimum Segment Duration of 3.0 seconds, does not appear to change the conclusion that *k*=3 for the positive slope segment data. Further, if the Maximum Error Tolerance is less than 30 dimer-lengths, the conclusion that *k*=3 is also upheld. Using this information in combination with the cluster profiles in **Figure S2.3**, we can conclude that the existence of stutters as a distinct behavior is a robust conclusion for the positive slope segment data of the full resolution *in silico* microtubules, but analysis must be conducted using segmentation parameters that are reasonable for capturing behaviors at the scale that stutters exist.

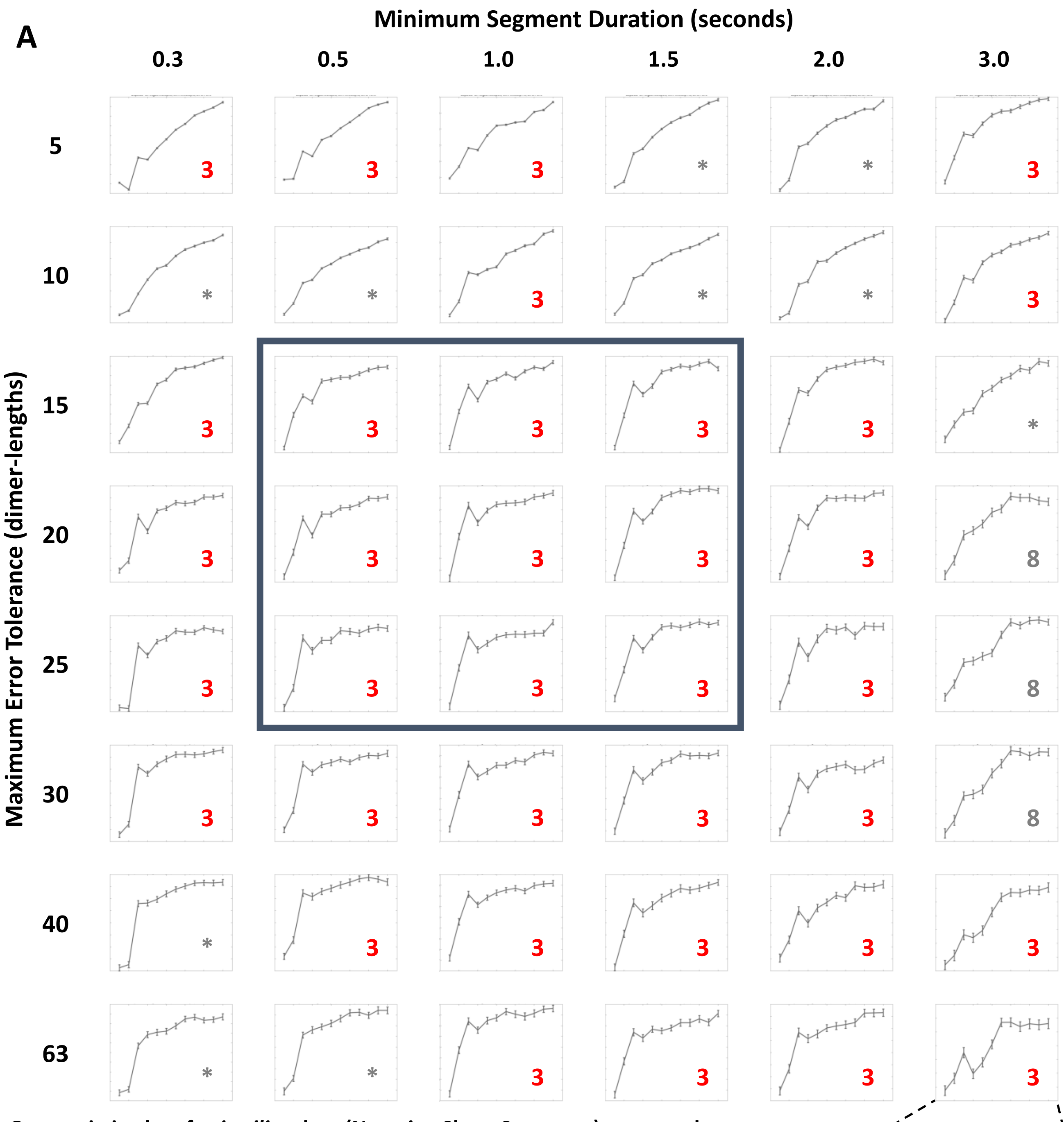


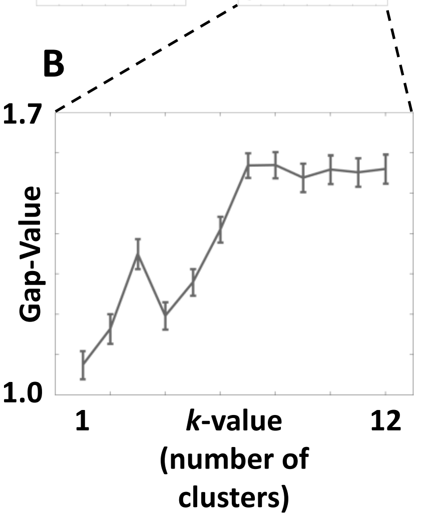


**Figure S2.2. Gap statistic plots for *in silico* data (Negative Slope Segments) support the conclusion of multiple growth behaviors when segmentation is conducted at relevant spatio-temporal scales. (A)** The gap statistic plots presented are for the negative slope segments and are the result of analyzing the **full resolution†** *in silico* dataset using the Diagnostic Mode of STADIA with the Minimum Segment Duration and Maximum Error Tolerance indicated by the column and row headings, respectively. Each gap statistic plot is labeled with the optimal *k*-value suggested (**red** is used to indicate agreement with the results for this dataset in the main text; **gray** indicates parameter combinations resulting in different *k*-values. Plots with * are plots where a clear local maximum is not evident and are 'uninterpretable gap statistics'.). The **dark blue** box indicates the parameter space for which cluster profiles and labeled length-history plots can be found in **Figure S2.3** and **S2.4**, respectively. **(B)** A representative gap statistic plot (bottom right) shows the axes for each plot in (A). The x-axis (*k*-value) range is the same for all plots. The y-axis (Gap-Value) has differing ranges (not shown) for each plot, but the specific numerical values of the gap statistic are not relevant to interpreting the plots because identification of the optimal *k*-value is based on local maxima within each gap statistic plot. In other words, the pertinent information is the relationship between the values of the gap statistic at different *k*-values within each plot, not the values themselves. **†** The **full resolution** *in silico* data have temporal resolution of one output per dimer-scale biochemical event (see Methods).

**<u>Figure S2.2 – Observations and Interpretations</u>**

**Observations:** Similar to the *in silico* positive slope segment data, within the full resolution *in silico* dataset, the *k*-value suggested by the gap statistic is relatively unchanged for the negative slope segment data across the range of Minimum Segment Durations analyzed (e.g., for a Maximum Error Tolerance of 20 dimer-lengths, the gap statistic suggests *k*=3 as the optimal number of clusters for the negative slope segment data for Minimum Segment Durations less than 3.0 seconds). Notable exceptions exist particularly at low Maximum Error Tolerances (5 or 10 dimer-lengths) which often result in gap statistic plots that are generally increasing and therefore not conclusive.

Changing the Maximum Error Tolerance has little impact on the *k*-value suggested by the gap statistic for the negative slope segments compared to the impact for the positive slope segments (e.g., for a Minimum Segment Duration of 1.0 seconds, *k*=3 is suggested by the gap statistic for all Maximum Error Tolerances tested). At long Minimum Segment Durations, the *k*-value suggested by these gap statistic plots is either not determinable or a high value (e.g., *k*=8; the gap statistic plots rely on clustering results from 500 random starts, and it is reasonable to expect some volatility especially at high *k*-values, so the optimal *k*-value suggested by these plots should be taken with caution).

**Interpretations:** Recall that the Minimum Segment Duration places a lower limit on the timescale of behaviors being analyzed, while the Maximum Error Tolerance places an upper limit on the difference between the piecewise linear approximation and the inputted length-history data. Limiting the timescale accuracy of the segmentation, even with a Minimum Segment Duration of 2.0 seconds, does not appear to change the conclusion that *k*=3 for the negative slope segment data. Further, for Maximum Error Tolerances from 15 to 40 dimer-lengths, the conclusion that *k*=3 is also supported in most cases. Using this information in combination with the cluster profiles in **Figure S2.3**, we can conclude that the existence of stutters as a distinct behavior is a robust conclusion for the negative slope segment data of the full resolution *in silico* microtubules, but analysis must be conducted using segmentation parameters that are reasonable for capturing behaviors at the scale that stutters exist.

**Additional discussion:** With regard to the gap statistic plots at long Minimum Segment Durations, consider that shortening or depolymerization behaviors exhibited by MTs often do not last 3 seconds (**Figure S1.8**), so it is conceivable that thresholds are interfering even with the appropriate detection of shortening phases when analysis is conducted at this scale. Thus, by the very nature of shortening phases of MTs, it follows that the gap statistic plots at long Minimum Segment Durations should be treated with caution.

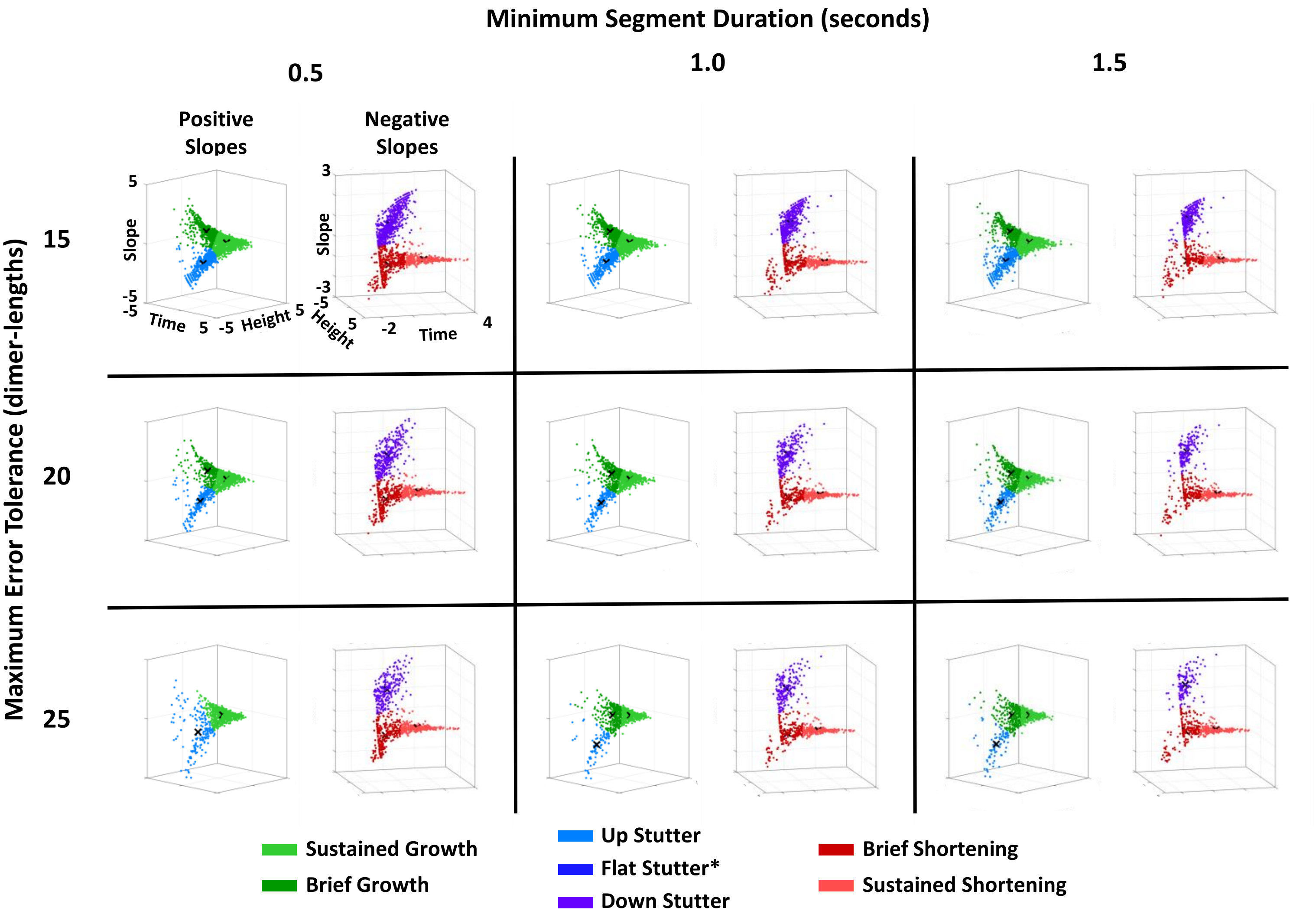


**Figure S2.3. Cluster profiles of positive and negative slope segment data for the full resolution *in silico* dataset provide support for conclusions suggested by the gap statistic plots.** The cluster plots represent clustering results from using STADIA in Automated Mode with the *k*-values indicated by the corresponding gap statistic plots in **Figures S2.1** and **S2.2**. Note that the data plotted in this figure is log-transformed and standardized (see Methods). ***Flat stutters** are not included in the cluster profiles because flat stutter segments are identified by user-defined thresholds, not by *k*-means clustering (see Methods and **Figure S1.1**).

**Figure S2.3 – Observations and Interpretations**

**Observations:** The cluster profiles for the positive and negative slope segment data maintain the same general shape over the range of Maximum Error Tolerances and Minimum Segment Durations shown here (i.e., the cluster profile still has 3 'appendages'). For both the positive and negative slope segments, a notable difference across the varying parameter values is that the overall density of data points decreases as the Maximum Error Tolerance and/or the Minimum Segment Duration is increased (i.e., highest density in the upper left of the grid, and lowest density in the bottom right of the grid).

**Interpretations:** The relative stability of the shape of the cluster profiles for the positive and negative slope segments bolsters the conclusions drawn from the gap statistic plots in **Figure S2.1** and **S2.2** (recall that the gap statistic drives the decision for the optimal *k*-value, but the cluster profiles are also used to inform the *k*-value). Additionally, with the use of the cluster profiles, we can see that while the number of clusters is generally unaffected by changing the user-input parameters, the shapes and centroid locations of the actual clusters can change (e.g., compare the positive slope segment clusters in the top left to the bottom right). Note that the appendage of the data structure with the shallow-slope segments (i.e., stutters) is present in all cases shown, even in the case where the gap plot indicated *k*=2 for the positive slope segments (Minimum Segment Duration = 0.5 second, Maximum Error Tolerance = 25 dimer-lengths).

With regards to the decrease in the density of the clusters as the input parameters increase, recall that the Minimum Segment Duration and the Maximum Error Tolerance directly affect the accuracy of the piecewise linear approximation. Higher values of Minimum Segment Duration and Maximum Error Tolerance result in a less accurate approximation and thus fewer line segments. Because segments are represented as data points in the cluster profile, fewer segments leads to cluster profiles with lower density.

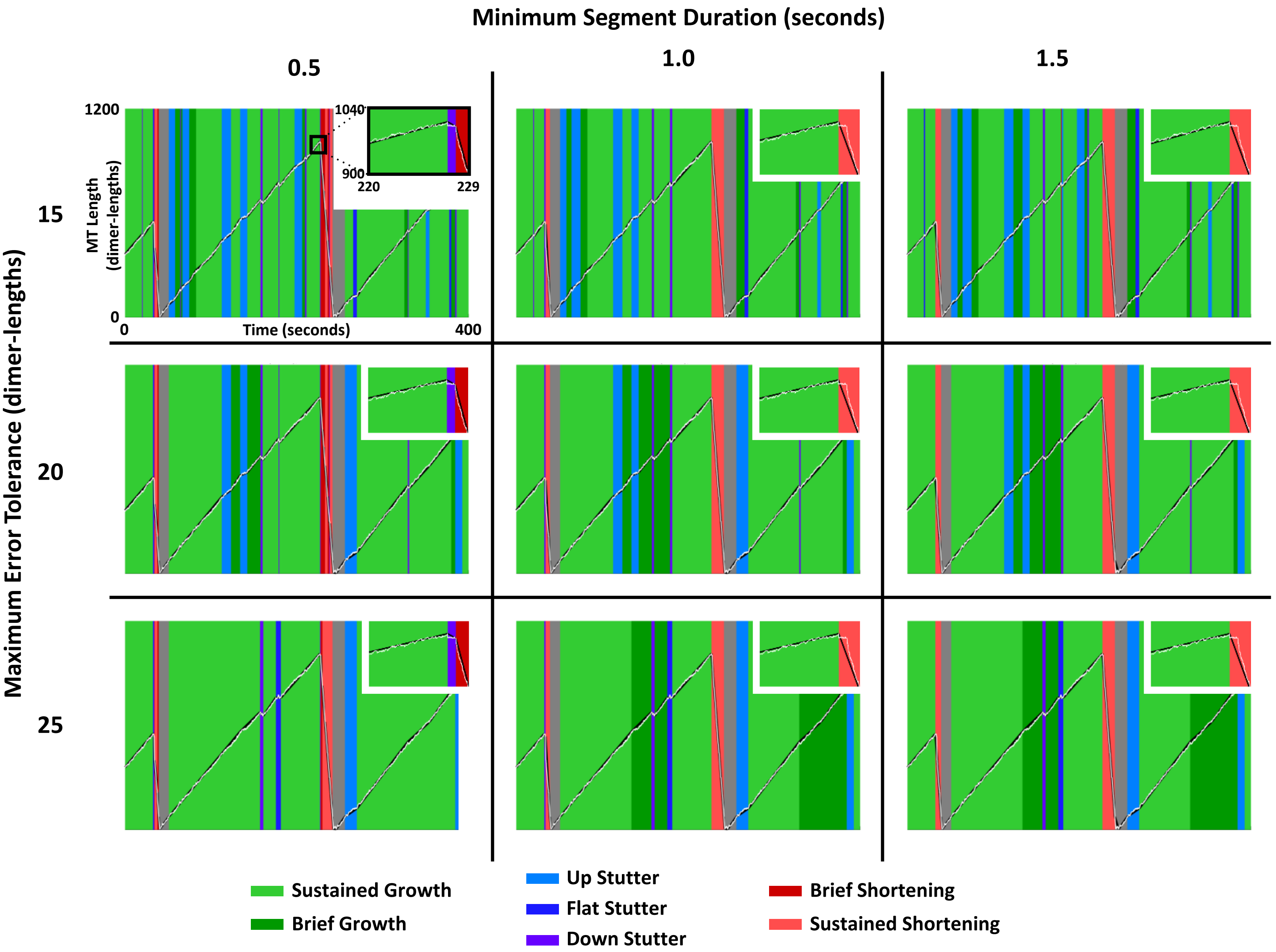


**Figure S2.4. Labeled length-history data for the full resolution *in silico* dataset demonstrate reduced detection of stutters when segmentation is performed using higher values for Minimum Segment Duration or Maximum Error Tolerance.** In the length-history plots, each segment is labeled based on the classification results from using STADIA in Automated Mode with the *k*-values indicated for each set of parameters by the corresponding gap statistic plots in **Figures S2.1** and **S2.2**. The zoomed-in portraits show the catastrophe indicated by the black box in the top right panel.

__Figure S2.4 – Observations and Interpretations__

**Observations:** The labeled length-history data illustrate that detection of stutters is dependent on the Minimum Segment Duration and the Maximum Error Tolerance. For example, the zoomed in portraits in the upper right corner of each length-history plot show a clear transitional catastrophe, outlined by the black box in the upper left plot, in the length-history data (white line) that is miscategorized as abrupt at Minimum Segment Duration values > 0.5 seconds. This transitional catastrophe is miscategorized because the piecewise linear approximation (black line segments) does not segment the data with enough accuracy to detect this stutter when using Minimum Segment Durations > 0.5 seconds. For increasing values of Maximum Error Tolerance, fewer stutters are detected throughout the plotted region of length-history data.

**Interpretations:** Labeled length-history plots provide a qualitative check on the classification and transition analyses and provide further insight into appropriate parameter choices. While the gap statistic and cluster profiles inform the *k*-value (i.e., the number of behaviors) and robustness with which those behaviors are detected (i.e., how often STADIA detects a behavior that is in the data), the labeled length-history data allow for visual inspection of the transitions between behaviors. For example, the labeled length-history plots shown here indicate that the parameters chosen for STADIA analysis throughout the paper (i.e., Minimum Segment Duration = 0.5 seconds, Maximum Error Tolerance = 20 dimer-lengths) are appropriate for accurately detecting the stutter before catastrophe shown here. The effect of the user-input parameters on the detection of transitional catastrophes more generally is examined in **Figures S2.9-S2.11**.

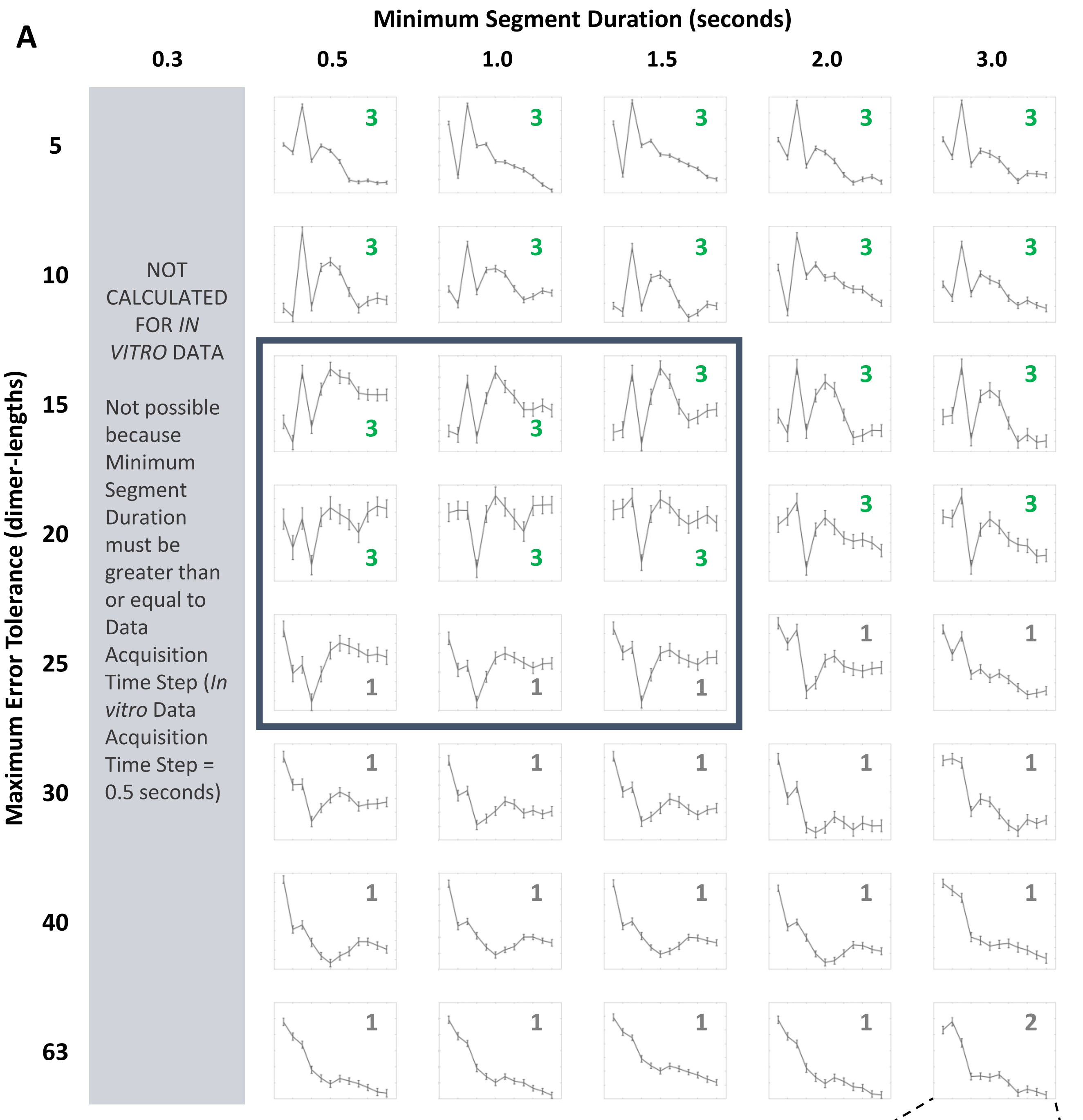


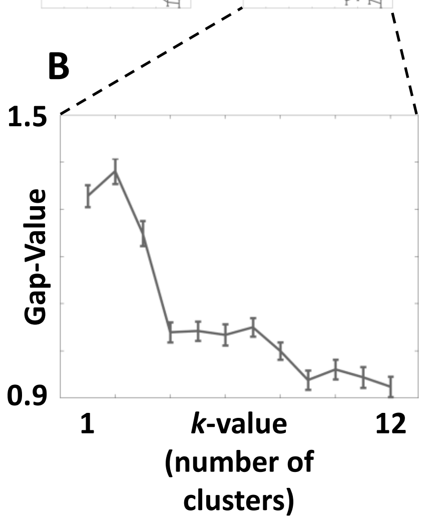


**Figure S2.5. Gap statistic plots for *in vitro* control data (Positive Slope Segments) support conclusion of multiple growth behaviors when segmentation is conducted at relevant spatiotemporal scales. (A)** The gap statistic plots presented are for the positive slope segments and are the result of analyzing the *in vitro* control dataset using the Diagnostic Mode of STADIA with the Minimum Segment Duration and Maximum Error Tolerance indicated by the column and row headings, respectively. Each gap statistic plot is labeled with the optimal *k*-value suggested (**green** is used to indicate agreement with the results for this dataset in the main text; **gray** indicates parameter combinations resulting in different *k*-values.). The **dark blue** box indicates the parameter space for which cluster profiles and labeled length-history plots can be found in **Figure S2.7** and **Figure S2.8**, respectively. **(B)** A representative gap statistic plot (bottom right) shows the axes for each plot in (A). The x-axis (*k*-value) range is the same for all plots. The y-axis (Gap-Value) has differing ranges (not shown) for each plot, but the specific numerical values of the gap statistic are not relevant to interpreting the plots because identification of the optimal *k*-value is based on local maxima within each gap statistic plot. In other words, the pertinent information is the relationship between the values of the gap statistic at different *k*-values within each plot, not the values themselves.

**<u>Figure S2.5 – Observations and Interpretations</u>**

**Observations:** Relatively consistent with what was seen for the full resolution *in silico* dataset (**Figure S2.1**), within the *in vitro* control dataset, the *k*-value suggested by the gap statistic is relatively unchanged for the positive slope segment data across the range of Minimum Segment Durations analyzed (e.g., for a Maximum Error Tolerance of 20, the gap statistic suggests *k*=3 as the optimal number of clusters for the positive slope segment data regardless of the Minimum Segment Duration). Changing the Maximum Error Tolerance has a markedly higher impact on the *k*-value suggested by the gap statistic: there is a clear trend towards lower *k*-values at Maximum Error Tolerances > 25 dimer-lengths.

**Interpretations:** Recall that the Minimum Segment Duration places a lower limit on the timescale of behaviors being analyzed, while the Maximum Error Tolerance places an upper limit on the difference between the piecewise linear approximation and the inputted length-history data. Limiting the timescale accuracy of the segmentation, even with a Minimum Segment Duration of 3.0 seconds, does not appear to change the conclusion that *k*=3 for the positive slope segment data. Further, if the Maximum Error Tolerance is less than 25 dimer-lengths, the conclusion that *k*=3 is also upheld. Using this information in combination with the cluster profiles in **Figure S2.7**, we can conclude that the existence of stutters as a distinct behavior is a robust conclusion for the positive slope segment data of the *in vitro* control microtubules, but analysis must be conducted using segmentation parameters that are reasonable for capturing behaviors at the scale that stutters exist.

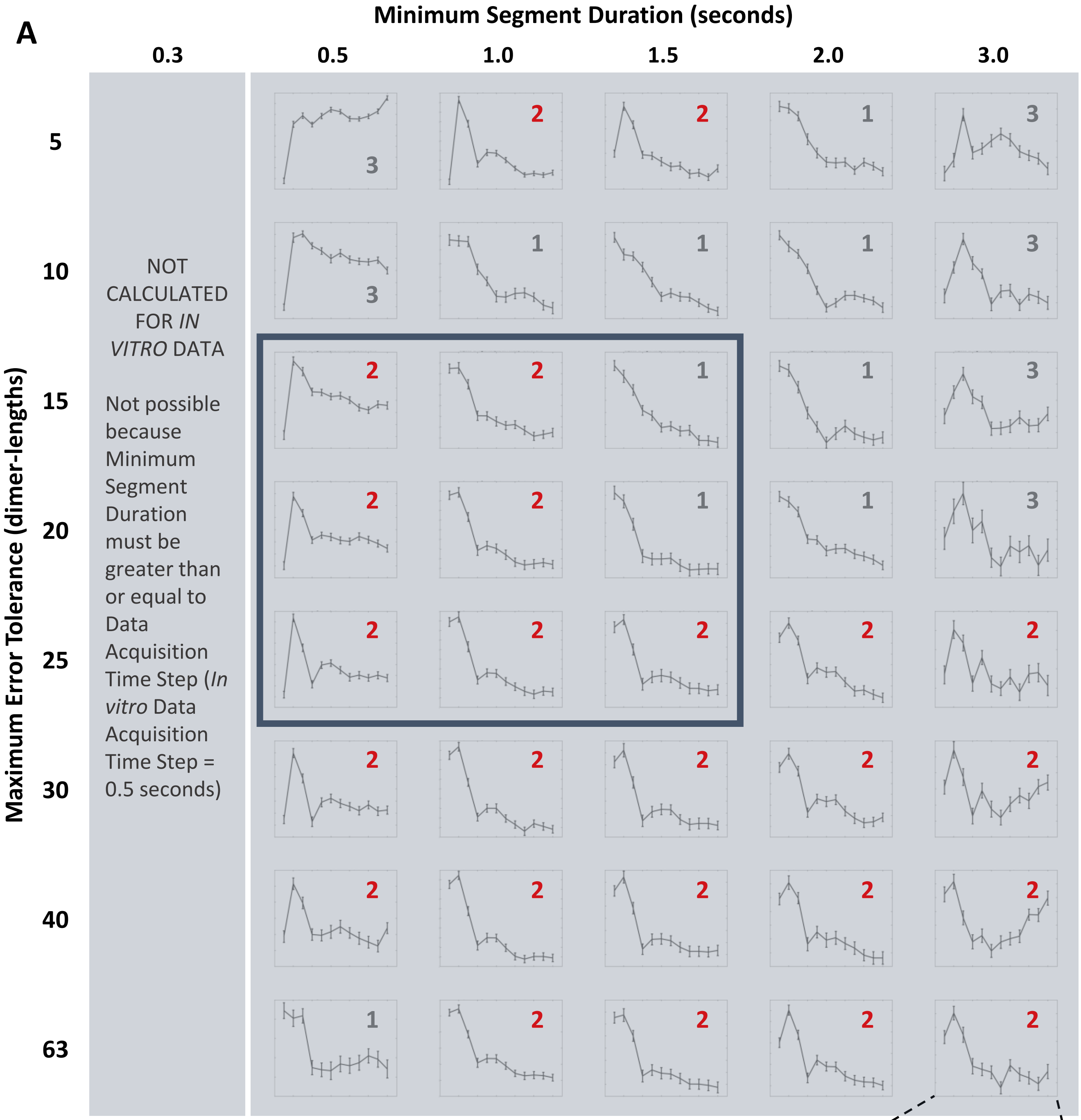


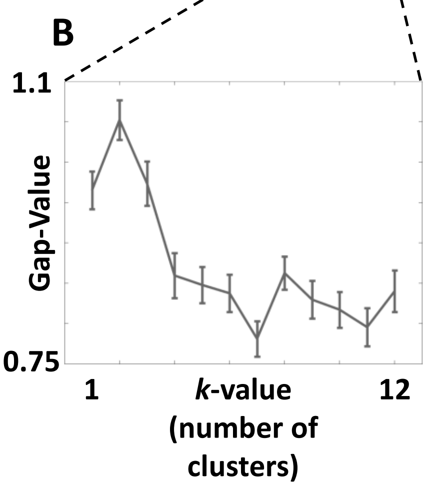


**Figure S2.6. Gap statistic plots for *in vitro* control data (Negative Slope Segments) yield inconclusive results regarding the number of behaviors detected when segmentation is conducted at relevant spatio-temporal scales. (A)** The gap statistic plots presented are for the negative slope segments and are the result of analyzing the full *in vitro* control dataset using the Diagnostic Mode of STADIA with the Minimum Segment Duration and Maximum Error Tolerance indicated by the column and row headings, respectively. Each gap statistic plot is labeled with the optimal *k*-value suggested (**red** is used to indicate agreement with the results for this dataset in the main text; **gray** indicates parameter combinations resulting in different *k*-values.). The **dark blue** box indicates the parameter space for which cluster profiles and labeled length-history plots can be found in **Figure S2.7** and **Figure S2.8**, respectively. The gray shading over the gap statistic plots is used to indicate the inconclusiveness of these results (recall that depolymerizations were not captured in their entirety for *in vitro* data). **(B)** A representative gap statistic plot (bottom right) shows the axes for each plot in (A). The x-axis (*k*-value) range is the same for all plots. The y-axis (Gap-Value) has differing ranges (not shown) for each plot, but the specific numerical values of the gap statistic are not relevant to interpreting the plots because identification of the optimal *k*-value is based on local maxima within each gap statistic plot. In other words, the pertinent information is the relationship between the values of the gap statistic at different *k*-values within each plot, not the values themselves.

**Figure S2.6 – Observations and Interpretations**

**Observations:** At Maximum Error Tolerances greater than 20 dimer-lengths, the gap statistic consistently suggests $k$=2 (in agreement with the results in the main text of the paper) as the optimal number of clusters for the negative slope segments in the *in vitro* control data. At lower Maximum Error Tolerances, however, there seems to be much more volatility. Recall that depolymerizations for the *in vitro* datasets (both with and without CLASP2γ) were not captured in their entirety, and so classification analysis of the negative slope segment data in this parameter sweep is not conclusive.

**Interpretations:** As indicated in the observations, interpreting these gap statistic plots is difficult because depolymerizations were only partially captured in the kymographs (and subsequently in the piecewise linear approximation). Even within this limited dataset, the gap statistic plots do indicate a relatively steady conclusion that multiple behaviors exist. However, explanations for the volatility in $k$-values suggested by the gap statistic are difficult without complete negative slope segment data.

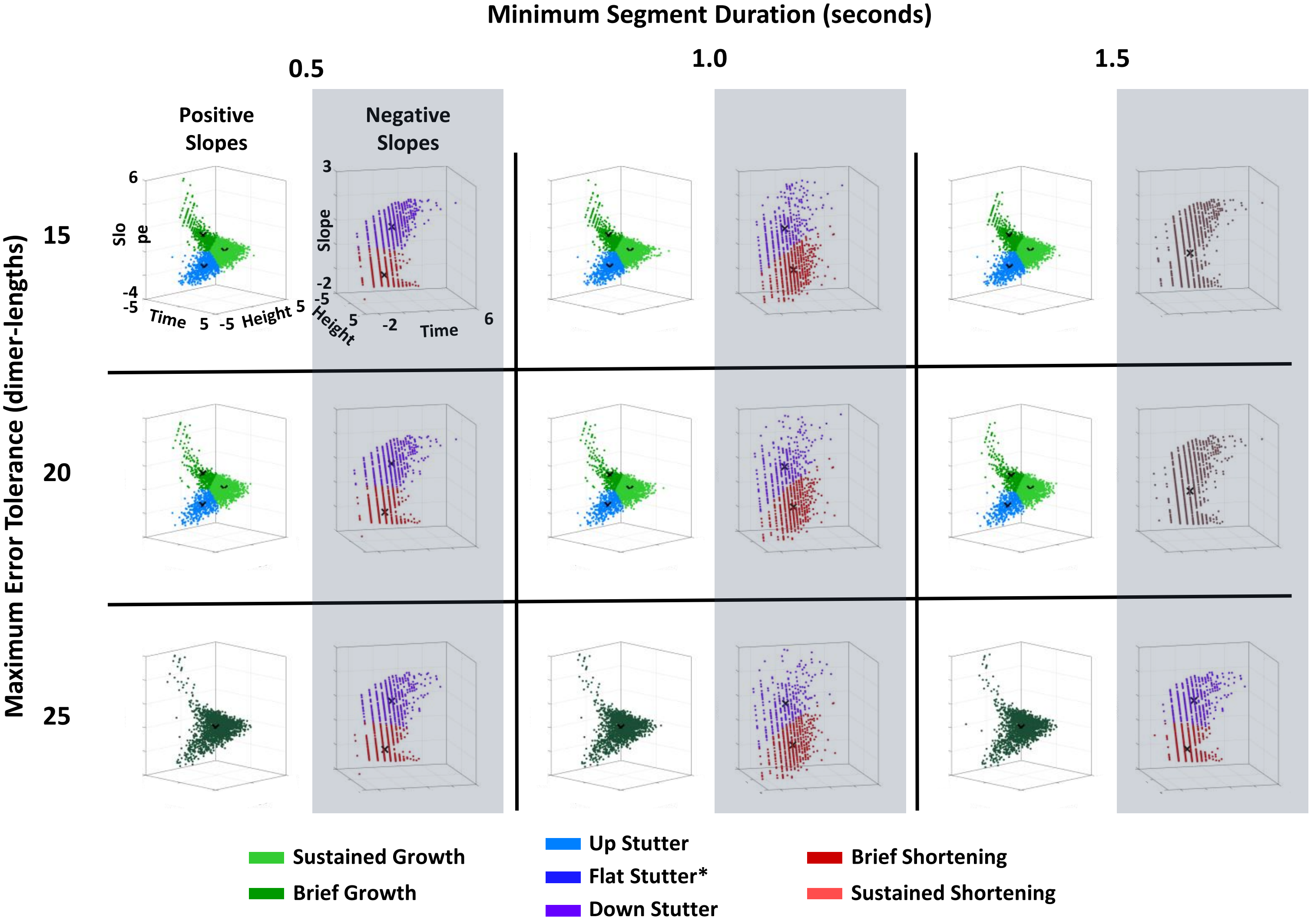


**Figure S2.7. Cluster profiles of positive and negative slope segment data for the *in vitro* control dataset provide support for conclusions suggested by the gap statistic plots.** The cluster plots represent clustering results from using STADIA in Automated Mode with the *k*-values indicated by the corresponding gap statistic plots in **Figure S2.5** and **S2.6**. The gray shading over the cluster profiles for the negative slope line segment data is used to indicate the inconclusiveness of these results (recall that depolymerizations were not captured in their entirety for the *in vitro* data). Note that the data plotted in this figure is log-transformed and standardized (see Methods). ***Flat stutters** are not included in the cluster profiles because flat stutter segments are identified by user-defined thresholds, not by *k*-means clustering (see Methods and **Figure S1.1**).

**Figure S2.7 – Observations and Interpretations**

**Observations:** Similar to what was observed for the *in silico* dataset, cluster profiles for positive slope segment data maintain the same general shape over the range of Maximum Error Tolerances or Minimum Segment Durations shown in the figure (i.e., there are still 3 'appendages' to the cluster profile for the positive slope segments).

Of note is that for a Maximum Error Tolerance of 25 dimer-lengths (where $k$=1 was suggested by the gap statistic), the general shape of the data is still consistent with the positive slope segment cluster profiles for smaller Maximum Error Tolerances. Overall, there appears to be less change in density of the data points (segments ~~in~~ from the piecewise linear approximation) for the *in vitro* data over this subset of the parameter space compared to the differences noticed in the *in silico* dataset (**Figure S2.3**). However, when loss of density does occur, much of the loss appears to be among the more rapid short-duration segments of the brief growth cluster, which is similar to the *in silico* results.

For the negative slope segment cluster profiles (again, incomplete because the experimentally obtained dataset contained only the beginning of depolymerization phases), the overall shape of the cluster profiles is fairly consistent across the parameter space. Even at Minimum Segment Duration = 1.5 seconds and Maximum Error Tolerances of 15 or 20 dimer-lengths (where the gap statistic suggested $k$=1), the overall shape is similar to the other cluster profiles.

**Interpretations:** The relative stability of the shape of the cluster profiles for positive slope segments bolsters the conclusions drawn from the gap statistic plots in **Figure S2.5** and **S2.6** (recall that the gap statistic drives the decision for the optimal $k$-value, but cluster profiles are also used to inform the $k$-value). Additionally, even when the gap statistic suggests a different $k$-value (which occurs for the positive slope segment data when analyzed using Maximum Error Tolerance of 25 dimer-lengths), the lack of major change in the cluster profile supports the conclusion that multiple behaviors exist. In particular, the portion of the data structure with the shallow-slope segments (i.e., stutters) is present in all cases shown.

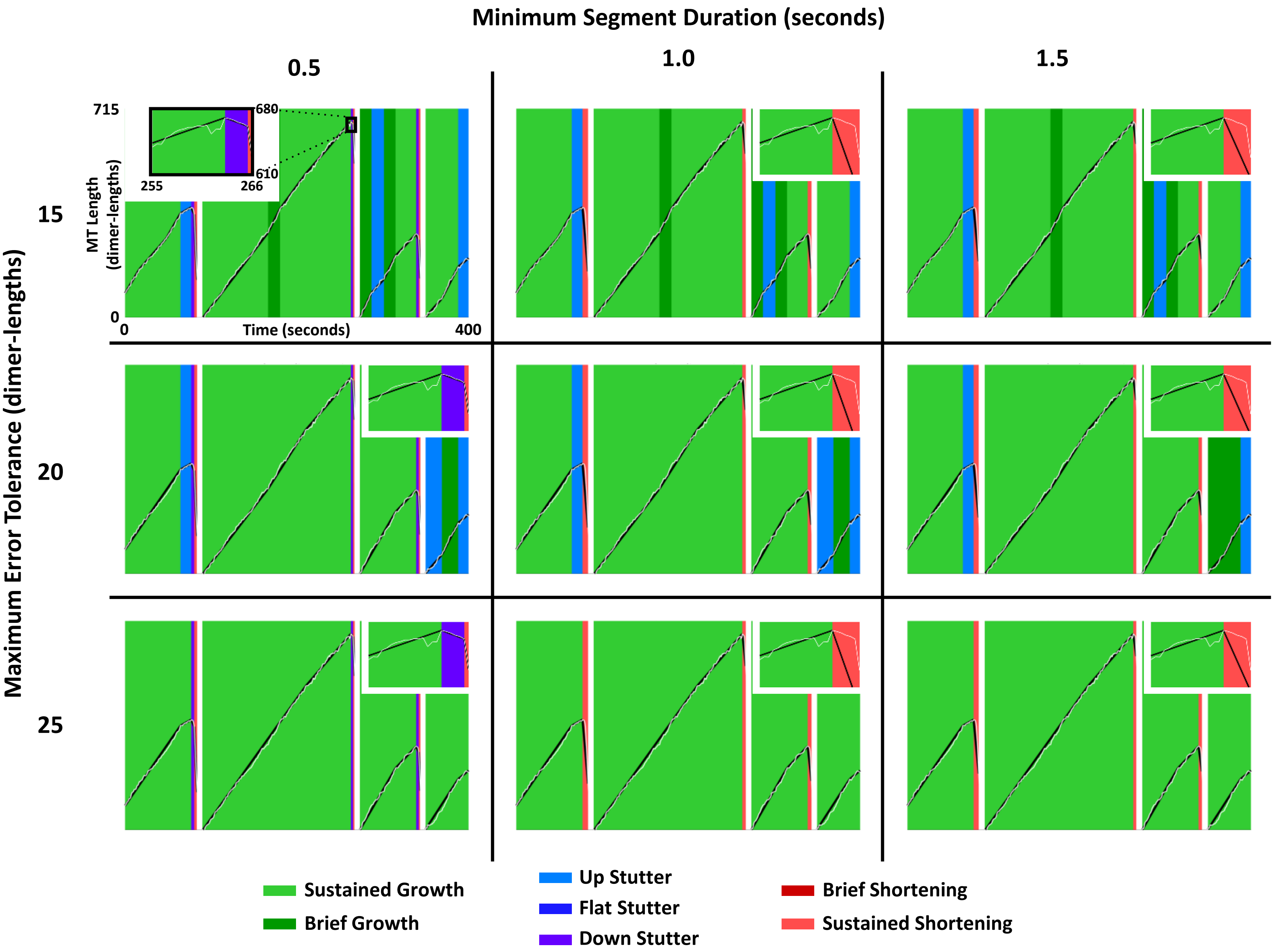


**Figure S2.8. Labeled length-history data for the *in vitro* control dataset demonstrate reduced detection of stutters when segmentation is performed using higher values for Minimum Segment Duration or Maximum Error Tolerance.** In the length-history plots, each segment is labeled based on the classification results from using STADIA in Automated Mode with the *k*-values indicated for each set of parameters by the corresponding gap statistic plots in **Figures S2.5** and **S2.6**. The zoomed-in portraits show the catastrophe indicated by the black box in the top right panel.

__Figure S2.8 – Observations and Interpretations__

**Observations:** Comparable to the labeled length-history plots from the analysis of the *in silico* dataset, the labeled length-history plots for the *in vitro* data illustrate that detection of stutters is dependent on the Minimum Segment Duration and the Maximum Error Tolerance. For example, the zoomed-in portraits in the upper right corner of each length-history plot show a clear transitional catastrophe, outlined by the black box in the upper left plot, in the length-history data (white line) that is miscategorized as abrupt at Minimum Segment Durations > 0.5 seconds. This transitional catastrophe is miscategorized because the piecewise linear approximation (black line segments) does not segment the data with enough accuracy to detect this stutter when using Minimum Segment Durations > 0.5 seconds. For increasing values of Maximum Error Tolerance, fewer stutters are detected throughout the plotted region of length-history data.

**Interpretations:** Labeled length-history plots provide a qualitative check on the classification and transition analyses and provide further insight into appropriate parameter choices. While the gap statistic and cluster profiles inform the *k*-value (i.e., the number of behaviors) and robustness with which those behaviors are detected (i.e., how often STADIA detects a behavior that is in the data), the labeled length-history data allow for visual inspection of the transitions between behaviors. For example, the labeled length-history plots shown here indicate that the parameters chosen for STADIA analysis throughout the paper (i.e., Minimum Segment Duration = 0.5 seconds, Maximum Error Tolerance = 20 dimer-lengths) are appropriate for accurately detecting the stutter before catastrophe shown here. The effect of the user-input parameters on the detection of transitional catastrophes more generally is examined in **Figures S2.9-S2.11**.

**A**

| Frequency of ABRUPT CATASTROPHE | | | | | | |
|---|---|---|---|---|---|---|
| Max Error Tolerance (dimer-lengths) | Minimum Segment Duration (s) | | | | | |
| | 0.3 | 0.5 | 1.0 | 1.5 | 2.0 | 3.0 |
| 5 | 0.0883 | 0.1107 | 0.1002 | 0.0642 | 0.0439 | 0.0186 |
| 10 | 0.0050 | 0.0067 | 0.0105 | 0.0149 | 0.0158 | 0.0151 |
| 15 | 0.0046 | 0.0029 | 0.0045 | 0.0061 | 0.0071 | 0.0075 |
| 20 | 0.0024 | 0.0025 | 0.0041 | 0.0059 | 0.0068 | 0.0070 |
| 25 | 0.0023 | 0.0031 | 0.0042 | 0.0057 | 0.0067 | 0.0068 |
| 30 | 0.0025 | 0.0027 | 0.0042 | 0.0059 | 0.0068 | 0.0068 |
| 40 | 0.0029 | 0.0031 | 0.0043 | 0.0059 | 0.0068 | 0.0066 |
| 63 | 0.0047 | 0.0048 | 0.0046 | 0.0060 | 0.0070 | 0.0057 |

**B**

| Frequency of TRANSITIONAL CATASTROPHE | | | | | | |
|---|---|---|---|---|---|---|
| Max Error Tolerance (dimer-lengths) | Minimum Segment Duration (s) | | | | | |
| | 0.3 | 0.5 | 1.0 | 1.5 | 2.0 | 3.0 |
| 5 | 0.0607 | 0.0648 | 0.0402 | 0.0217 | 0.0127 | 0.0052 |
| 10 | 0.0131 | 0.0122 | 0.0094 | 0.0076 | 0.0068 | 0.0045 |
| 15 | 0.0101 | 0.0095 | 0.0071 | 0.0053 | 0.0042 | 0.0030 |
| 20 | 0.0090 | 0.0085 | 0.0060 | 0.0042 | 0.0030 | 0.0021 |
| 25 | 0.0083 | 0.0074 | 0.0055 | 0.0038 | 0.0026 | 0.0019 |
| 30 | 0.0075 | 0.0072 | 0.0052 | 0.0035 | 0.0024 | 0.0017 |
| 40 | 0.0067 | 0.0064 | 0.0049 | 0.0033 | 0.0023 | 0.0018 |
| 63 | 0.0045 | 0.0043 | 0.0040 | 0.0029 | 0.0019 | 0.0009 |

**C**

| Frequency of INTERRUPTED GROWTH | | | | | | |
|---|---|---|---|---|---|---|
| Max Error Tolerance (dimer-lengths) | Minimum Segment Duration (s) | | | | | |
| | 0.3 | 0.5 | 1.0 | 1.5 | 2.0 | 3.0 |
| 5 | 0.5378 | 0.3763 | 0.1998 | 0.1404 | 0.1000 | 0.0574 |
| 10 | 0.1321 | 0.1155 | 0.0758 | 0.0537 | 0.0431 | 0.0338 |
| 15 | 0.0332 | 0.0343 | 0.0290 | 0.0228 | 0.0198 | 0.0152 |
| 20 | 0.0104 | 0.0107 | 0.0092 | 0.0084 | 0.0079 | 0.0061 |
| 25 | 0.0033 | 0.0013 | 0.0035 | 0.0033 | 0.0032 | 0.0028 |
| 30 | 0.0015 | 0.0015 | 0.0015 | 0.0014 | 0.0014 | 0.0016 |
| 40 | 0.0003 | 0.0003 | 0.0004 | 0.0004 | 0.0004 | 0.0007 |
| 63 | 0.0000 | 0.0000 | 0.0000 | 0.0000 | 0.0001 | 0.0001 |

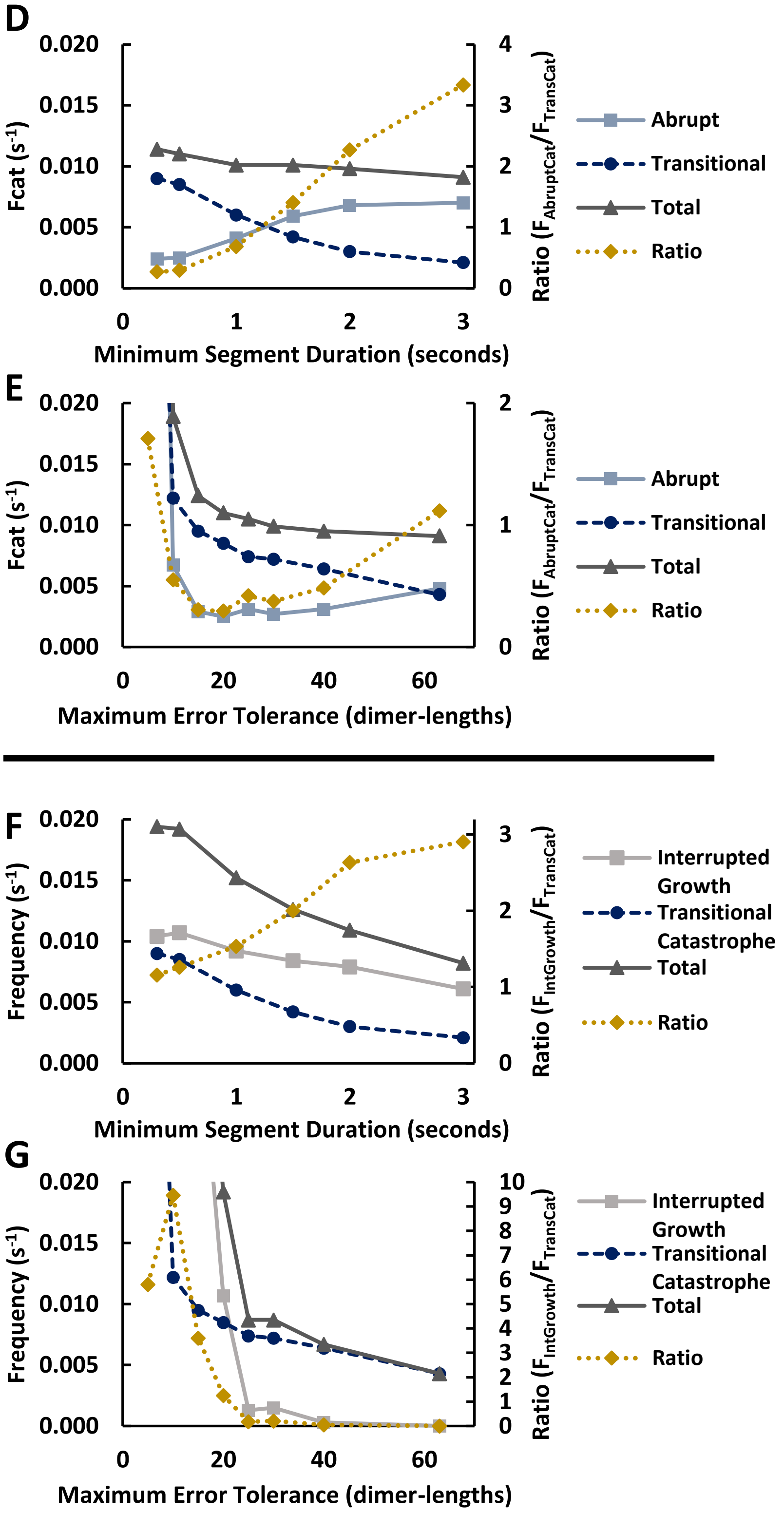


**Figure S2.9. Transition analysis across Minimum Segment Durations and Maximum Error Tolerances provide robust support for prevalence of transitional catastrophes in the full resolution dimer-scale *in silico* dataset.** These data are the results from analyzing the *in silico* dataset with STADIA using *k*=3 for both positive and negative slope segments for all user-input parameter combinations. **(A-C)** Transition frequencies across the full parameter space considered in **Supplemental Section 2**. The frequency of abrupt catastrophe ($F_{AbruptCat}$, A), frequency of transitional catastrophe ($F_{TransCat}$, B), and frequency of interrupted growth ($F_{IntGrowth}$, C) are reported in table form. The highlighted column (Minimum Segment Duration = 0.5 seconds) and row (Maximum Error Tolerance = 20 dimer-lengths) of each table correspond to data plotted in (D-G). **(D,E)** $F_{AbruptCat}$ and $F_{TransCat}$ for varying Minimum Segment Durations while holding the Maximum Error Tolerance constant at 20 dimer-lengths (D) and for varying Maximum Error Tolerances while holding the Minimum Segment Duration constant at 0.5 seconds (E). The raw frequencies are plotted corresponding to the left-hand y-axes; the ratio of $F_{AbruptCat}$ to $F_{TransCat}$ is plotted on the right-hand y-axes. (Note that parameter combinations where $F_{AbruptCat}/F_{TransCat} < 1$ indicates $F_{TransCat} > F_{AbruptCat}$, which is consistent with the conclusions in the main text for the *in silico* data). **(F,G)** $F_{TransCat}$ and $F_{IntGrowth}$ for varying Minimum Segment Durations while holding the Maximum Error Tolerance constant at 20 dimer-lengths (F) and for varying Maximum Error Tolerances while holding the Minimum Segment Duration constant at 0.5 seconds (G). The raw frequencies are plotted corresponding to the left-hand y-axes; the ratio of $F_{IntGrowth}$ to $F_{TransCat}$ is plotted on the right-hand y-axes. For small Maximum Error Tolerances, please see the tables for the frequencies that are too high to be visible in (E,G).

**Figure S2.9 A,B,D,E – Observations and Interpretations**

**Observations:** For the *in silico* data, the total $F_{cat}$ (which equals $F_{AbruptCat} + F_{TransCat}$ ) is relatively steady over the ranges of Minimum Segment Durations (D) and Maximum Error Tolerances (E) tested (a notable exception can be seen clearly for Maximum Error Tolerance = 5 dimer-lengths; see Interpretations). More importantly, the ratio between the individual frequencies ($F_{AbruptCat}/F_{TransCat}$) is consistently less than 1 (i.e., $F_{TransCat} > F_{AbruptCat}$) for Maximum Error Tolerances between 10 and 40 dimer-lengths (Minimum Segment Duration = 0.5 seconds) and for Minimum Segment Durations < 1.5 seconds (Maximum Error Tolerance = 20 dimer-lengths).

**Interpretations:** The observations above and interpretations below support the conclusion that catastrophes are more often transitional than abrupt for the full resolution *in silico* length-history data when segmentation is performed with the Minimum Segment Duration less than or equal to 1.0 seconds and the Maximum Error Tolerance between 10 and 40 dimer-lengths (i.e., between 80 and 320 nm).

For increasing Minimum Segment Durations beyond 0.5 seconds (Maximum Error Tolerance = 20 dimer-lengths, D), the decrease in $F_{TransCat}$ and increase $F_{AbruptCat}$ is indicative of a decrease in stutter detection. This effect of Minimum Segment Duration on STADIA's ability to detect stutters is also evidenced by the decreasing density of the cluster plots in **Figure S2.3** and by the loss of detection of a transitional catastrophe in the labeled length-history plots in **Figure S2.4**.

The results of varying the Maximum Error Tolerance (Minimum Segment Duration = 0.5 second, E) show that the ratio of $F_{AbruptCat}/F_{TransCat} > 1$ in two cases: Maximum Error Tolerance = 5 dimer-lengths and Maximum Error Tolerance = 63 dimer-lengths ≈ 0.5 microns. For the smaller error tolerance, the piecewise linear approximation may capture tip extensions, where collapsing tip extensions may present as abrupt catastrophes. In contrast, the larger error tolerance results in a coarse approximation, with fewer stutters detected (similar to what was seen for large Minimum Segment Durations).

**Figure S2.9 B,C,F,G – Observations and Interpretations**

**Observations:** Compared to the total $F_{cat}$ (= $F_{AbruptCat} + F_{TransCat}$ ), the frequency of growth-to-stutter transitions ($F_{IntGrowth} + F_{TransCat}$) appears to change more significantly with changes to either the Minimum Segment Duration (F) or Maximum Error Tolerance (G). For Maximum Error Tolerance fixed at 20 dimer-lengths and varying Minimum Segment Duration (F), the ratio of $F_{IntGrowth}$ to $F_{TransCat}$ is greater than 1 for all values tested (i.e., $F_{IntGrowth} > F_{TransCat}$), while both $F_{IntGrowth}$ and $F_{TransCat}$ decrease with increasing Minimum Segment Duration. For Minimum Segment Duration fixed at 0.5 seconds (G), the ratio of $F_{IntGrowth}$ to $F_{TransCat}$ is greater than 1 for Maximum Error Tolerances < 25 dimer-lengths. While $F_{IntGrowth}$ and $F_{TransCat}$ each decrease with increasing Maximum Error Tolerances, the decreases in both the total frequency ($F_{IntGrowth} + F_{TransCat}$) and the ratio can be attributed primarily to the large decrease in the frequency of interrupted growth.

**Interpretations:** The decrease in each of $F_{IntGrowth}$ and $F_{TransCat}$ with increasing Minimum Segment Durations or increasing Maximum Error Tolerances is not surprising. Both $F_{IntGrowth}$ and $F_{TransCat}$ involve the detection of stutters, which becomes more difficult at higher values of Minimum Segment Duration and/or Maximum Error Tolerance.

The large decrease in $F_{IntGrowth}$ with increasing Maximum Error Tolerance suggests that detection of stutters in growing MTs is much more sensitive to the Maximum Error Tolerance than is detection of stutters preceding catastrophes. In contrast, $F_{TransCat}$ is slightly more sensitive than $F_{IntGrowth}$ to changes in the Minimum Segment Duration.

**Additional Discussion:** For varying Minimum Segment Durations, the consistent pattern of $F_{IntGrowth} > F_{TransCat}$ is interesting as it suggests general stability in the relationship between $F_{IntGrowth}$ and $F_{TransCat}$.

**A**

| Frequency of ABRUPT CATASTROPHE | | | | | | |
|---|---|---|---|---|---|---|
| Max Error Tolerance (dimer-lengths) | Minimum Segment Duration (s) | | | | | |
| | 0.3 | 0.5 | 1.0 | 1.5 | 2.0 | 3.0 |
| 5 | Not Calculated for *In Vitro* data | 0.0012 | 0.0072 | 0.0062 | 0.0047 | 0.0037 |
| 10 | | 0.0013 | 0.0069 | 0.0059 | 0.0045 | 0.0038 |
| 15 | | 0.0015 | 0.0065 | 0.0105 | 0.0047 | 0.0040 |
| 20 | | 0.0018 | 0.0062 | 0.0098 | 0.0050 | 0.0041 |
| 25 | | 0.0030 | 0.0072 | 0.0035 | 0.0051 | 0.0046 |
| 30 | | 0.0028 | 0.0063 | 0.0059 | 0.0055 | 0.0050 |
| 40 | | 0.0032 | 0.0067 | 0.0062 | 0.0057 | 0.0053 |
| 63 | | 0.0061 | 0.0072 | 0.0068 | 0.0064 | 0.0063 |

**B**

| Frequency of TRANSITIONAL CATASTROPHE | | | | | | |
|---|---|---|---|---|---|---|
| Max Error Tolerance (dimer-lengths) | Minimum Segment Duration (s) | | | | | |
| | 0.3 | 0.5 | 1.0 | 1.5 | 2.0 | 3.0 |
| 5 | Not Calculated for *In Vitro* data | 0.0166 | 0.0069 | 0.0061 | 0.0058 | 0.0059 |
| 10 | | 0.0137 | 0.0049 | 0.0043 | 0.0045 | 0.0052 |
| 15 | | 0.0121 | 0.0032 | 0.0051 | 0.0033 | 0.0043 |
| 20 | | 0.0108 | 0.0025 | 0.0042 | 0.0026 | 0.0037 |
| 25 | | 0.0077 | 0.0002 | 0.0072 | 0.0019 | 0.0030 |
| 30 | | 0.0085 | 0.0020 | 0.0017 | 0.0018 | 0.0026 |
| 40 | | 0.0067 | 0.0015 | 0.0014 | 0.0013 | 0.0019 |
| 63 | | 0.0031 | 0.0006 | 0.0005 | 0.0005 | 0.0007 |

**C**

| Frequency of INTERRUPTED GROWTH | | | | | | |
|---|---|---|---|---|---|---|
| Max Error Tolerance (dimer-lengths) | Minimum Segment Duration (s) | | | | | |
| | 0.3 | 0.5 | 1.0 | 1.5 | 2.0 | 3.0 |
| 5 | Not Calculated for *In Vitro* data | 0.0846 | 0.0822 | 0.0739 | 0.0495 | 0.0325 |
| 10 | | 0.0205 | 0.0202 | 0.0197 | 0.0178 | 0.0156 |
| 15 | | 0.0077 | 0.0078 | 0.0058 | 0.0072 | 0.0064 |
| 20 | | 0.0036 | 0.0038 | 0.0029 | 0.0035 | 0.0034 |
| 25 | | 0.0006 | 0.0007 | 0.0006 | 0.0020 | 0.0019 |
| 30 | | 0.0010 | 0.0010 | 0.0010 | 0.0009 | 0.0009 |
| 40 | | 0.0003 | 0.0003 | 0.0002 | 0.0003 | 0.0002 |
| 63 | | 0.0001 | 0.0001 | 0.0001 | 0.0001 | 0.0000 |

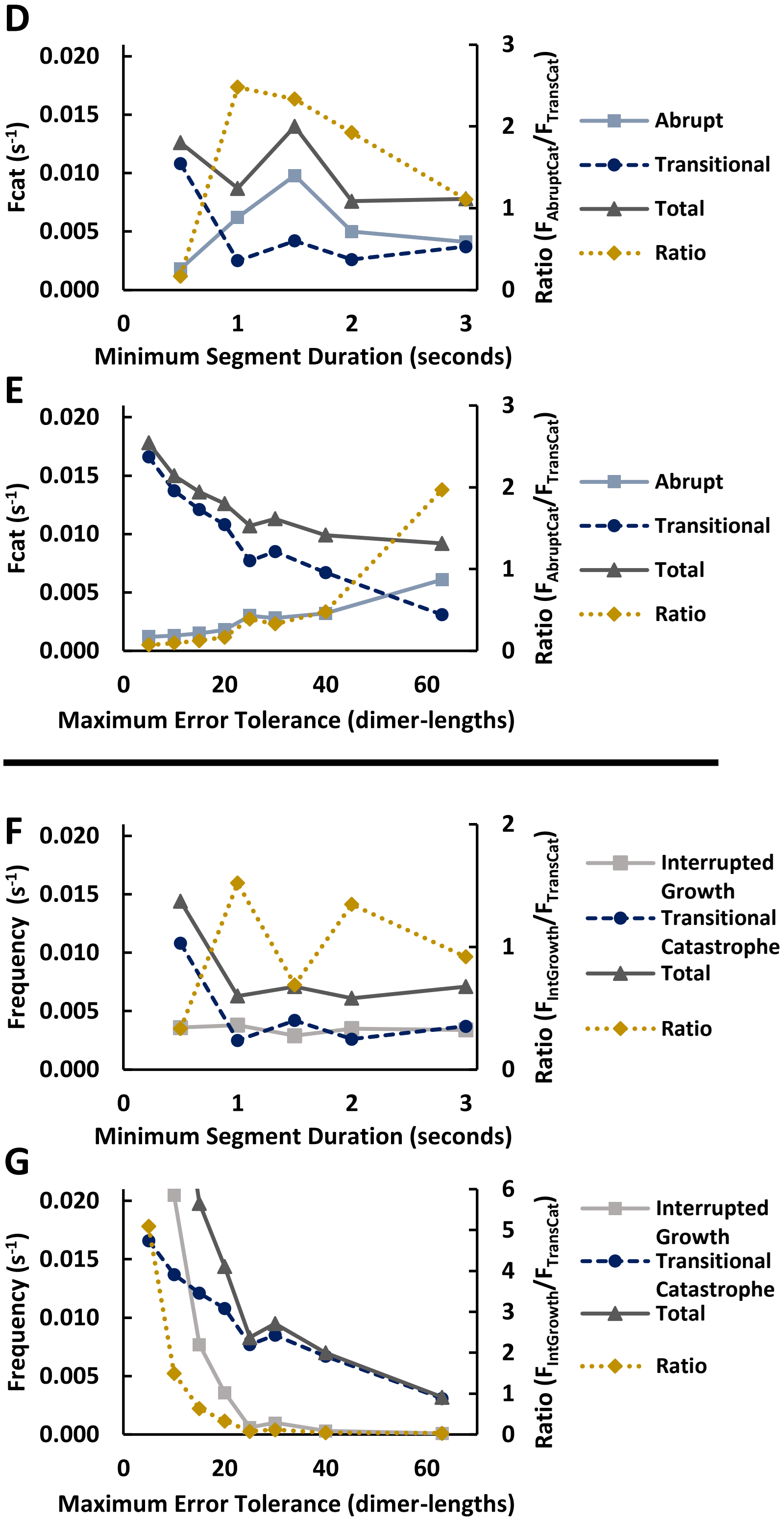


**Figure S2.10. Transition analysis across Minimum Segment Durations and Maximum Error Tolerances provide robust support for prevalence of transitional catastrophes in the *in vitro* control dataset.** These data are the results from analyzing the *in vitro* control dataset with STADIA using $k$=3 for positive and $k$=2 for negative slope segments for all user-input parameter combinations. **(A-C)** Transition frequencies across the full parameter space considered in **Supplemental Section 2**. The frequency of abrupt catastrophe ($F_{AbruptCat}$, A), frequency of transitional catastrophe ($F_{TransCat}$, B), and frequency of interrupted growth ($F_{IntGrowth}$, C) are reported in table form. The highlighted column (Minimum Segment Duration = 0.5 seconds) and row (Maximum Error Tolerance = 20 dimer-lengths) of each table correspond to data plotted in (D-G). **(D,E)** $F_{AbruptCat}$ and $F_{TransCat}$ for varying Minimum Segment Durations while holding the Maximum Error Tolerance constant at 20 dimer-lengths (D) and for varying Maximum Error Tolerances while holding the Minimum Segment Duration constant at 0.5 seconds (E). The raw frequencies are plotted corresponding to the left-hand y-axes; the ratio of $F_{AbruptCat}$ to $F_{TransCat}$ is plotted on the right-hand y-axes. Note that parameter combinations where $F_{AbruptCat}/F_{TransCat} < 1$ indicates $F_{TransCat} > F_{AbruptCat}$, which is consistent with the conclusions in the main text for the *in vitro* control data. **(F,G)** $F_{TransCat}$ and $F_{IntGrowth}$ for varying Minimum Segment Durations while holding the Maximum Error Tolerance constant at 20 dimer-lengths (F) and for varying Maximum Error Tolerances while holding the Minimum Segment Duration constant at 0.5 seconds (G). The raw frequencies are plotted corresponding to the left-hand y-axes; the ratio of $F_{IntGrowth}$ to $F_{TransCat}$ is plotted on the right-hand y-axes. For small Maximum Error Tolerance, please see the tables for the frequencies that are too high to be visible in (G).

**Figure S2.10 A,B,D,E – Observations and Interpretations**

**Observations:** For the *in vitro* control data, the total $F_{cat}$ (which equals $F_{AbruptCat} + F_{TransCat}$ ) is noisier compared to the *in silico* dataset, but still relatively steady over the Minimum Segment Durations tested (D). A steadier decrease in the total $F_{cat}$ can be seen across the Maximum Error Tolerances (E) tested compared to the sudden decrease followed by relatively steady values for the *in silico* data. For Maximum Error tolerance = 20 dimer-lengths (D), the ratio $F_{AbruptCat}/F_{TransCat}$ for the *in vitro* control data is less than 1 for Minimum Segment Durations = 0.5 seconds (consistent with the findings of in the main text), but is greater than 1 for Minimum Segment Durations greater than or equal to 1.0 seconds (in comparison, for the *in silico* data, this switch occurred at a slightly longer Minimum Segment Duration). For Minimum Segment Duration = 0.5 seconds (E), the ratio $F_{AbruptCat}/F_{TransCat}$ is consistently less than 1 for Maximum Error Tolerances less than or equal to 40 dimer-lengths (including Maximum Error Tolerance = 5 dimer-lengths, which is in contrast to the *in silico* results).

**Interpretations:** As was the case for the full resolution *in silico* dataset, observations and interpretations from analysis of the *in vitro* control dataset support the conclusion that catastrophes are more often transitional than abrupt. This conclusion is robust for the *in vitro* control dataset with Minimum Segment Durations less than 1.0 seconds and Maximum Error Tolerances less than or equal to 40 dimer-lengths (or 320 nm).

While subtle differences remain between the analysis of the *in vitro* control and *in silico* datasets, general patterns are consistent. Sufficiently small Minimum Segment Durations and Maximum Error Tolerances result in ratios of $F_{AbruptCat}$ to $F_{TransCat}$ less than 1 (i.e., $F_{TransCat} > F_{AbruptCat}$). The differences lie in where that relationship changes, but the other interpretations remain largely the same.

**Additional Discussion:** Increasing the Minimum Segment Duration makes detecting stutters more challenging, which explains the significant impact of changing the Minimum Segment Duration on $F_{TransCat}$ and $F_{AbruptCat}$. Increasing the Maximum Error Tolerance also affects the ability of STADIA to reliably detect stutters and therefore has an effect on the ratio. However, the effect from increasing the Maximum Error Tolerance does not drive the ratio above 1 until the tolerance is at 63 dimer-lengths (~0.5 microns). And so, the Maximum Error Tolerance is not as important for consistency of the transition analysis results as the Minimum Segment Duration.

**Figure S2.10 B,C,F,G – Observations and Interpretations**

**Observations:** As was seen for the *in silico* data, the frequency of growth-to-stutter transitions ($F_{IntGrowth} + F_{TransCat}$) appears to change more significantly with changes to either the Minimum Segment Duration (F) or Maximum Error Tolerance (G) compared to the overall $F_{cat}$. For varying the Minimum Segment Duration (F), the ratio of $F_{IntGrowth}$ to $F_{TransCat}$ is much less than 1 (i.e., $F_{IntGrowth} < F_{TransCat}$) for just the Minimum Segment Duration = 0.5 seconds. For Minimum Segment Durations > 0.5 seconds, the ratio oscillates around 1 (initial change in the ratio is due primarily to decrease in detection of transitional catastrophes). For the Minimum Segment Duration fixed at 0.5 seconds (G), the ratio of $F_{IntGrowth}$ to $F_{TransCat}$ is greater than 1 for Maximum Error Tolerances < 15 dimer-lengths. Similar to what was seen in the *in silico* dataset, but in contrast to the statement regarding the Minimum Segment Durations, the decrease in both the total frequency ($F_{IntGrowth} + F_{TransCat}$) and the ratio with increasing Maximum Error Tolerances can be attributed primarily to the large decrease in the frequency of interrupted growth.

**Interpretations:** The decrease in total frequency with increasing Minimum Segment Durations is not surprising. Both frequencies being considered involve the detection of stutters which becomes more difficult for STADIA at longer Minimum Segment Durations. The consistent pattern of $F_{IntGrowth} \approx F_{TransCat}$ for Minimum Segment Durations > 0.5 seconds is interesting as it suggests general stability in the relationship between $F_{IntGrowth}$ and $F_{TransCat}$ with respect to the segmentation parameters. The large decrease in $F_{IntGrowth}$ with increasing Maximum Error Tolerance suggests that detection of stutters in growing MTs is much more sensitive to the Maximum Error Tolerance than is detection of stutters preceding catastrophes.

The differences in sensitivity for each of the growth-to-stutter frequencies to Minimum Segment Duration and Maximum Error Tolerance are notable. Specifically (similar to the *in silico* data), the Maximum Error Tolerance value has a much greater effect on the detection of stutters within growth phases, whereas the Minimum Segment Duration value has a somewhat greater effect on the detection of stutters preceding catastrophes.

**A**

| Frequency of ABRUPT CATASTROPHE | | | | | | |
|---|---|---|---|---|---|---|
| Max Error Tolerance (dimer-lengths) | Minimum Segment Duration (s) | | | | | |
| | 0.3 | 0.5 | 1.0 | 1.5 | 2.0 | 3.0 |
| 5 | Not Calculated for *In Vitro* data | 0.0379 | 0.0412 | 0.0160 | 0.0078 | 0.0112 |
| 10 | | 0.0176 | 0.0178 | 0.0165 | 0.0109 | 0.0059 |
| 15 | | 0.0087 | 0.0082 | 0.0089 | 0.0083 | 0.0050 |
| 20 | | 0.0049 | 0.0057 | 0.0055 | 0.0074 | 0.0050 |
| 25 | | 0.0037 | 0.0041 | 0.0042 | 0.0048 | 0.0024 |
| 30 | | 0.0042 | 0.0046 | 0.0049 | 0.0042 | 0.0025 |
| 40 | | 0.0034 | 0.0033 | 0.0035 | 0.0059 | 0.0055 |
| 63 | | 0.0023 | 0.0021 | 0.0054 | 0.0050 | 0.0054 |

**B**

| Frequency of TRANSITIONAL CATASTROPHE | | | | | | |
|---|---|---|---|---|---|---|
| Max Error Tolerance (dimer-lengths) | Minimum Segment Duration (s) | | | | | |
| | 0.3 | 0.5 | 1.0 | 1.5 | 2.0 | 3.0 |
| 5 | Not Calculated for *In Vitro* data | 0.0148 | 0.0144 | 0.0124 | 0.0088 | 0.0053 |
| 10 | | 0.0074 | 0.0063 | 0.0033 | 0.0032 | 0.0029 |
| 15 | | 0.0046 | 0.0040 | 0.0029 | 0.0024 | 0.0017 |
| 20 | | 0.0034 | 0.0024 | 0.0018 | 0.0022 | 0.0015 |
| 25 | | 0.0032 | 0.0023 | 0.0026 | 0.0044 | 0.0030 |
| 30 | | 0.0017 | 0.0019 | 0.0019 | 0.0028 | 0.0025 |
| 40 | | 0.0011 | 0.0009 | 0.0010 | 0.0008 | 0.0008 |
| 63 | | 0.0003 | 0.0003 | 0.0000 | 0.0000 | 0.0000 |

**C**

| Frequency of INTERRUPTED GROWTH | | | | | | |
|---|---|---|---|---|---|---|
| Max Error Tolerance (dimer-lengths) | Minimum Segment Duration (s) | | | | | |
| | 0.3 | 0.5 | 1.0 | 1.5 | 2.0 | 3.0 |
| 5 | Not Calculated for *In Vitro* data | 0.1129 | 0.0720 | 0.0760 | 0.0460 | 0.0279 |
| 10 | | 0.0321 | 0.0291 | 0.0204 | 0.0162 | 0.0102 |
| 15 | | 0.0137 | 0.0124 | 0.0100 | 0.0089 | 0.0061 |
| 20 | | 0.0088 | 0.0075 | 0.0039 | 0.0071 | 0.0056 |
| 25 | | 0.0050 | 0.0030 | 0.0022 | 0.0044 | 0.0013 |
| 30 | | 0.0030 | 0.0018 | 0.0016 | 0.0042 | 0.0011 |
| 40 | | 0.0020 | 0.0014 | 0.0010 | 0.0020 | 0.0008 |
| 63 | | 0.0008 | 0.0008 | 0.0007 | 0.0020 | 0.0011 |

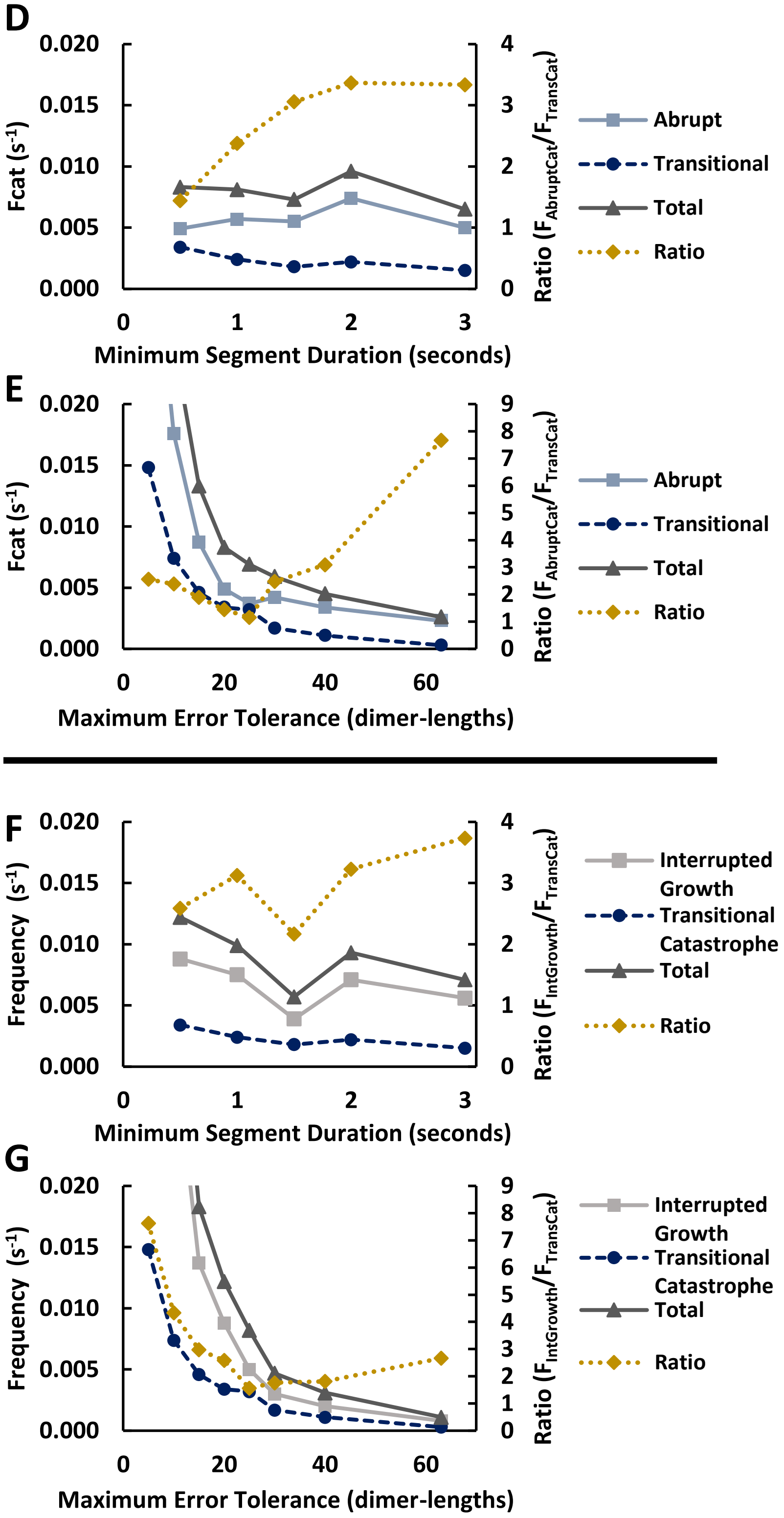


**Figure S2.11. Transition analysis across Minimum Segment Durations and Maximum Error Tolerances provide robust support for reduced transitional catastrophe frequency and increased interrupted growth frequency in the *in vitro* CLASP2γ dataset.** These data are the results from analyzing the *in vitro* CLASP2γ dataset with STADIA using *k*=3 for positive and *k*=2 for negative slope segments for all user-input parameter combinations. **(A-C)** Transition frequencies across the full parameter space considered in **Supplemental Section 2**. The frequency of abrupt catastrophe ($F_{AbruptCat}$, A), frequency of transitional catastrophe ($F_{TransCat}$, B), and frequency of interrupted growth ($F_{IntGrowth}$, C) are reported in table form. The highlighted column (Minimum Segment Duration = 0.5 seconds) and row (Maximum Error Tolerance = 20 dimer-lengths) of each table correspond to data plotted in (D-G). **(D,E)** $F_{AbruptCat}$ and $F_{TransCat}$ for varying Minimum Segment Durations while holding the Maximum Error Tolerance constant at 20 dimer-lengths (D) and for varying Maximum Error Tolerances while holding the Minimum Segment Duration constant at 0.5 seconds (E). The raw frequencies are plotted corresponding to the left-hand y-axes; the ratio of $F_{AbruptCat}$ to $F_{TransCat}$ is plotted on the right-hand y-axes. Note that parameter combinations where $F_{AbruptCat}/F_{TransCat} > 1$ indicates $F_{TransCat} < F_{AbruptCat}$, which is consistent with the conclusions in the main text for the *in vitro* CLASP2γ data. **(F,G)** $F_{TransCat}$ and $F_{IntGrowth}$ for varying Minimum Segment Durations while holding the Maximum Error Tolerance constant at 20 dimer-lengths (F) and for varying Maximum Error Tolerances while holding the Minimum Segment Duration constant at 0.5 seconds (G). The raw frequencies are plotted corresponding to the left-hand y-axes; the ratio of $F_{IntGrowth}$ to $F_{TransCat}$ is plotted on the right-hand y-axes. For small Maximum Error Tolerances, please see the tables for the frequencies that are which are too high to be visible in (E,G).

**Figure S2.11 A,B,D,E – Observations and Interpretations**

**Observations:** The patterns observed for analysis of the *in vitro* CLASP2γ dataset with varying Minimum Segment Durations are remarkably similar to the results for the *in vitro* control dataset: total $F_{cat}$ is relatively stable and the ratio $F_{AbruptCat}/F_{TransCat}$ is greater than 1 (i.e., $F_{AbruptCat} > F_{TransCat}$) for Minimum Segment Durations > 0.5 seconds. The obvious difference, however, is that $F_{AbruptCat}/F_{TransCat} > 1$ for Minimum Segment Duration = 0.5 seconds in the CLASP2γ data but not in the control data (see Interpretations). In complete contrast to the *in vitro* control data, $F_{AbruptCat}/F_{TransCat} > 1$ with CLASP2γ regardless of the Maximum Error Tolerance used.

**Interpretations:** The above observations support the relationship found in the main text that when CLASP2γ is present, STADIA finds that *in vitro* MTs undergo abrupt catastrophes more often than transitional catastrophes (see **Figure 6A**). This conclusion is robust regardless of the segmentation parameters tested.

**Figure S2.11 B,C,F,G – Observations and Interpretations**

**Observations:** Unlike what was seen in the analysis of the *in vitro* control dataset, $F_{IntGrowth} > F_{TransCat}$ with CLASP2γ for all data plotted in (F) and (G).

**Interpretations:** The consistent pattern of $F_{IntGrowth} > F_{TransCat}$ for all Minimum Segment Durations and Maximum Error Tolerances is of particular note (see **Figure S2.12** for more on this). The large decrease in $F_{IntGrowth}$ with increasing Maximum Error Tolerance suggests that detection of stutters in growing MTs is much more sensitive to the Maximum Error Tolerance than is detection of stutters preceding catastrophes.

The fact that $F_{IntGrowth} > F_{TransCat}$ regardless of segmentation parameters supports the conclusion the CLASP2γ promotes growth of stuttering MTs.

**A** *In Vitro* (control)

| Frequency of TRANSITIONAL CATASTROPHE | | | | | | |
|---|---|---|---|---|---|---|
| Max Error Tolerance (dimer-lengths) | Minimum Segment Duration (s) | | | | | |
| | 0.3 | 0.5 | 1.0 | 1.5 | 2.0 | 3.0 |
| 5 | Not Calculated for *In Vitro* data | 0.0166 | 0.0069 | 0.0061 | 0.0058 | 0.0059 |
| 10 | | 0.0137 | 0.0049 | 0.0043 | 0.0045 | 0.0052 |
| 15 | | 0.0121 | 0.0032 | 0.0051 | 0.0033 | 0.0043 |
| 20 | | 0.0108 | 0.0025 | 0.0042 | 0.0026 | 0.0037 |
| 25 | | 0.0077 | 0.0002 | 0.0072 | 0.0019 | 0.0030 |
| 30 | | 0.0085 | 0.0020 | 0.0017 | 0.0018 | 0.0026 |
| 40 | | 0.0067 | 0.0015 | 0.0014 | 0.0013 | 0.0019 |
| 63 | | 0.0031 | 0.0006 | 0.0005 | 0.0005 | 0.0007 |

**C** *In Vitro* (CLASP2γ)

| Frequency of TRANSITIONAL CATASTROPHE | | | | | | |
|---|---|---|---|---|---|---|
| Max Error Tolerance (dimer-lengths) | Minimum Segment Duration (s) | | | | | |
| | 0.3 | 0.5 | 1.0 | 1.5 | 2.0 | 3.0 |
| 5 | Not Calculated for *In Vitro* data | 0.0148 | 0.0144 | 0.0124 | 0.0088 | 0.0053 |
| 10 | | 0.0074 | 0.0063 | 0.0033 | 0.0032 | 0.0029 |
| 15 | | 0.0046 | 0.0040 | 0.0029 | 0.0024 | 0.0017 |
| 20 | | 0.0034 | 0.0024 | 0.0018 | 0.0022 | 0.0015 |
| 25 | | 0.0032 | 0.0023 | 0.0026 | 0.0044 | 0.0030 |
| 30 | | 0.0017 | 0.0019 | 0.0019 | 0.0028 | 0.0025 |
| 40 | | 0.0011 | 0.0009 | 0.0010 | 0.0008 | 0.0008 |
| 63 | | 0.0003 | 0.0003 | 0.0000 | 0.0000 | 0.0000 |

**B**

| Frequency of INTERRUPTED GROWTH | | | | | | |
|---|---|---|---|---|---|---|
| Max Error Tolerance (dimer-lengths) | Minimum Segment Duration (s) | | | | | |
| | 0.3 | 0.5 | 1.0 | 1.5 | 2.0 | 3.0 |
| 5 | Not Calculated for *In Vitro* data | 0.0846 | 0.0822 | 0.0739 | 0.0495 | 0.0325 |
| 10 | | 0.0205 | 0.0202 | 0.0197 | 0.0178 | 0.0156 |
| 15 | | 0.0077 | 0.0078 | 0.0058 | 0.0072 | 0.0064 |
| 20 | | 0.0036 | 0.0038 | 0.0029 | 0.0035 | 0.0034 |
| 25 | | 0.0006 | 0.0007 | 0.0006 | 0.0020 | 0.0019 |
| 30 | | 0.0010 | 0.0010 | 0.0010 | 0.0009 | 0.0009 |
| 40 | | 0.0003 | 0.0003 | 0.0002 | 0.0003 | 0.0002 |
| 63 | | 0.0001 | 0.0001 | 0.0001 | 0.0001 | 0.0000 |

**D**

| Frequency of INTERRUPTED GROWTH | | | | | | |
|---|---|---|---|---|---|---|
| Max Error Tolerance (dimer-lengths) | Minimum Segment Duration (s) | | | | | |
| | 0.3 | 0.5 | 1.0 | 1.5 | 2.0 | 3.0 |
| 5 | Not Calculated for *In Vitro* data | 0.1129 | 0.0720 | 0.0760 | 0.0460 | 0.0279 |
| 10 | | 0.0321 | 0.0291 | 0.0204 | 0.0162 | 0.0102 |
| 15 | | 0.0137 | 0.0124 | 0.0100 | 0.0089 | 0.0061 |
| 20 | | 0.0088 | 0.0075 | 0.0039 | 0.0071 | 0.0056 |
| 25 | | 0.0050 | 0.0030 | 0.0022 | 0.0044 | 0.0013 |
| 30 | | 0.0030 | 0.0018 | 0.0016 | 0.0042 | 0.0011 |
| 40 | | 0.0020 | 0.0014 | 0.0010 | 0.0020 | 0.0008 |
| 63 | | 0.0008 | 0.0008 | 0.0007 | 0.0020 | 0.0011 |

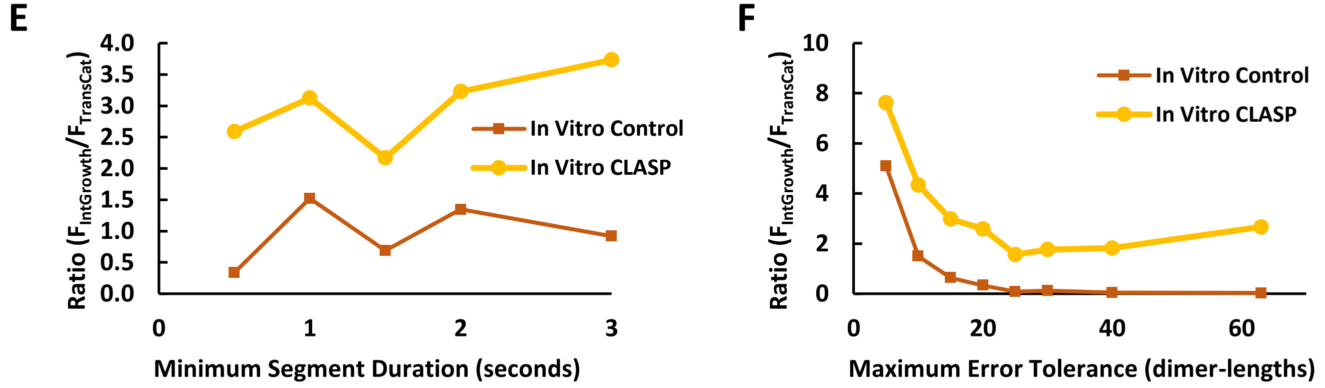


**Figure S2.12. Comparison of transition analysis results across Minimum Segment Durations and Maximum Error Tolerances supports conclusion that CLASP2γ suppress transitional catastrophes and promotes interrupted growths. (A-D)** Frequencies of transitional catastrophes ($F_{TransCat}$; A,C) and interrupted growths ($F_{IntGrowth}$; B,D) for the *in vitro* control data (A,B) and the *in vitro* CLASP2γ data (C,D). Note that this data is identical to the transition data in **Figure S2.10 B,C** (*in vitro* control) and **Figure S2.11 B,C** (*in vitro* CLASP2γ). **(E,F)** Ratios of $F_{IntGrowth}$ to $F_{TransCat}$ for varying Minimum Segment Durations while holding the Maximum Error Tolerance constant at 20 dimer-lengths (E) and varying Maximum Error Tolerances while holding the Minimum Segment Duration constant at 0.5 seconds (F).

**Figure S2.12 – Observations and Interpretations**

**Observations:** For all combinations of Minimum Segment Duration and Maximum Error Tolerance plotted in (E,F), the ratio of $F_{IntGrowth}$ to $F_{TransCat}$ is greater in the *in vitro* CLASP2γ dataset (yellow) than in the *in vitro* control dataset (orange).

**Interpretations:** The observation above demonstrates that the conclusion that CLASP2γ suppresses catastrophe by promoting growth of stuttering MTs is robust with respect to the STADIA segmentation parameters.

### Supplemental Section 3:
#### Data acquisition rate sensitivity analysis

**<u>Overview</u>:** The goal of the analyses presented the last section (2) was to test the robustness of the main conclusions of the manuscript to variation in the values of the user-input STADIA segmentation parameters "Minimum Segment Duration" and "Maximum Error Tolerance". In this section, we examine sensitivity to the temporal resolution of the inputted length-history data itself.

In this section, the conclusions being considered are as follows: **(1)** MTs exhibit more behaviors than just growth and shortening, with stutters being distinguishable behaviors that are prevalent throughout length-history data, and **(2)** transitional catastrophes are more frequent than abrupt catastrophes. The table of contents below directs readers to the figures related to each of the above conclusions. In addition, the next page contains information to assist readers in understanding and interpreting the analyses performed.

**<u>Table of Contents</u>**


**Supplemental Section 3: Parameter sweep analysis of dimer-scale *in silico* datasets with varying data acquisition rates**

**1.** Analysis of *in silico* data with varying Data Acquisition Time Steps and Maximum Error Tolerances using a consistent **Minimum Segment Duration of 0.5 seconds**

- **a.** Gap statistic plots – **Figures S3.1** (positive slope segments), **S3.2** (negative slope segments)
  *Useful for conclusion (1)*
- **b.** Cluster profiles of positive and negative slope segment data – **Figure S3.3 (A)**
  *Useful for conclusion (1)*
- **c.** Labeled length-history plots – **Figure S3.3 (B)**
  *Useful for conclusions (1) and (2)*

**2.** Analysis of *in silico* data with varying Data Acquisition Time Steps and Maximum Error Tolerances using a consistent **Minimum Segment Duration of 1.0 seconds**

- **a.** Gap statistic plots – **Figures S3.4** (positive slope segments), **S3.5** (negative slope segments)
  *Useful for conclusion (1)*
- **b.** Cluster profiles of positive and negative slope segment data – **Figure S3.6 (A)**
  *Useful for conclusion (1)*
- **c.** Labeled length-history plots – **Figure S3.6 (B)**
  *Useful for conclusions (1) and (2)*

**3.** Analysis of *in silico* data with varying Data Acquisition Time Steps and Maximum Error Tolerances using a consistent **Minimum Segment Duration of 3.0 seconds**

- **a.** Gap statistic plots – **Figures S3.7** (positive slope segments), **S3.8** (negative slope segments)
  *Useful for conclusion (1)*
- **b.** Cluster profiles of positive and negative slope segment data – **Figure S3.9 (A)**
  *Useful for conclusion (1)*
- **c.** Labeled length-history plots – **Figure S3.9 (B)**
  *Useful for conclusions (1) and (2)*


**Note to Readers:** As noted in the introduction to Supplemental Section 2, each figure is accompanied by a legend as well as explanatory text that contains observations, interpretations, and in some cases additional discussion. For most of the figures, some of the accompanying text is on the page following the figure. Since parameter sweeps are repetitive by nature, we use similar language in the text accompanying many figures. Despite some apparent redundancy, we use this approach to make each figure independently interpretable.

#### **Introduction to Section 3**, Data acquisition rate sensitivity analysis

In Section 2, we explored the impact of changing STADIA parameters on all three datasets used in the main text. In this section, we examine the effect of changing the data acquisition rate (“frame” rate). This analysis uses the *in silico* data to allow for varying the data acquisition rate across a wide range of values including rates faster than the *in vitro* frame rate. We achieve this variation by imposing various fixed data acquisition rates on the original full resolution dataset. We then compare the results from STADIA analysis of these reduced resolution datasets to those obtained with the full resolution dataset.

In the following figures, “Full Resolution Data” refers to the raw simulation output, which includes a data point for every dimer-scale biochemical event. For the input parameters used in the simulations in this manuscript, approximately 1650 events occurred per second. See Methods for more information. The *in silico* data used in the main text and in **Supplemental Sections 1 & 2** was also the full resolution data.

The setup of the following analyses is similar to the parameter sweep analyses presented in Section 2. However, an additional complexity is introduced by varying the Data Acquisition Time Step (‘data acquisition rate in frames per second’ = 1/‘Data Acquisition Time Step in seconds’), in addition to varying the two STADIA parameters examined in Section 2. As a result of the need to examine three variables, the organization of figures in Section 3 differs from Section 2. Specifically, here in Section 3, there are three sets of three figures. Each set (i.e., **Figures S3 1-3, 4-6, or 7-9**) corresponds to a particular Minimum Segment Duration (0.5, 1.0, or 3.0 seconds, respectively). Within each set, the three figures are positive slope gap statistic plots (**Figures S3.1, S3.4, S3.7**), negative slope gap statistic plots (**Figures S3.2, S3.5, S3.8**), and cluster profiles for the positive and negative slope segments along with labeled length-history plots (**Figures S3.3, S3.6, S3.9**). Within the gap statistic figures, the columns correspond to varying values of the Data Acquisition Time Step and the rows correspond to varying values of the Maximum Error Tolerance. Within the cluster profile and length-history figures, the Maximum Error Tolerance is fixed at 20 dimer-lengths and the rows now correspond to varying Data Acquisition Time Steps.

With the above difference in mind, the general question being addressed with this set of figures is similar to that addressed in **Supplemental Section 2**. Here, do the main conclusions of this manuscript change when the temporal resolution of the inputted length-history data is varied?

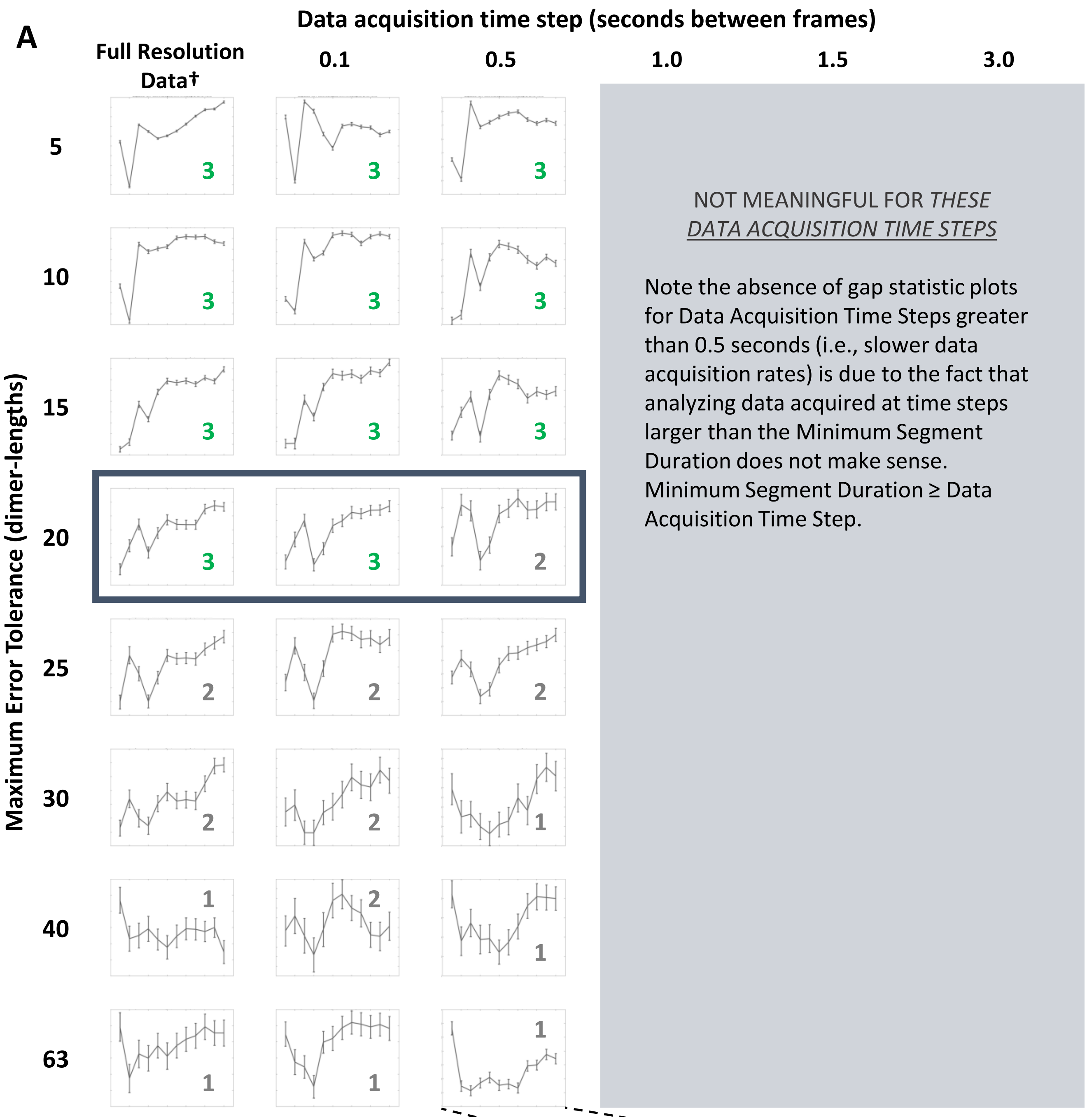


**Figure S3.1. Gap statistic plots for *in silico* data (Positive Slope Segments) support conclusion of multiple shortening behaviors at relevant spatial scales (using a Minimum Segment Duration of 0.5 seconds) regardless of data acquisition rate. (A)** The gap statistic plots presented are for the positive slope segments and are the result of analyzing the *in silico* dataset at varying data acquisition rates using the Diagnostic Mode of STADIA with constant Minimum Segment Duration of 0.5 seconds and varying Maximum Error Tolerance (row headings) and the Data Acquisition Time Step (column headings). Each gap statistic plot is labeled with the optimal *k*-value suggested (green is used to indicate agreement with the results for this dataset in main text; gray indicates parameter combinations resulting in different *k*-values. The **dark blue** box indicates the parameter space for which cluster profiles and labeled length-history plots can be found in **Figure S3.3**. **(B)** A representative gap statistic plot (bottom right) shows the axes for each plot in (A). The x-axis (*k*-value) range is the same for all plots. The y-axis (Gap-Value) has differing ranges (not shown) for each plot, but the specific numerical values of the gap statistic are not relevant to interpreting the plots because identification of the optimal *k*-value is based on local maxima within each gap statistic plot. In other words, the pertinent information is the relationship between the values of the gap statistic at different *k*-values within each plot, not the values themselves. † The **full resolution** *in silico* data have temporal resolution of one output per dimer-scale biochemical event (see Methods).

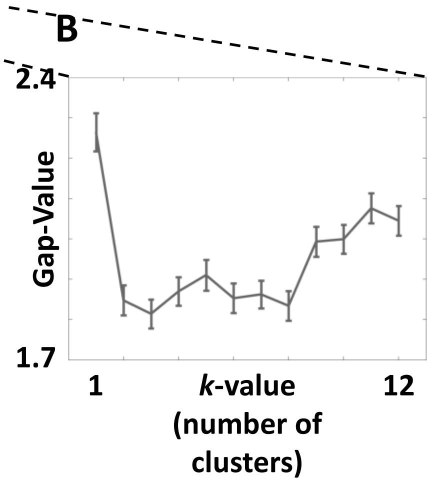

**Figure S3.1 – Observations and Interpretations**

**Observations:** When using a Minimum Segment Duration of 0.5 seconds, the results of analyzing the positive slope line segment data from piecewise linear approximation of the *in silico* data captured at 0.1 and 0.5 second Data Acquisition Time Steps are very similar to the results from analysis of the full resolution data. The general pattern of $k$=3 at sufficiently small Maximum Error Tolerances and a diminishing number of clusters with increasing Maximum Error Tolerance is upheld across the Data Acquisition Time Steps considered.

**Interpretations:** The stability of the $k$-value suggested by the gap statistic plots across the Data Acquisition Time Steps is an encouraging result for the conclusion that stutters exist and are not simply a result of analyzing data at fine resolution. The sensitivity to changes in the user-input parameter Maximum Error Tolerance is also generally uniform across the Data Acquisition Time Steps. The results in this figure are especially relevant because the analysis of the *in silico* data in the main text of this manuscript was conducted using a Minimum Segment Duration of 0.5 seconds (as was done in this figure).

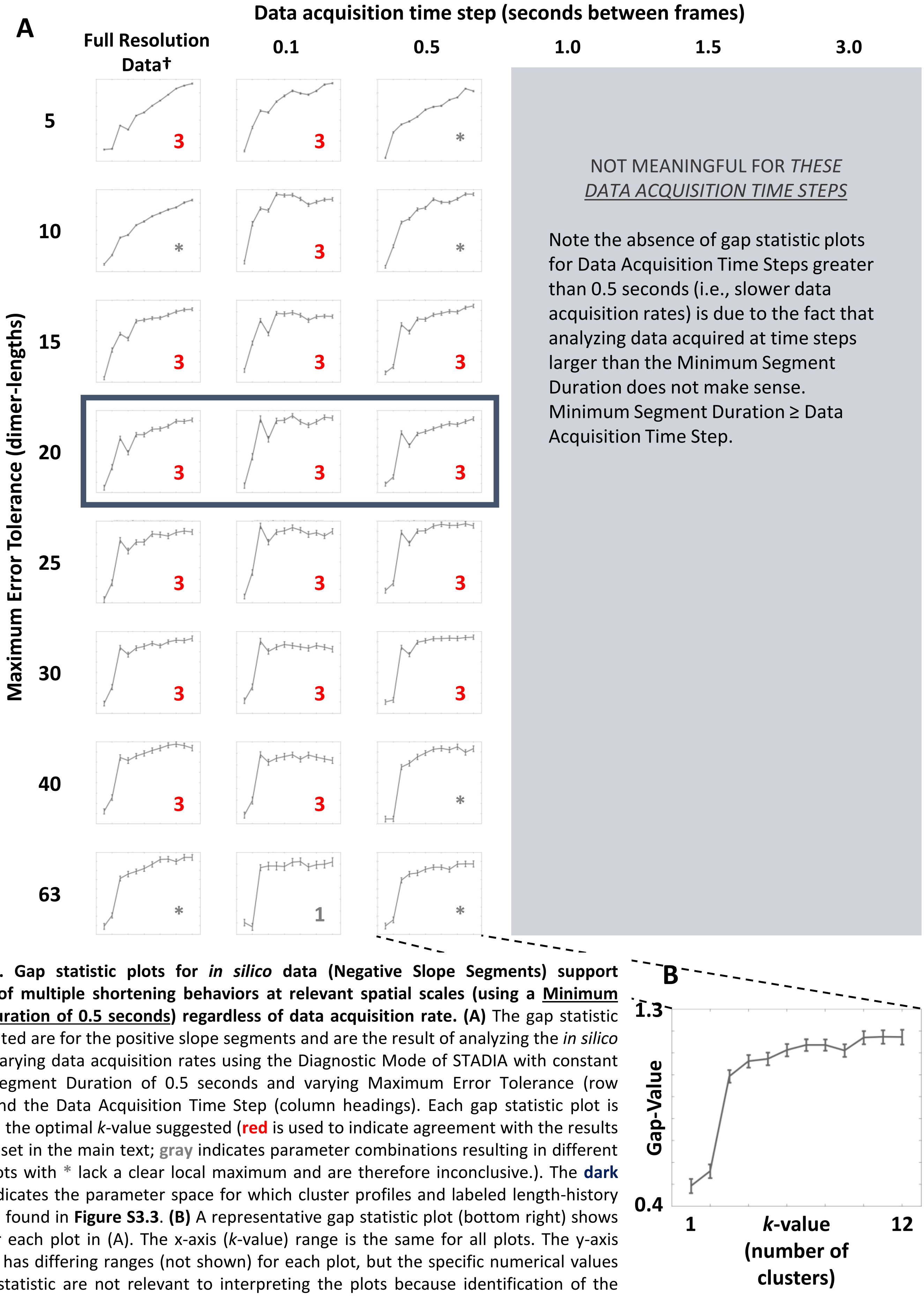


**Figure S3.2. Gap statistic plots for *in silico* data (Negative Slope Segments) support conclusion of multiple shortening behaviors at relevant spatial scales (using a Minimum Segment Duration of 0.5 seconds) regardless of data acquisition rate. (A)** The gap statistic plots presented are for the positive slope segments and are the result of analyzing the *in silico* dataset at varying data acquisition rates using the Diagnostic Mode of STADIA with constant Minimum Segment Duration of 0.5 seconds and varying Maximum Error Tolerance (row headings) and the Data Acquisition Time Step (column headings). Each gap statistic plot is labeled with the optimal *k*-value suggested (**red** is used to indicate agreement with the results for this dataset in the main text; **gray** indicates parameter combinations resulting in different *k*-values. Plots with * lack a clear local maximum and are therefore inconclusive.). The **dark blue** box indicates the parameter space for which cluster profiles and labeled length-history plots can be found in **Figure S3.3**. **(B)** A representative gap statistic plot (bottom right) shows the axes for each plot in (A). The x-axis (*k*-value) range is the same for all plots. The y-axis (Gap-Value) has differing ranges (not shown) for each plot, but the specific numerical values of the gap statistic are not relevant to interpreting the plots because identification of the optimal *k*-value is based on local maxima within each gap statistic plot. In other words, the pertinent information is the relationship between the values of the gap statistic at different *k*-values within each plot, not the values themselves. † The **full resolution** *in silico* data have temporal resolution of one output per dimer-scale biochemical event (see Methods).

**Figure S3.2 – Observations and Interpretations**

**Observations:** When using a Minimum Segment Duration of 0.5 seconds, the results of analyzing the negative slope line segment data from piecewise linear approximation of the *in silico* data captured at 0.1 and 0.5 second Data Acquisition Time Steps are very similar to the results from analysis of the full resolution data. Across the Data Acquisition Time Steps, the peak at *k*=3 is most prominent for Maximum Error Tolerances of 15 to 30 dimer-lengths. Above and below that range of Maximum Error Tolerances, there is ambiguity in the suggested *k*-value and somewhat greater variability across the Data Acquisition Time Steps.

**Interpretations:** Similar to the positive slope segments, the stability of the *k*-value suggested by the gap statistic plots across the Data Acquisition Time Steps for the negative slope segments is an encouraging result for the conclusion that stutters exist and are not simply a result of analyzing data at fine resolution. The sensitivity to changes in the user-input parameter Maximum Error Tolerance is also generally uniform across the Data Acquisition Time Steps. These results are especially relevant because analysis of the *in silico* data in the main text of this manuscript was conducted using a Minimum Segment Duration of 0.5 seconds (as was done in this figure).

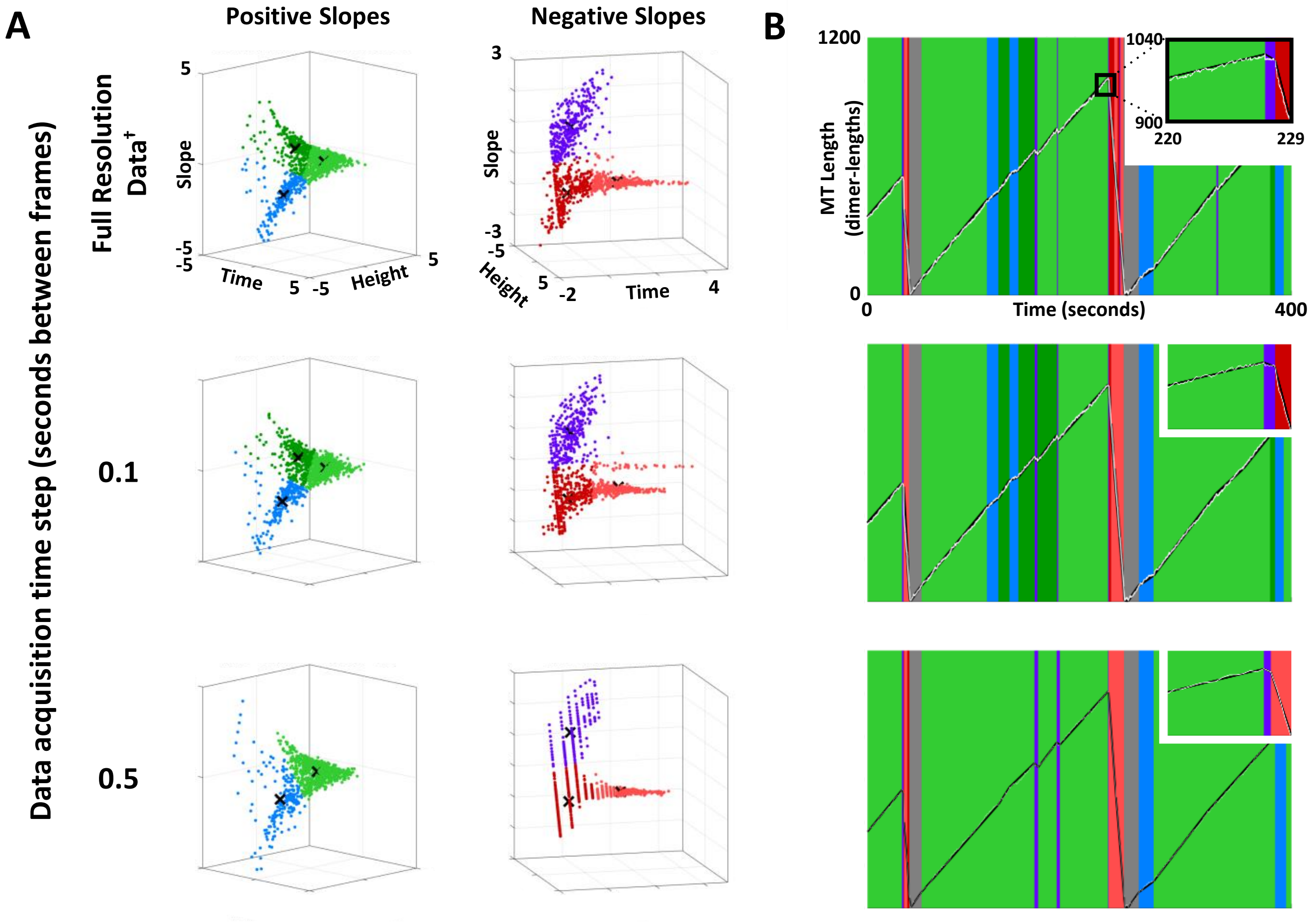


**Figure S3.3. Cluster profiles of positive and negative slope segment data (A) with representative labeled length-history plots (B).** The cluster plots and labeled length-history plots represent classification results from using STADIA in Automated Mode with the *k*-values indicated by the corresponding gap statistic plots in **Figures S3.1** and **S3.2**. Data acquisition rates are indicated by row headers. The Minimum Segment Duration was set to 0.5 seconds and the Maximum Error Tolerance was set to 20 dimer-lengths. **†** The **full resolution** *in silico* data have temporal resolution of one output per dimer-scale biochemical event (see Methods).

**Figure S3.3 A – Observations and Interpretations**

**Observations:** The cluster profiles for the positive and negative slope segment data maintain the same general shape regardless of the Data Acquisition Time Step when the data are analyzed using a Minimum Segment Duration of 0.5 seconds and Maximum Error Tolerance of 20 subunits. A notable difference across the varying Data Acquisition Time Steps, however, is that the overall density of data points decreases with increasing Data Acquisition Time Steps (i.e., moving down the rows).

**Interpretations:** The relative stability of the shape of the cluster profiles for the positive and negative slope segments with varying Data Acquisition Time Steps bolsters the conclusions about the number of clusters drawn from the gap statistic plots in **Figures S3.1** and **S3.2** (recall that the gap statistic drives the decision for the optimal *k*-value, but cluster profiles are also used to inform the *k*-value).

**Additional Discussion:** The appearance of a loss of density (particularly in the shortening segments) is due, in part, to the data points collapsing onto lines that correspond to multiples of the Data Acquisition Time Step. Additionally, with the longer Data Acquisition Time Steps, more loss of density occurs in the stutters and brief growth and shortening than in the sustained growth and shortening (there are not as many short segments).

**Figure S3.3 B – Observations and Interpretations**

**Observations:** For increasing values of the Data Acquisition Time Step, fewer stutters are detected throughout the plotted region of length-history data.

Additionally, the zoomed-in portraits in (B) provide an example of transitional catastrophe detection being somewhat reliant on the Data Acquisition Time Step. STADIA is able to correctly identify this catastrophe as transitional regardless of the Data Acquisition Time Step, but by eye the catastrophe appears to become somewhat less transitional as the Data Acquisition Time Step increases.

**Interpretations:** This example illustrates that STADIA's ability to detect individual stutter segments can depend on the temporal resolution of the data.

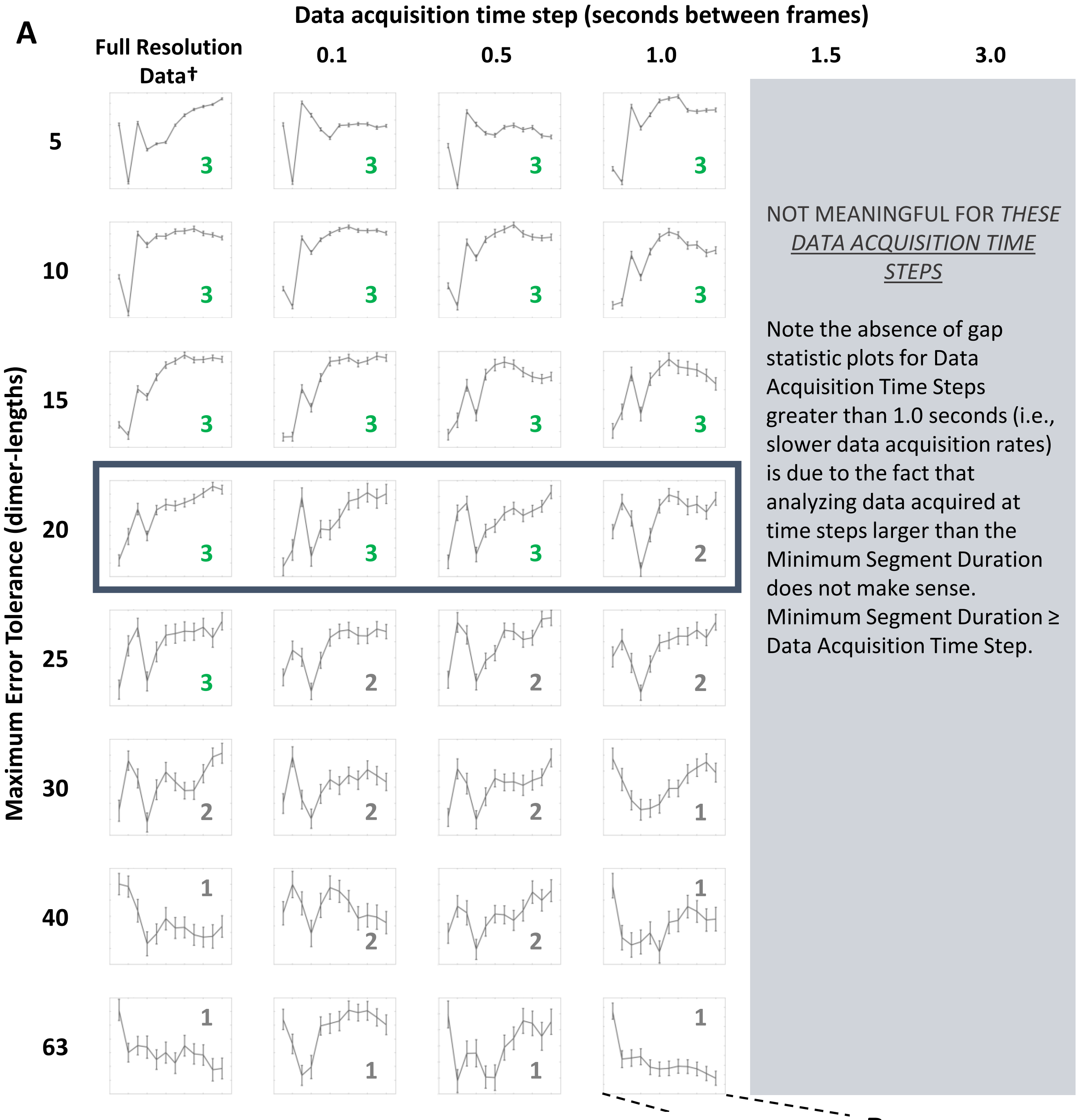


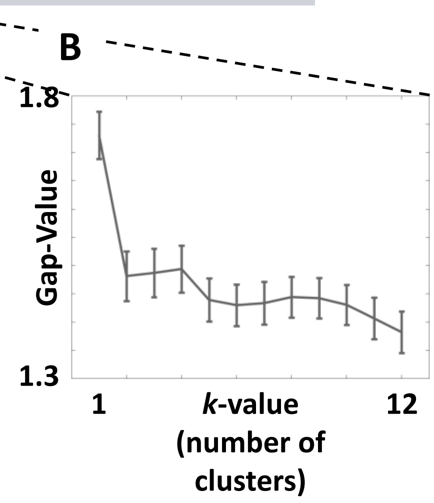


**Figure S3.4. Gap statistic plots for *in silico* data (Positive Slope Segments) support conclusion of multiple shortening behaviors at relevant spatial scales (using a <u>Minimum Segment Duration of 1.0 seconds</u>) regardless of data acquisition rate. (A)** The gap statistic plots presented are for the positive slope segments and are the result of analyzing the *in silico* dataset at varying data acquisition rates using the Diagnostic Mode of STADIA with constant Minimum Segment Duration of 1.0 seconds and varying Maximum Error Tolerance (row headings) and the Data Acquisition Time Step (column headings). Each gap statistic plot is labeled with the optimal *k*-value suggested (**green** is used to indicate agreement with the results for this dataset in the main text; **gray** indicates parameter combinations resulting in different *k*-values.). The **dark blue** box indicates the parameter space for which cluster profiles and labeled length-history plots can be found in **Figure S3.6. (B)** A representative gap statistic plot (bottom right) shows the axes for each plot in (A). The x-axis (*k*-value) range is the same for all plots. The y-axis (Gap-Value) has differing ranges (not shown) for each plot, but the specific numerical values of the gap statistic are not relevant to interpreting the plots because identification of the optimal *k*-value is based on local maxima within each gap statistic plot. In other words, the pertinent information is the relationship between the values of the gap statistic at different *k*-values within each plot, not the values themselves.

† The **full resolution** *in silico* data have temporal resolution of one output per dimer-scale biochemical event (see Methods).

**Figure S3.4 – Observations and Interpretations**

**Observations:** When using a Minimum Segment Duration of 1.0 seconds, the results of analyzing the positive slope line segment data from piecewise linear approximation of the *in silico* data captured at 0.1, 0.5, and 1.0 second Data Acquisition Time Steps are very similar to the results from analysis of the full resolution data. The general pattern of $k$=3 at sufficiently small Maximum Error Tolerances and a diminishing number of clusters with increasing Maximum Error Tolerance is upheld across the Data Acquisition Time Steps considered.

**Interpretations:** The stability of the $k$-value suggested by the gap statistic plots across the Data Acquisition Time Steps is an encouraging result for the conclusion that stutters exist and are not simply a result of analyzing data at fine resolution. The sensitivity to changes in the user-input parameter Maximum Error Tolerance is also generally uniform across the Data Acquisition Time Steps. From the parameter sweep results for the full resolution data in **Supplemental Section 2**, however, we know that even when stutters are detected as a distinct cluster, the detection of individual stutter segments starts to decrease when using a Minimum Segment Duration of 1.0 seconds (**Figures S2.3, S2.4, S2.9**).

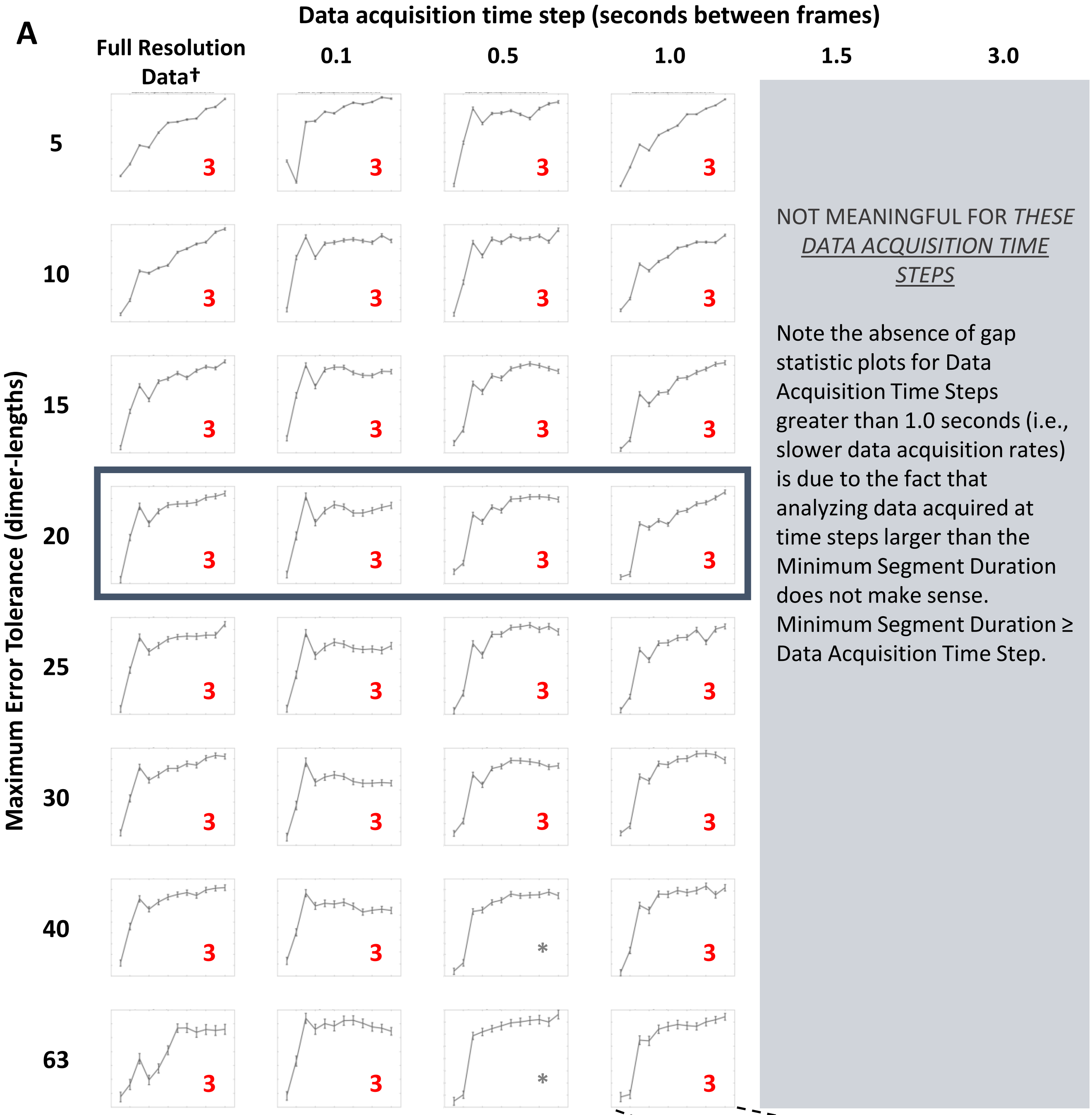


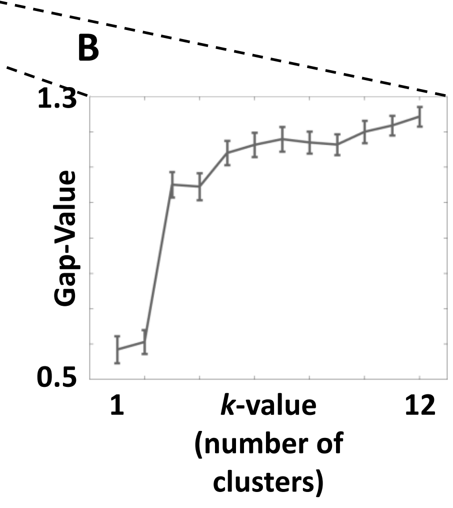


**Figure S3.5. Gap statistic plots for *in silico* data (Negative Slope Segments) support conclusion of multiple shortening behaviors at relevant spatial scales (using a Minimum Segment Duration of 1.0 seconds) regardless of data acquisition rate. (A)** The gap statistic plots presented are for the positive slope segments and are the result of analyzing the *in silico* dataset at varying data acquisition rates using the Diagnostic Mode of STADIA with constant Minimum Segment Duration of 1.0 seconds and varying Maximum Error Tolerance (row headings) and the Data Acquisition Time Step (column headings). Each gap statistic plot is labeled with the optimal *k*-value suggested (**red** is used to indicate agreement with the results for this dataset in the main text; **gray** indicates parameter combinations resulting in different *k*-values.). Plots with * lack a clear local maximum and are therefore inconclusive.). The **dark blue** box indicates the parameter space for which cluster profiles and labeled length-history plots can be found in **Figure S3.6. (B)** A representative gap statistic plot (bottom right) shows the axes for each plot in (A). The x-axis (*k*-value) range is the same for all plots. The y-axis (Gap-Value) has differing ranges (not shown) for each plot, but the specific numerical values of the gap statistic are not relevant to interpreting the plots because identification of the optimal *k*-value is based on local maxima within each gap statistic plot. In other words, the pertinent information is the relationship between the values of the gap statistic at different *k*-values within each plot, not the values themselves. † The **full resolution** *in silico* data have temporal resolution of one output per dimer-scale biochemical event (see Methods).

**Figure S3.5 – Observations and Interpretations**

**Observations:** When using a Minimum Segment Duration of 1.0 seconds, the results of analyzing the negative slope line segment data from piecewise linear approximation of the *in silico* data captured at 0.1, 0.5, and 1.0 second Data Acquisition Time Steps are very similar to the results from analysis of the full resolution data. The general pattern of a peak at *k*=3 is consistent across almost all the tested combinations of Data Acquisition Time Step and Maximum Error Tolerance.

**Interpretations:** The stability of the *k*-value suggested by the gap statistic plots across the Data Acquisition Time Steps is an encouraging result for the conclusion that stutters exist and are not simply a result of analyzing data at high temporal resolution. The sensitivity to changes in the user-input parameter Maximum Error Tolerance is also generally similar across the Data Acquisition Time Steps. From the parameter sweep results for the full resolution data in **Supplemental Section 2**, however, we know that even when stutters are detected as a distinct cluster, the detection of individual stutter segments (particularly those preceding catastrophe) decreases when using Minimum Segment Durations of 1.0 seconds or more (**Figures S2.3, S2.4, S2.9**). This observation is not surprising given that some events that would be interpreted by eye as stutters last less than one second.

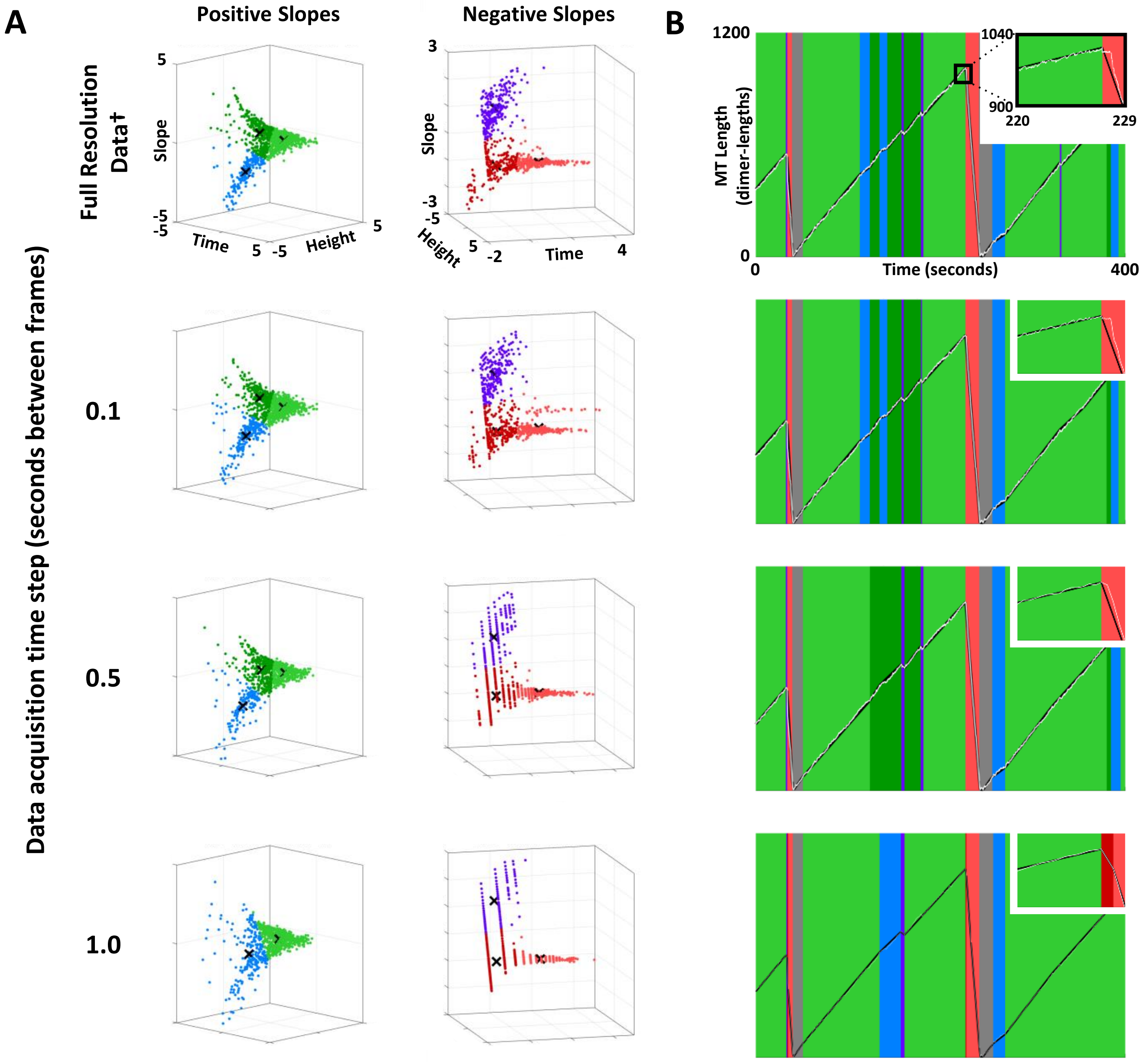


**Figure S3.6. Cluster profiles of positive and negative slope segment data (A) with representative labeled length-history plots (B).** The cluster plots and labeled length-history plots represent classification results from using STADIA in Automated Mode with the *k*-values indicated by the corresponding gap statistic plots in **Figure S3.4** and **S3.5**. Data acquisition rates are indicated by row headers. The Minimum Segment Duration was set to 1.0 seconds and the Maximum Error Tolerance was set to 20 dimer-lengths. † The **full resolution** *in silico* data have temporal resolution of one output per dimer-scale biochemical event (see Methods).

**<u>Figure S3.6 A – Observations and Interpretations</u>**

**Observations:** The cluster profiles for the positive and negative slope segment data maintain the same general shape regardless of the Data Acquisition Time Step when the data are analyzed using a Minimum Segment Duration of 1.0 seconds and Maximum Error Tolerance of 20 subunits. A notable difference across the varying Data Acquisition Time Steps, however, is that the overall density of data points decreases with increasing Data Acquisition Time Steps (i.e., moving down the rows).

**Interpretations:** The relative stability of the shape of the cluster profiles for the positive and negative slope segments with varying Data Acquisition Time Steps bolsters the conclusions about the number of clusters drawn from the gap statistic plots in **Figures S3.4** and **S3.5** (recall that the gap statistic drives the decision for the optimal *k*-value, but cluster profiles are also used to inform the *k*-value).

**Additional Discussion:** The appearance of a loss of density (particularly in the shortening segments) is due, in part, to the data points collapsing onto lines that correspond to multiples of the Data Acquisition Time Step. Additionally, with the longer Data Acquisition Time Steps, more loss of density occurs in the stutters and brief growth and shortening than in the sustained growth and shortening (there are not as many short segments).

Lastly, similar to the analysis of the full resolution data with varying Minimum Segment Durations (**Figure S2.3**), changing the data acquisition rate can affect the shapes and centroid locations of the clusters even if the number of clusters remains the same (e.g., compare the positive slope segment clusters for the full resolution data to the positive slope segment clusters for Data Acquisition Time Step = 0.5 seconds).

**<u>Figure S3.6 B – Observations and Interpretations</u>**

**Observations:** The labeled length-history data illustrate that detection of stutters is reliant on both the Data Acquisition Time Step and the Minimum Segment Duration. In contrast to the analysis conducted at Minimum Segment Duration = 0.5 seconds (**Figure S3.3**), STADIA is not capable of identifying the catastrophe in the zoomed-in portraits in (B) as transitional. Recall that by eye, the catastrophe is clearly transitional in the full resolution data. The catastrophe as seen in the length-history data itself (white line) appears to become less transitional as the Data Acquisition Time Step increases, until the catastrophe appears to be undeniably abrupt at Data Acquisition Time Step = 1.0 seconds. Additionally, for increasing values of the Data Acquisition Time Step, the number of stutters are detected throughout the plotted region of length-history data generally decreases.

**Interpretations:** This example demonstrates that detection of stutters is reliant on both the Minimum Segment Duration and the Data Acquisition Time Step. Note that the real constraint being applied in this analysis is the Data Acquisition Time Step because that limits the user-input Minimum Segment Duration to choices greater than or equal to the largest Data Acquisition Time Step considered in this figure (i.e., 1.0 seconds). Since the parameter sweep in **Supplemental Section 2** demonstrated decreased detection of stutters for Minimum Segment Durations > 0.5 seconds, we conclude that data acquired at rates slower than 2 fps are not ideal for detection of stutters.

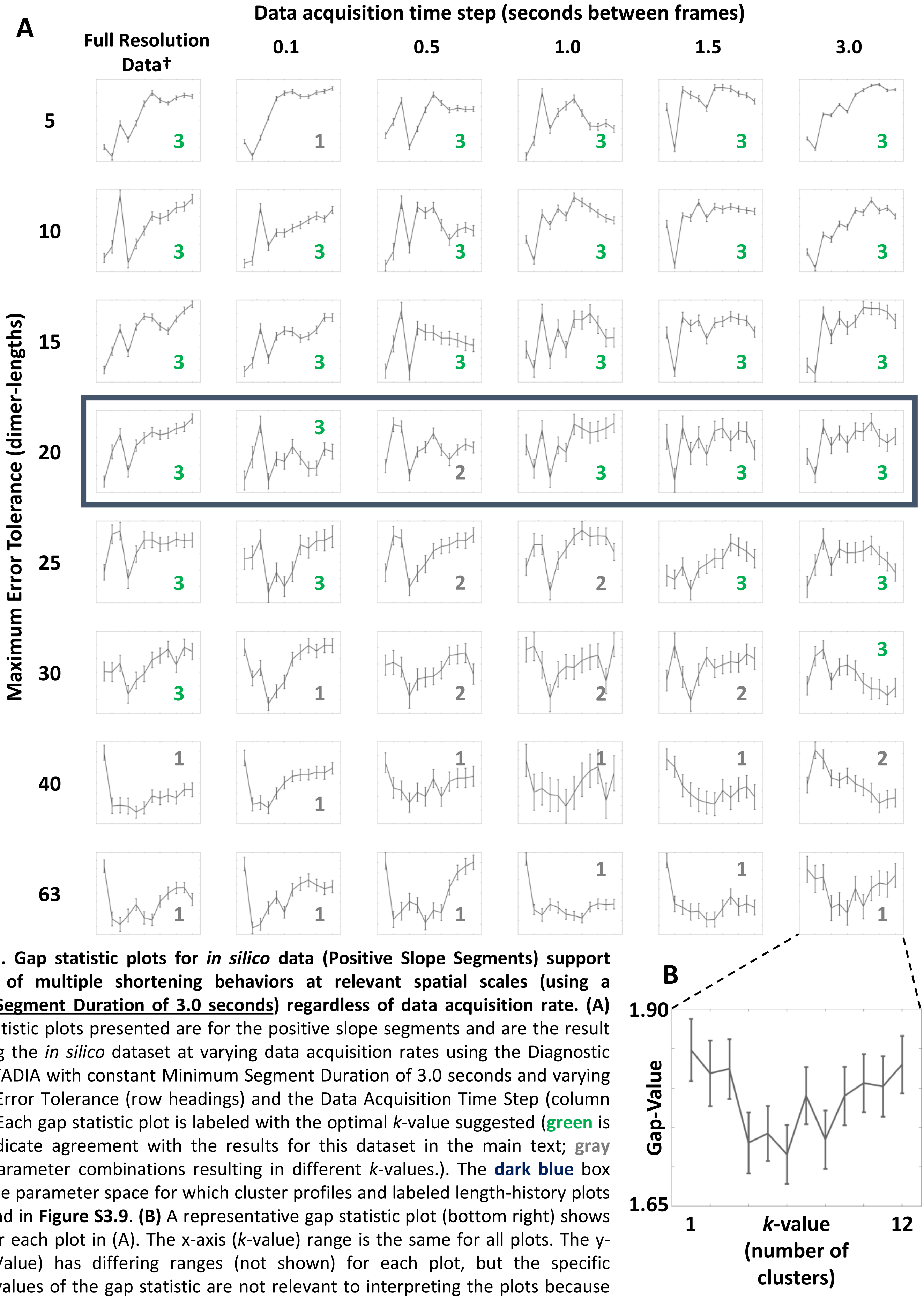


**Figure S3.7. Gap statistic plots for *in silico* data (Positive Slope Segments) support conclusion of multiple shortening behaviors at relevant spatial scales (using a <u>Minimum Segment Duration of 3.0 seconds</u>) regardless of data acquisition rate. (A)** The gap statistic plots presented are for the positive slope segments and are the result of analyzing the *in silico* dataset at varying data acquisition rates using the Diagnostic Mode of STADIA with constant Minimum Segment Duration of 3.0 seconds and varying Maximum Error Tolerance (row headings) and the Data Acquisition Time Step (column headings). Each gap statistic plot is labeled with the optimal *k*-value suggested (green is used to indicate agreement with the results for this dataset in the main text; gray indicates parameter combinations resulting in different *k*-values.). The **dark blue** box indicates the parameter space for which cluster profiles and labeled length-history plots can be found in **Figure S3.9**. **(B)** A representative gap statistic plot (bottom right) shows the axes for each plot in (A). The x-axis (*k*-value) range is the same for all plots. The y-axis (Gap-Value) has differing ranges (not shown) for each plot, but the specific numerical values of the gap statistic are not relevant to interpreting the plots because identification of the optimal *k*-value is based on local maxima within each gap statistic plot. In other words, the pertinent information is the relationship between the values of the gap statistic at different *k*-values within each plot, not the values themselves. † The **full resolution** *in silico* data have temporal resolution of one output per dimer-scale biochemical event (see Methods).

**<u>Figure S3.7 – Observations and Interpretations</u>**

**Observations:** When using a Minimum Segment Duration of 3.0 seconds, the results of analyzing the positive slope line segment data from piecewise linear approximation of the *in silico* data captured at 0.1, 0.5, 1.0, 1.5, and 3.0 second Data Acquisition Time Steps are similar to, albeit noisier than, the results from analysis of the full resolution data. The general pattern of *k*=3 at sufficiently small Maximum Error Tolerances and a diminishing number of clusters with increasing Maximum Error Tolerance is upheld across the Data Acquisition Time Steps considered. The noise (as indicated by the size of the error bars) seems to increase with larger Data Acquisition Time Steps when analysis is conducted using a Minimum Segment Duration of 3.0 seconds.

**Interpretations:** The stability of the *k*-value suggested by the gap statistic plots across the Data Acquisition Time Steps is an encouraging result for the conclusion that stutters exist and are not simply a result of analyzing data at fine resolution. The sensitivity to changes in the user-input parameter Maximum Error Tolerance is also generally similar across the Data Acquisition Time Steps. From the parameter sweep results for the full resolution data in **Supplemental Section 2**, however, we know that even when stutters are detected as a distinct cluster, individual stutter segments of all kinds start to be missed when using a Minimum Segment Duration of 3.0 seconds (e.g., **Figure S2.9**).

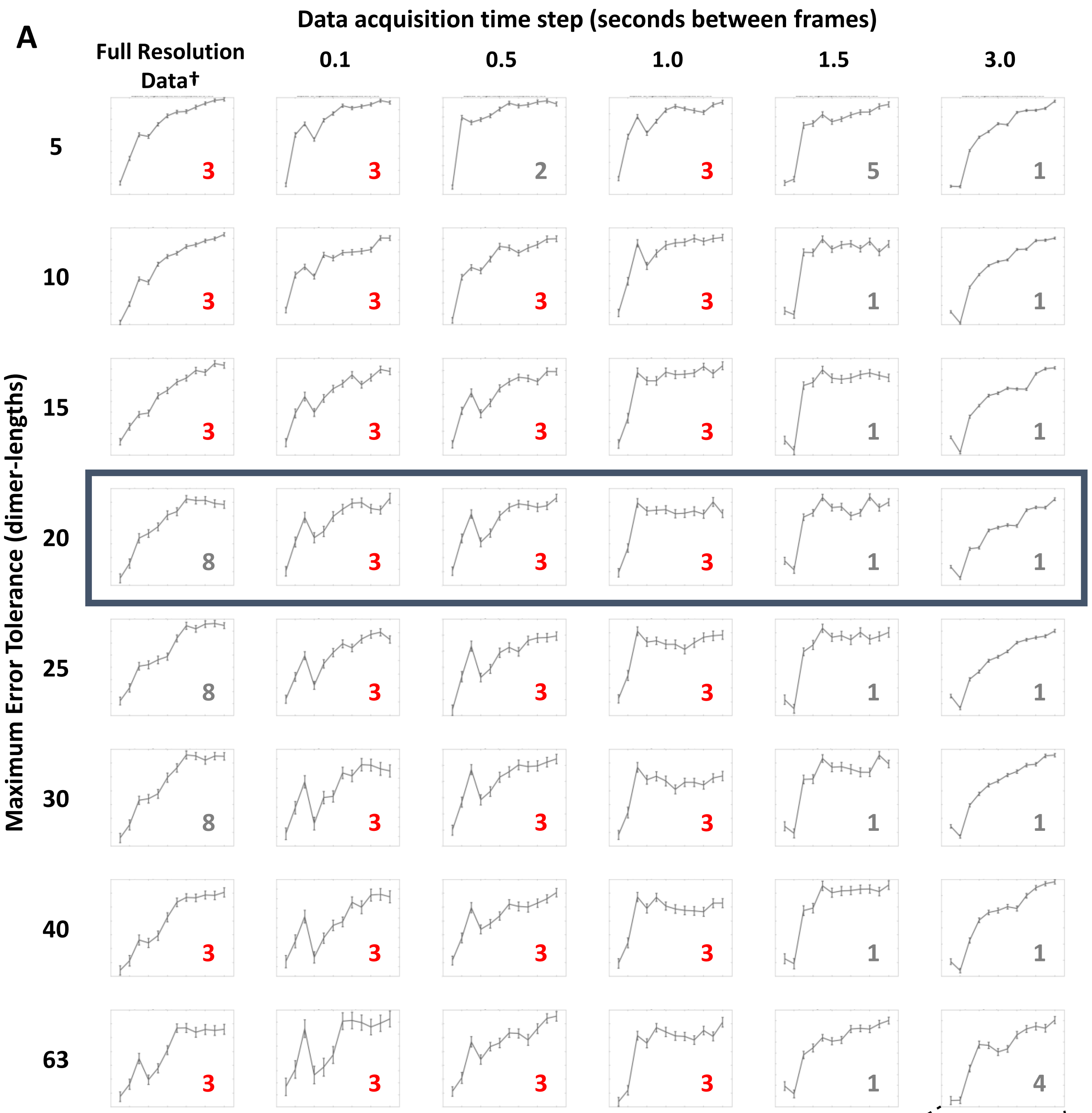


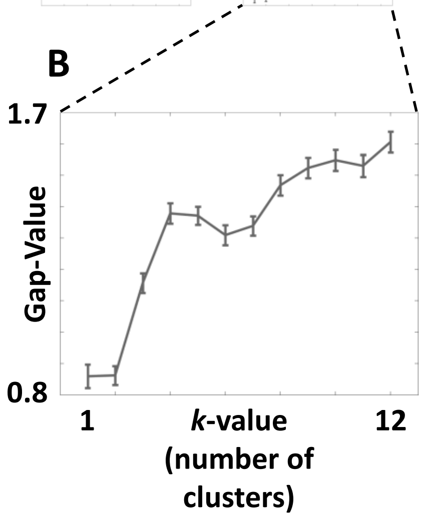


**Figure S3.8. Gap statistic plots for *in silico* data (Negative Slope Segments) support conclusion of multiple shortening behaviors at relevant spatial scales (using a <u>Minimum Segment Duration of 3.0 seconds</u>) regardless of data acquisition rate. (A)** The gap statistic plots presented are for the positive slope segments and are the result of analyzing the *in silico* dataset at varying data acquisition rates using the Diagnostic Mode of STADIA with constant Minimum Segment Duration of 3.0 seconds and varying Maximum Error Tolerance (row headings) and the Data Acquisition Time Step (column headings). Each gap statistic plot is labeled with the optimal *k*-value suggested (red is used to indicate agreement with the results for this dataset in the main text; gray indicates parameter combinations resulting in different *k*-values.). The dark blue box indicates the parameter space for which cluster profiles and labeled length-history plots can be found in **Figure S3.9. (B)** A representative gap statistic plot (bottom right) shows the axes for each plot in (A). The x-axis (*k*-value) range is the same for all plots. The y-axis (Gap-Value) has differing ranges (not shown) for each plot, but the specific numerical values of the gap statistic are not relevant to interpreting the plots because identification of the optimal *k*-value is based on local maxima within each gap statistic plot. In other words, the pertinent information is the relationship between the values of the gap statistic at different *k*-values within each plot, not the values themselves. **†** The **full resolution** *in silico* data have temporal resolution of one output per dimer-scale biochemical event (see Methods).

**Figure S3.8 – Observations and Interpretations**

**Observations:** When using a Minimum Segment Duration of 3.0 seconds, the results from analyzing the negative slope line segment data from piecewise linear approximation of the *in silico* data captured at 0.1, 0.5, and 1.0 second Data Acquisition Time Steps are similar to each other, and somewhat similar to the results from analysis of the full resolution data depending on the Maximum Error Tolerance. Clear differences occur in two places: 1) as was seen in **Supplemental Figure S2.2**, the gap statistic plots for the full resolution data are generally increasing when analyzed using a Minimum Segment Duration of 3.0 seconds, while the gap statistic plots corresponding to Data Acquisition Time Steps of 0.1, 0.5, and 1.0 seconds clearly suggest $k$=3 as the optimal number of clusters; and 2) analysis of data acquired at very slow acquisition rates (i.e., Data Acquisition Time Steps of 1.5 and 3.0 seconds) usually results in $k$=1 suggested by the gap statistic.

**Interpretations:** The stability of the $k$-value suggested by the gap statistic plots across most of the Data Acquisition Time Steps (i.e., 0.1, 0.5, and 1.0 seconds) is an encouraging result for the conclusion that stutters exist and are not simply a result of analyzing data at fine resolution. For the Data Acquisition Time Steps of 0.1 seconds or more, there is little sensitivity to changes in the user-input parameter Maximum Error Tolerance. The loss of detection of stutters at the very slow data acquisition rates (i.e., 1.5 and 3.0 seconds) is not surprising. As was noted in the parameter sweep with the full resolution data in **Supplemental Section 2**, MT depolymerizations occur at relatively short time scales (often < 3 seconds); therefore, reliably approximating depolymerizations can become challenging, let alone detecting multiple behaviors within the negative slope segments, when data is acquired with time steps this long.

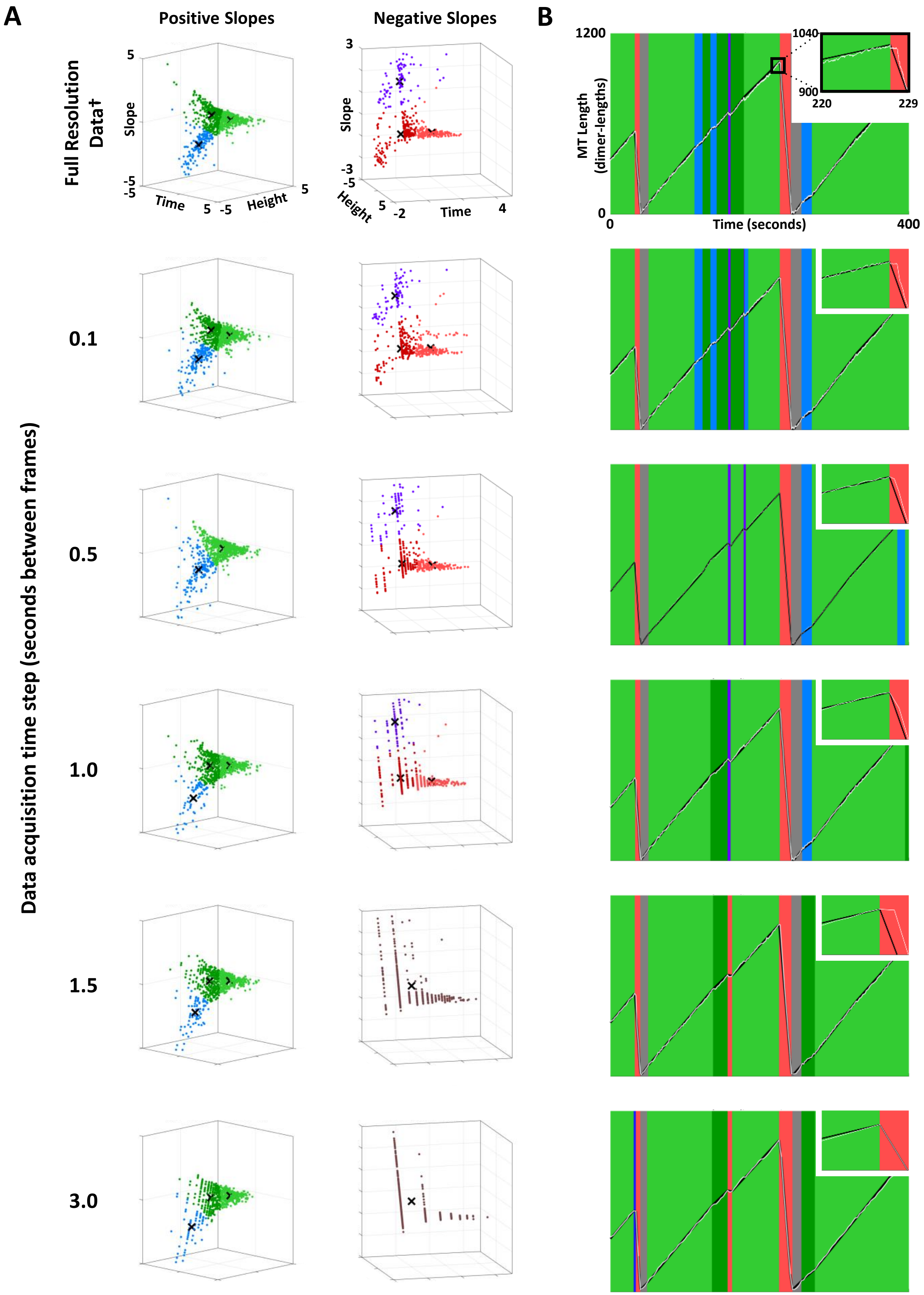


**Figure S3.9. Cluster profiles of positive and negative slope segment data (A) with representative labeled length-history plots (B).** The cluster plots and labeled length-history plots represent classification results from using STADIA in Automated Mode with the *k*-values indicated by the corresponding gap statistic plots in **Figure S3.7** and **S3.8**. Data acquisition rates are indicated by row headers. The <u>Minimum Segment Duration was set to 3.0 seconds</u> and the Maximum Error Tolerance was set to 20 dimer-lengths. **†** The **full resolution** *in silico* data have temporal resolution of one output per dimer-scale biochemical event (see Methods).

**Figure S3.9 A – Observations and Interpretations**

**Observations:** The cluster profiles for the positive slope data maintain the same general shape regardless of the Data Acquisition Time Step when the data are analyzed using a Minimum Segment Duration of 3.0 seconds and Maximum Error Tolerance of 20 subunits. The negative slope segment data have similar shapes as well, however the number of (distinguishable) data points at longer Data Acquisition Time Steps is so minimal that without the profiles from smaller Data Acquisition Time Steps, it would be difficult to confidently say what the shape of the profile is. It is also notable that datasets that are ~~that~~ this sparse would generally not be considered good candidates for *k*-means clustering.

**Interpretations:** The relative stability of the shape of the cluster profiles for positive slope segments with varying Data Acquisition Time Steps bolsters the conclusions about the number of clusters drawn from the gap statistic plots in **Figures S3.7** and **S3.8** (recall that the gap statistic drives the decision for the optimal *k*-value, but cluster profiles are also used to inform the *k*-value).

**Additional Discussion:** The appearance of a loss of density (particularly in the shortening segments) is due, in part, to the data points collapsing onto lines that correspond to multiples of the Data Acquisition Time Step. Additionally, with the longer Data Acquisition Time Steps, more loss of density occurs in the stutters and brief growth and shortening than in the sustained growth and shortening (there are not as many short segments). Particularly for the negative slope segment data, it is evident that the datasets at large Data Acquisition Time Steps (i.e., 1.5 and 3.0 seconds) are too sparsely filled to draw conclusion about the number of behaviors.

Lastly, similar to the analysis of the full resolution data with varying Minimum Segment Durations (**Figure S2.3**), changing the data acquisition rate can affect the shapes and centroid locations of the clusters even if the number of clusters remains the same (e.g., compare the positive slope segment clusters for the full resolution data to the positive slope segment clusters for Data Acquisition Time Step = 3.0 seconds). Interestingly, the shapes of the positive slope segment clusters for the higher Data Acquisition Time Steps appear more similar to the results from the full resolution mean PF data than the full resolution max PF data (**Figure S1.4**).

**Figure S3.9 B – Observations and Interpretations**

**Observations:** The labeled length-history data illustrate that detection of stutters is reliant on both the Data Acquisition Time Step as well as the Minimum Segment Duration. In contrast to the analysis conducted at Minimum Segment Duration = 0.5 seconds (**Figure S3.3**), STADIA is not capable of identifying the catastrophe in the zoomed-in portraits in (B) as transitional. Recall that by eye, the catastrophe was clearly transitional in the full resolution data. The catastrophe as seen in the length-history data itself (white line) appears to become less transitional as the Data Acquisition Time Step increases until the catastrophe appears to be undeniably abrupt at Data Acquisition Time Step = 1.0 seconds. When the Data Acquisition Time Step is 1.5 seconds, the location of the catastrophe changes and, by chance, the captured data points result in what appears to be a transitional catastrophe again but is not detected as transitional using a Minimum Segment Duration of 3.0 seconds. When the data acquisition rate is increased to 3.0 seconds, the catastrophe is completely abrupt in appearance. Additionally, for increasing values of the Data Acquisition Time Step, the number of stutters are detected throughout the plotted region of length-history data generally decreases.

**Interpretations:** This example demonstrates that detection of stutters is reliant on both the Minimum Segment Duration and the Data Acquisition Time Step. Note that the real constraint being applied in this analysis is the Data Acquisition Time Step because that limits the user-input parameter, Minimum Segment Duration, to choices greater than or equal to the largest Data Acquisition Time Step considered in this figure (i.e., 3.0 seconds). Since the parameter sweep in **Supplemental Section 2** demonstrated decreased detection of stutters for Minimum Segment Durations > 0.5 seconds, we conclude that data acquired at rates slower than 2 fps are not ideal for detection of stutters.

#### Summary of Conclusions: Section 3, Data acquisition rate sensitivity analysis

The analyses in this section aid in answering the following question: do the main conclusions of this manuscript change when the temporal resolution of inputted length-history data is varied?

The results of these analyses indicate that the main conclusions of this manuscript are sound regardless of temporal resolution so long as the Data Acquisition Time Step and Minimum Segment Duration are sufficiently small. Based on the loss of density in the cluster profiles at slower data acquisition rates, we recommend using Data Acquisition Time Steps less than or equal to 0.5 seconds. This recommendation is also consistent with the Minimum Segment Duration results in **Supplemental Section 2**.

The results of these analyses demonstrate that the detection of three clusters is upheld in the positive slope segment data for Data Acquisition Time Steps up to 3 seconds, and in the negative slope segment data for Data Acquisition Time Steps up to 1 second, assuming reasonable choices for Maximum Error Tolerance and Minimum Segment Duration. However, even when three clusters are detected, fewer of the shortest duration segments in the clusters of up/down stutter and brief growth/shortening are detected as the Data Acquisition Time Step and/or the Minimum Segment Duration increase. For example, detection of the particular stutter shown in the zoomed-in length-history plots is lost at Minimum Segment Durations of 1 second or greater.

### **Supplemental Section 4:**
Negative Control: Simulations of a Two-State (Growth-Shortening) Model

**Table of Contents**


**1. Text**
- **a.** ***Summary*** – page 59
- **b.** ***Two-state simulation method*** – page 60
- **c.** ***Results of STADIA analysis of the two-state simulation data*** – page 61

**2. Figures**
- **a.** Analysis of **Positive** Slope Segments – **Figure S4.1**, page 62
  *Gap statistic plots, cluster profiles, and segment features box plots*
- **b.** Analysis of **Negative** Slope Segments – **Figure S4.2**, page 63
  *Gap statistic plots, cluster profiles, and segment features box plots*
- **c.** Labeled length-history plots – **Figure S4.3**, page 64

### Supplemental Section 4:
#### Negative Control: Simulations of a Two-State (Growth-Shortening) Model

***Summary***

As a negative control, we ran STADIA on length-history simulation data from a model designed to have only two states: growth and shortening. In this two-state model, the traditional DI parameters ($V_{growth}$, $V_{short}$, $F_{cat}$, and $F_{res}$) are inputs into the model.

In the STADIA analysis of the two-state simulation data, we varied the values of the Minimum Segment Duration and the Maximum Error Tolerance, as well as the time between data points in the length-history data inputted into STADIA (see **Supplemental Sections 2 & 3** for results of varying these parameter values in the STADIA analysis of the datasets used in the main text). As discussed in the main text, the values of the Minimum Segment Duration and the Maximum Error Tolerance determine how closely the piecewise linear approximation produced by the segmentation stage of STADIA matches the length-history data inputted into STADIA.

For the STADIA analysis of the two-state simulation data, **Supplemental Figures S4.1** and **S4.2** show the resulting gap statistic plots (used to inform the number of clusters), cluster plots in the log-transformed and standardized feature space, and box plots of the segment features (time duration, height change, and slope) for each cluster. The clusters identified in **Supplemental Figures S4.1** and **S4.2** are applied to produce the labeled length-history plots in **Supplemental Figure S4.3.**

The results show that STADIA did *not* identify clusters of up stutters or down stutters when analyzing the positive slope segments or the negative slope segments, respectively (**Supplemental Figures S4.1, S4.2**). For data with relatively large time steps between data points (0.5 seconds or more), STADIA did identify a small number of flat stutters in rare cases where a time step between two consecutive data points happened to straddle a catastrophe or rescue in such a way as to produce a near-flat segment (one example in **Supplemental Figure S4.3, row D, right panel, orange bracket**). These flat stutters were very short in duration and totaled to less than 0.09% of the total simulation time; this percentage is approximately two orders of magnitude smaller than the percent time spent in stutters in the detailed dimer-scale 13-PF simulation data (see **Supplemental Figures S1.9, S1.10**). Below we provide more details about the two-state simulation method and the STADIA analysis of the two-state simulation data.

***Two-state simulation method***

In the two-state simulations, each microtubule grows from a stable seed and is assumed to be in either a growth state or a shortening state at each point in time. The traditional DI parameters are input parameters in the two-state model; this is in contrast to the dimer-scale 13-PF model, where the DI parameters are emergent properties resulting from dimer-scale biochemical events and are measured from the length-history data outputted by the simulation.

To generate the two-state length-history data, the DI parameter values measured from the Peak-Valley analysis of the dimer-scale 13-PF simulation data (first row of **Table 1**) were used as the values of the inputted DI parameters in the two-state simulations. The time duration of each growth segment (i.e., the time from the start of a growth segment until a catastrophe) was sampled from an exponential distribution with rate $F_{cat}$ = 0.659 $min^{-1}$. The slope or growth velocity ($V_{growth}$) of each growth segment was sampled from a normal distribution with mean of $V_{growth}$ = 46.1 nm/s and standard deviation of $V_{growth}$ = 5.1 nm/s. The length change of each growth segment was calculated as the sampled time duration multiplied by the sampled slope.

Similarly, the time from the beginning of a shortening segment until a potential rescue was sampled from an exponential distribution with rate $F_{res}$ = 2.483 $min^{-1}$. The slope or shortening velocity ($V_{short}$) of each shortening segment was sampled from a normal distribution with mean of $V_{short}$ = 540.0 nm/s and standard deviation of $V_{short}$ = 47.9 nm/s. The length change of each shortening segment was calculated as the sampled time duration multiplied by the sampled slope. If this calculated length change would result in the MT length becoming negative, then the shortening segment was terminated when the MT length reached zero. In this case, the actual time duration of the segment was less than, not equal to, the sampled time until a potential rescue; then, the actual time duration was calculated from the sampled slope and the length change from the start of the shortening segment until the MT length reached zero.

One MT was simulated for 10 hours of simulation time, chosen to produce a comparable amount of data as the dimer-scale 13-PF simulations. To generate the length-history data to input into STADIA, the raw length-history data from the two-state simulations was interpolated to have a fixed data acquisition rate (“frame” rate) of 20 fps, 2 fps, or 1/3 fps. As in Section 3, ‘data acquisition rate in frames per second’ = 1/‘Data Acquisition Time Step in seconds’.

***Results of STADIA analysis of the two-state simulation data***

The results of the STADIA analysis of the two-state simulation data are shown in **Supplemental Figures S4.1-S4.3**. In these figures, the values of the data acquisition rate, the Minimum Segment Duration, and the Maximum Error Tolerance used in the STADIA analysis are indicated above each row. The clustering results for the positive and negative slope segments are provided in **Supplemental Figures S4.1** and **S4.2** respectively, and portions of the resulting labeled length-history plots are shown in **Supplemental Figure S4.3**.

Depending on the values of the data acquisition rate, Minimum Segment Duration, and Maximum Error Tolerance, the gap statistic plots indicate a *k*-value (number of clusters) equal to either 1 or 2, or an increasing gap statistic plot with no clear local maximum (gap plots in first column of **Supplemental Figures S4.1, S4.2**). This result differs from the *k*-value of 3 identified in the dimer-scale 13-PF simulation data and the *in vitro* data (as discussed in the main text, the negative slope segments from the *in vitro* data yielded *k*=2, with the two clusters corresponding to down stutters and brief shortening; this occurred because shortening phases were not captured in their entirety in the *in vitro* data, so the sustained shortening cluster was missing).

Furthermore, the cluster plots for the two-state data (second column **Supplemental Figures S4.1, S4.2**) show a cloud of points that is oblong but has no distinct appendages, in contrast to the 3 appendages visible in the cluster plots from the dimer-scale 13-PF simulation data and the positive slope segments of the *in vitro* data (**Figure 3 A,B,D,E, Supplemental Figures S1.4, S1.5**).

In cases where the gap statistic indicates *k*=2 for the positive or negative slope segments from the two-state simulation data, the two clusters are essentially indistinguishable in slope; they differ primarily in time duration and correspondingly in height change (see box plots in the third, fourth, and fifth columns of **Supplemental Figures S4.1, S4.2**). In contrast, for the dimer-scale 13-PF simulation data and the *in vitro* data, slope was the primary factor differentiating the clusters of up and down stutters from the clusters of (brief/sustained) growth and shortening (**Figure 3 C,F** and **Supplemental Figure S1.8**).

As discussed in the main text, STADIA identifies near-zero slope segments called flat stutters before performing the clustering step separately on the remaining positive and negative slope segments. In the two-state simulation data at 20 fps (0.05 s between data points), no flat stutters were identified. At 2 fps (0.5 s between data points), STADIA identified 26 flat stutters, of which 25 had a time duration of 0.5 s and one had a time duration of 1.5 s; these time durations total to 14 s, which is approximately 0.039% of the total 10-hour simulation time. At 1/3 fps (3 s between data points), STADIA identified 10 flat stutters, all with a time duration of 3 s, totaling to 30 s or approximately 0.083% of the total time. The segments identified as flat stutters occurred in the rare cases when a time step between two consecutive data points in the data with an imposed fixed frame rate happened to straddle a catastrophe or rescue from the original raw data in a way that produced a near-flat segment (one example in **Supplemental Figure S4.3**, compare right panel of row D to right panels of the other rows). Furthermore, none of the flat stutters detected at 1/3 fps overlapped with any of the flat stutters detected at 2 fps.

Thus, STADIA did not detect robust flat stutters nor any up/down stutters in the two-state simulation data, contrasting the robust detection of stutters in the dimer-scale 13-PF simulation data and the *in vitro* data (see **Supplemental Sections 2 & 3** for parameter sweep analyses testing robustness).

### Negative Control: Two-state (Growth-Shortening) Simulations
### POSITIVE Slope Segments

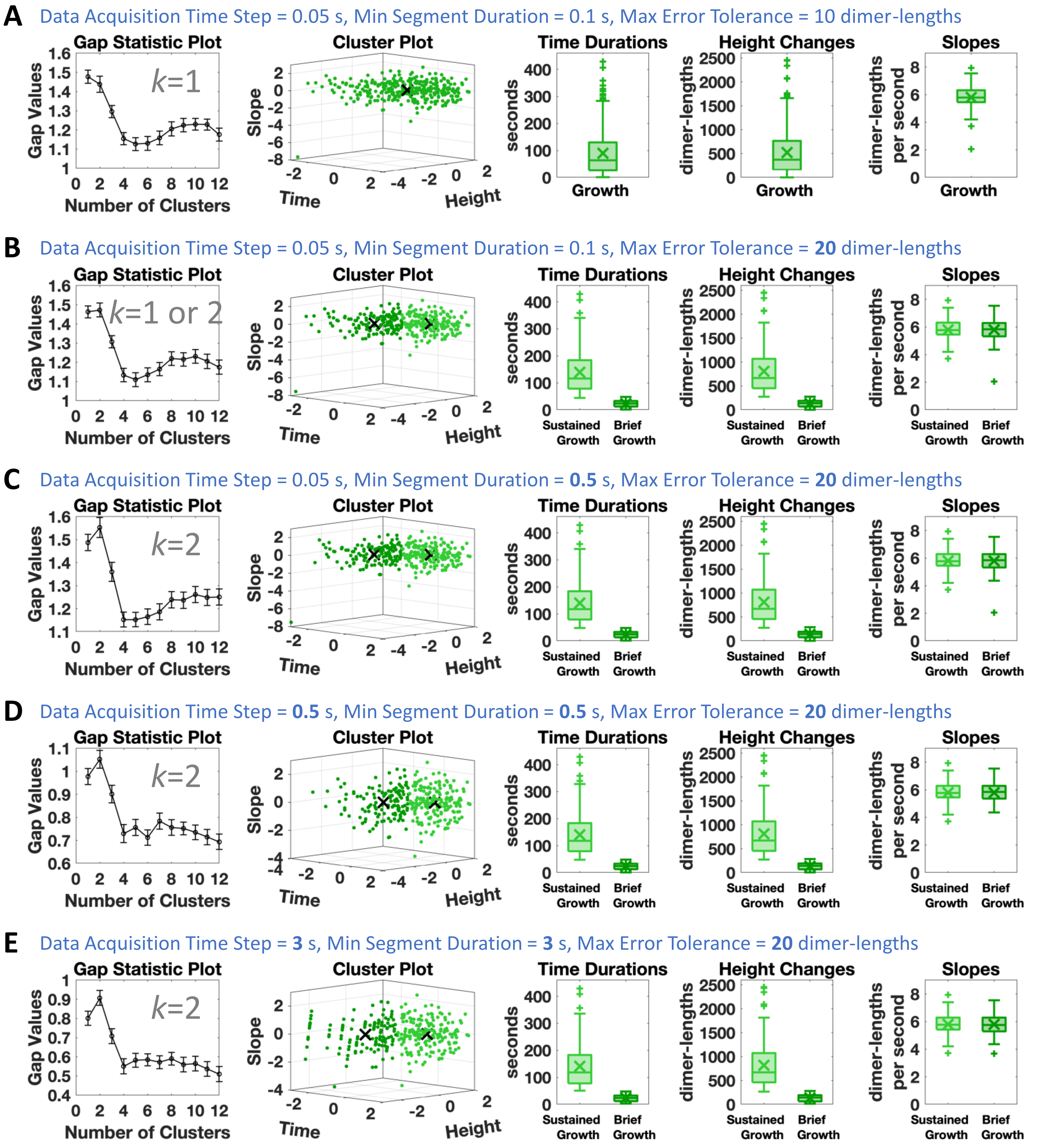


**Figure S4.1. Negative Control: STADIA analysis of *positive* slope segments from a two-state (growth-shortening) model.** From top (row A) to bottom (row E), the piecewise linear approximation to the original raw length-history data becomes coarser as the parameter values in the row headings are varied (values altered from the top row are in bold). In the resulting gap statistic plots, the first local maximum indicates either *k*=1 (row A) or *k*=2 (rows C,D,E). In row B, the first local maximum occurs at *k*=2, but is not conclusively larger than the gap value at *k*=1 when considering the error bars. For the cases where *k*=2, the slopes of the two clusters are indistinguishable. No cluster with shallower slopes (i.e., no cluster of up-stutters) is detected in the positive slope segments. Please see the text on pages 59-61 of this file for additional information and interpretations.

#### Negative Control: Two-state (Growth-Shortening) Simulations
#### NEGATIVE Slope Segments

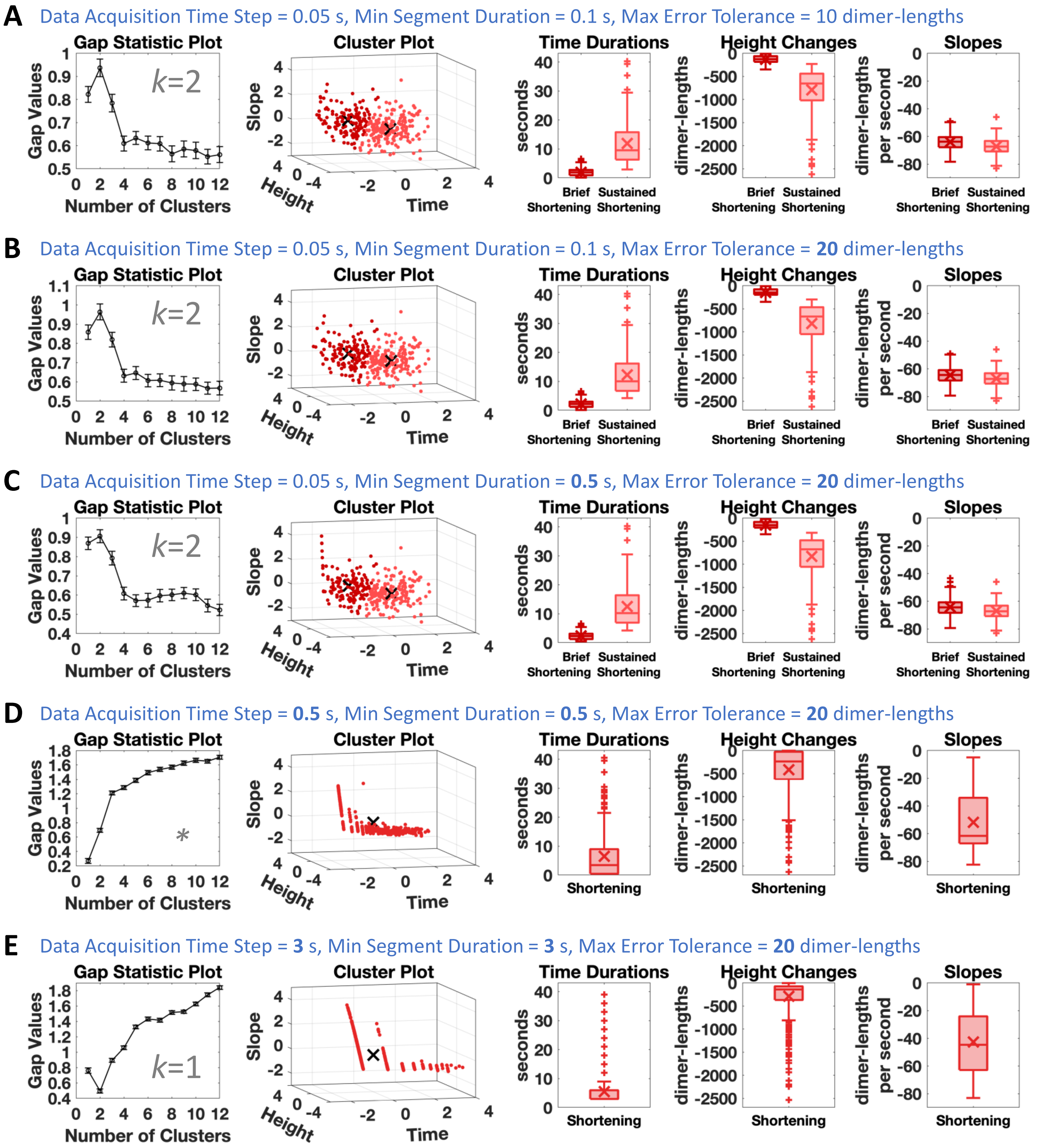


**Figure S4.2. Negative Control: STADIA analysis of *negative* slope segments from a two-state (growth-shortening) model.** The parameter values in the row headings are the same as in **Figure S4.1** (values altered from the top row are in bold). In the resulting gap statistic plots, the first local maximum indicates either *k*=1 (row E) or *k*=2 (rows A,B,C), or the gap statistic plot is generally increasing with no clear local maximum (*, row D). For the cases where *k*=2, the slopes of the two clusters are nearly indistinguishable. No cluster with shallower slopes (i.e., no cluster of down-stutters) is detected in the negative slope segments. Please see the text on pages 59-61 of this file for additional information and interpretations.

### Negative Control: Two-state (Growth-Shortening) Simulations

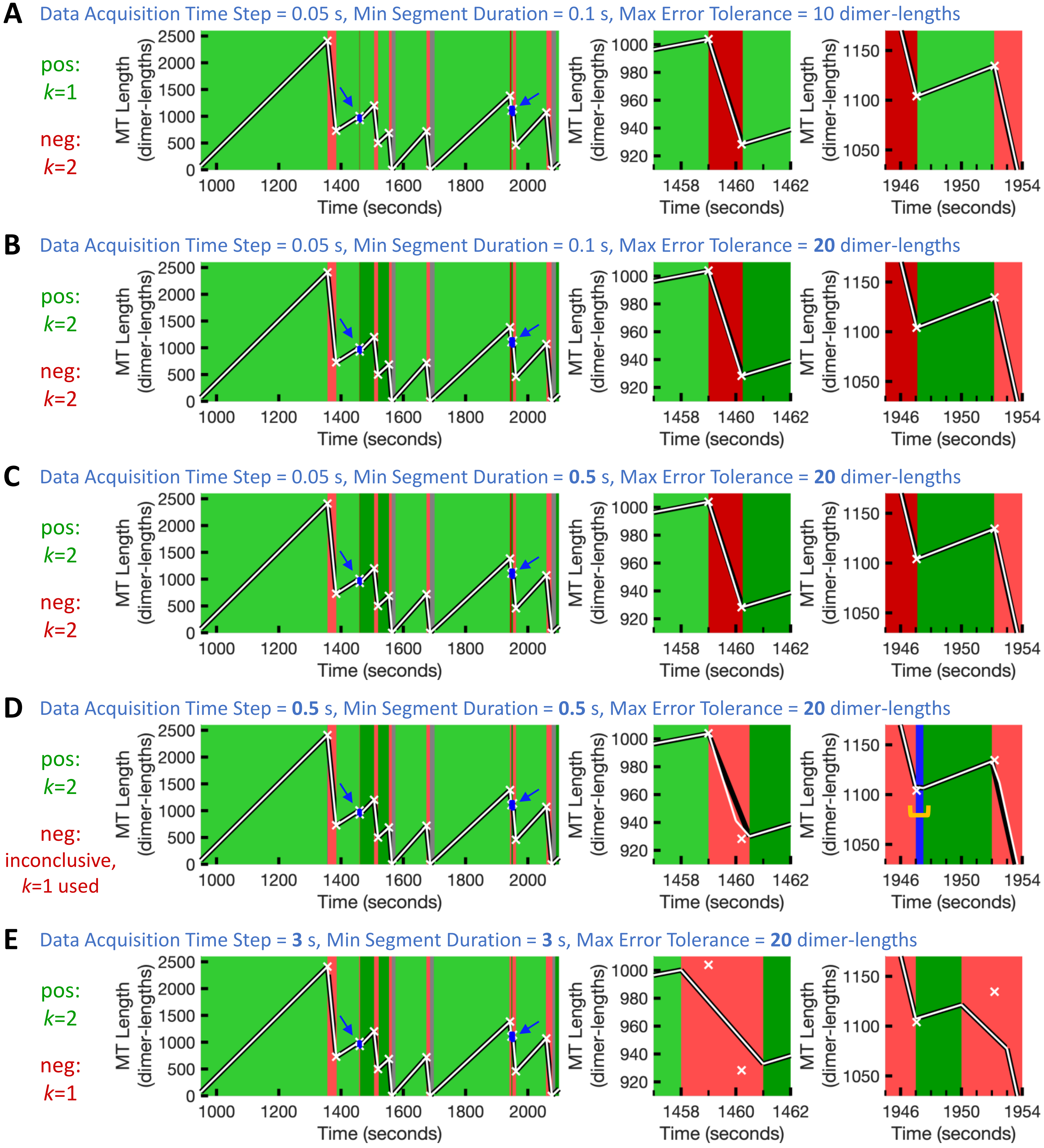


**Figure S4.3. Negative Control: labeled length-history plots from STADIA analysis of two-state (growth-shortening) simulation data.** The parameter values in the row headings are the same as in **Figures S4.1** and **S4.2** (values altered from the top row are in bold). The numbers of clusters (*k*-values) were determined from the gap plots in **Figures S4.1** and **S4.2** The white x-symbols mark the exact points of the transitions (catastrophes, rescues, and complete depolymerizations to the seed) in the raw two-state simulation data (same points in all rows). The white lines show the length-history data inputted into STADIA after the indicated data acquisition rate was imposed. The black lines represent the STADIA piecewise linear approximation. The small blue rectangles (with the blue arrows) in the first column demarcate the regions shown in the zoomed-in plots in the second and third columns. The orange bracket in row D, third column, indicates a rare artifactual detection of a flat stutter. Please see the text on pages 59-61 of this file for additional information and interpretations.
